# Targeting nucleolin facilitates the development of blood-tumor barrier-penetrating and glioblastoma-specific PROTACs

**DOI:** 10.1101/2025.07.22.666084

**Authors:** Hongzhen Chen, Junyi Zhao, Fang Qiu, Shiding Ying, Zhuqian Wang, Aiping Lu, Chao Liang

## Abstract

Targeted therapy for glioblastoma (GBM) is challenged by the blood-tumor barrier (BTB). Extracellular vesicles (EVs) from GBM cells play a role in transforming BBB into BTB, although the mechanisms are not fully understood. This study identifies nucleolin (NCL), a nucleomembrane shuttling protein, as being transferred from GBM cells to the surface of brain capillary endothelial cells, facilitating BTB formation. The aptamer AS1411, which targets NCL, is shown to cross the BTB via receptor-mediated transcytosis (RMT) and selectively recognize GBM cells in an NCL-dependent manner. Beyond its targeting capabilities, AS1411 has been repurposed to recruit the E3 ligase MDM2 in PROTACs, leveraging the intracellular interaction of NCL with MDM2. Utilizing AS1411’s multifunctionality in BTB penetration, GBM cell targeting, and MDM2 recruitment, we conjugated AS1411 to VEGFR2 or EGFR ligands to create PROTAC degraders. These constructs induce NCL- and MDM2-dependent ubiquitination and degradation of VEGFR2 or EGFR, demonstrating significant anti-GBM efficacy both in vitro and in vivo, with no toxicity to normal cells. Overall, NCL from GBM-derived EVs emerges as a crucial mediator of BTB formation and serves as a receptor for AS1411-mediated RMT, paving the way for developing PROTACs that effectively traverse BTB and target GBM.

**GRAPHICAL ABSTRACTS:** 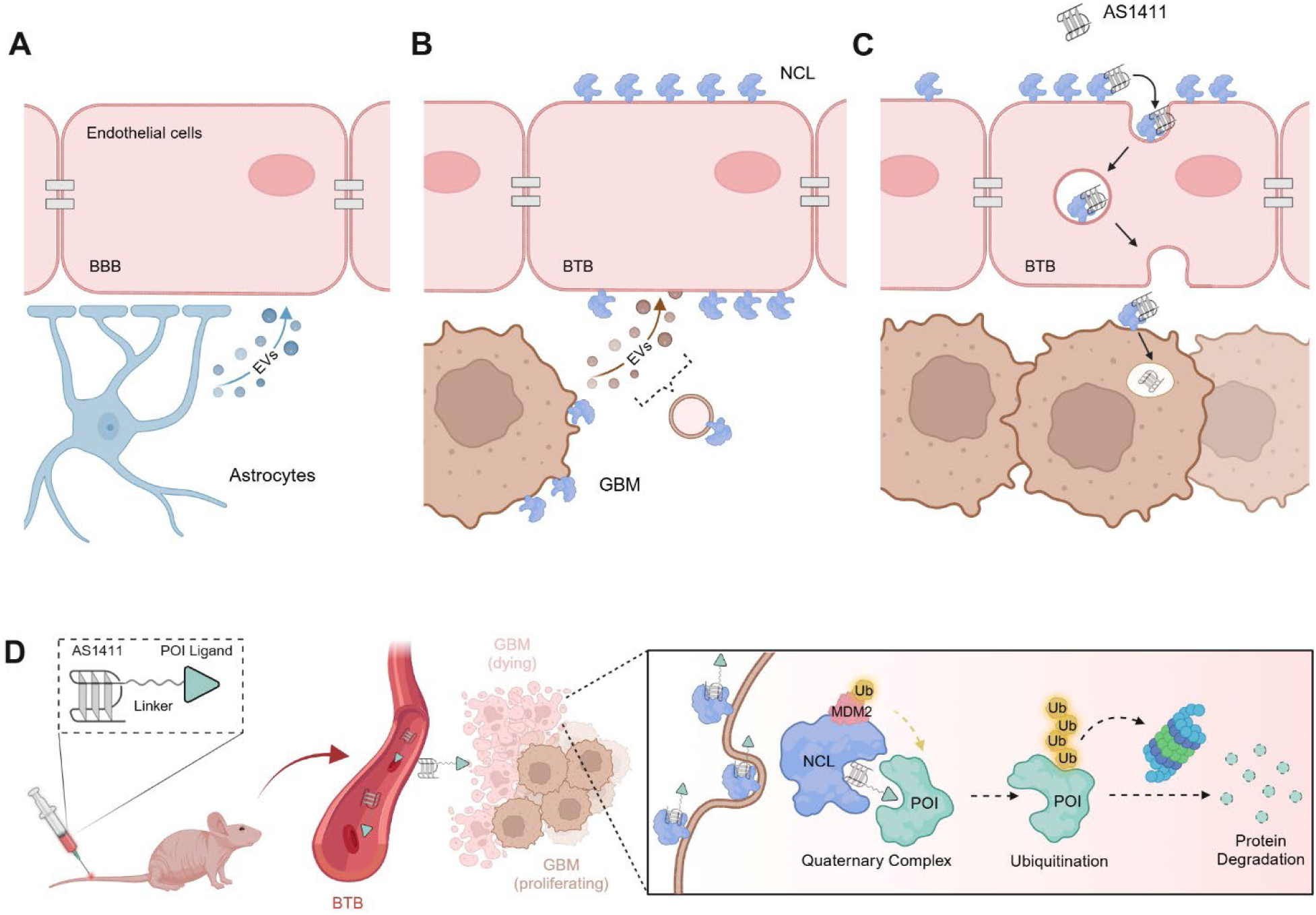

## INTRODUCTION

Glioblastoma (GBM) is one of the most aggressive and lethal brain malignancies, characterized by rapid progression, extensive invasiveness, and a dismal prognosis^1^. Despite advances in standard treatments, including surgery, radiotherapy, and chemotherapy, clinical outcomes for GBM patients remain unacceptably low^2–5^, with a median survival time of only 15 to 18 months and fewer than 5% of patients surviving beyond 5 years post-diagnosis^6^. A major obstacle to effective treatment is the blood-brain barrier (BBB), a highly selective structure comprised of endothelial cells connected by tight junctions that line brain capillaries^7^. The BBB plays a crucial role in maintaining central nervous system homeostasis by facilitating the transport of essential nutrients while preventing most therapeutic agents in the bloodstream from entering the brain^8^.

Recently, it has been recognized that GBM alters the brain microenvironment and reshapes the adjacent BBB, resulting in the formation of a dysfunctional vasculature known as the blood-tumor barrier (BTB)^9–11^. Within the BTB, capillary endothelial cells exhibit distinct phenotypic and functional characteristics compared to those in the healthy BBB^11–13^. However, the precise mechanisms by which GBM induces pathological changes in BTB capillaries remain poorly understood^14^. Extracellular vesicles (EVs), a family of membrane-limited vesicles, have emerged as key mediators of intercellular communication between GBM and capillary endothelial cells via the transfer of biomolecules^15, 16^. Specifically, GBM-derived EVs can travel to the brain capillaries and deliver pro-angiogenic factors to capillary endothelial cells, facilitating the abnormal transformation of the BBB into BTB^17–20^. In contrast, normal astrocyte-derived EVs lack this capability^21, 22^. Thus, a deeper exploration of mechanisms by which GBM-derived EVs contribute to BTB formation is essential, as such insights will not only clarify the differences between the BTB and BBB but also provide a foundation for designing BTB-penetrating and GBM-targeting therapies^11^.

In our study, we revealed that EVs derived from GBM cells carried a significant abundance of nucleolin (NCL), a multifunctional protein that shuttles between the nucleus and cytoplasm and is also partially translocated on the surface of cancer cells, including GBM, but not on normal cells^23–25^. We established an *in vitro* BBB or BTB model using a coculture system, revealing that NCL was transferred via GBM-derived EVs to the surface of cocultured capillary endothelial cells. In contrast, normal astrocyte-derived EVs failed to transport NCL to the surface of endothelial cells in the coculture. Moreover, NCL from GBM-derived EVs actively altered the phenotypes of the cocultured capillary endothelial cells, facilitating the pathological transformation of BBB into BTB. Notably, NCL has been reported to function as an internalization receptor for specific ligands^26, 27^. AS1411, a 26-base G-quadruplex-forming single-stranded DNA (ssDNA) aptamer, specifically targets NCL^28^. We found that NCL on the surface of BTB endothelial cells, transferred from GBM-derived EVs, served as a receptor for receptor-mediated transcytosis (RMT), enabling the selective transcytosis of AS1411 across the BTB. Once across the BTB, AS1411 selectively recognized and entered GBM cells.

Proteolysis-targeting chimeras (PROTACs) are emerging therapeutic entities designed to degrade proteins of interest (POIs)^29^. These molecules are heterobifunctional, consisting of an E3 ligase recruiter and a POI-targeting ligand, connected by a linker^30^. This design enables the ubiquitination and subsequent degradation of the POIs^31, 32^. Murine double minute 2 (MDM2) stands out among various E3 ligases due to its prevalent overexpression in multiple cancers, including GBM^33^, making it an attractive E3 for the design of antitumor PROTACs^34^. Intracellularly, NCL interacts with MDM2, not as a substrate for ubiquitination but as an auxiliary partner^35^. We recently discovered that AS1411, upon entering cancer cells, could recruit MDM2 by employing NCL as a molecular bridge^36^. This property allows AS1411 to be incorporated into PROTACs, facilitating the selective degradation of POIs in cancer cells^36^.

Collectively, given the multifunctionality of AS1411 in penetrating the BTB, targeting GBM cells, and recruiting MDM2, we hypothesized that conjugating AS1411 with ligands for POIs in GBM could facilitate the development of PROTACs capable of crossing BTB and targeting GBM. Vascular endothelial growth factor receptor 2 (VEGFR2) and epidermal growth factor receptor (EGFR) are promising therapeutic POIs that are overexpressed in GBM^37–39^. To test our hypothesis, we conjugated AS1411 with a VEGFR2 inhibitor, Cediranib (Ced)^40^, or an EGFR inhibitor, Gefitinib (Gef)^41^, to create PROTACs. Our results showed that these PROTACs effectively brought the NCL-MDM2 complex into spatial proximity with VEGFR2 or EGFR, leading to their ubiquitination and proteasomal degradation in both NCL- and MDM2-dependent manners. The AS1411-based PROTACs exhibited potent anti-tumor effects *in vitro* and *in vivo*, including inhibition of GBM growth, induction of apoptosis, and extension of survival in orthotopic GBM mouse models, all while showing no significant toxicity to normal cells or tissues.

In summary, our findings indicate that GBM cells can transfer NCL to the surface of capillary endothelial cells via EVs, driving the transformation of BBB into BTB. The transferred NCL functions as an RMT receptor on endothelial cells, mediating the specific penetration of AS1411 across the BTB. Building on this newly identified BTB-penetrating ability of AS1411, along with its previously established GBM-targeting and MDM2- recruiting properties, we illustrate that PROTACs integrating AS1411 offer a promising avenue for targeted GBM therapy.

## RESULTS

### GBM-derived EVs transfer NCL to the surface of capillary endothelial cells, facilitating BTB formation

To investigate the difference in protein cargo between GBM-derived EVs and astrocyte-derived EVs, we isolated EVs from the GBM cell line U87MG and primary normal astrocytes for proteomic analysis (**Fig. 1A**). Our analysis confirmed the presence of several EV marker proteins, including CD63, ARF6, TSG101, and ANXA6^42^, and the absence of an EV-negative marker GM130^43, 44^ in both EV sample groups (**Table S1**). Principal component analysis (PCA) indicated a clear separation between the U87MG-derived EVs and the astrocyte-derived EVs (**Fig. 1B**). Notably, U87MG-derived EVs exhibited a significant abundance of NCL and previously reported angiogenic factors, including vascular endothelial growth factor A (VEGFA) and fibroblast growth factors (FGF2 and FGF13)^22^, compared to astrocyte-derived EVs (**Fig. 1C**). Functional enrichment analysis indicated that the protein profile of U87MG-derived EVs was linked to signaling pathways involved in cell proliferation, migration and angiogenesis (**Fig. 1D**). Flow cytometric results confirmed that NCL was highly expressed on the surface of U87MG cells but not on the surface of astrocytes (**Fig. S1A**), which aligns with previous studies^24, 45^.

**Fig. 1.**
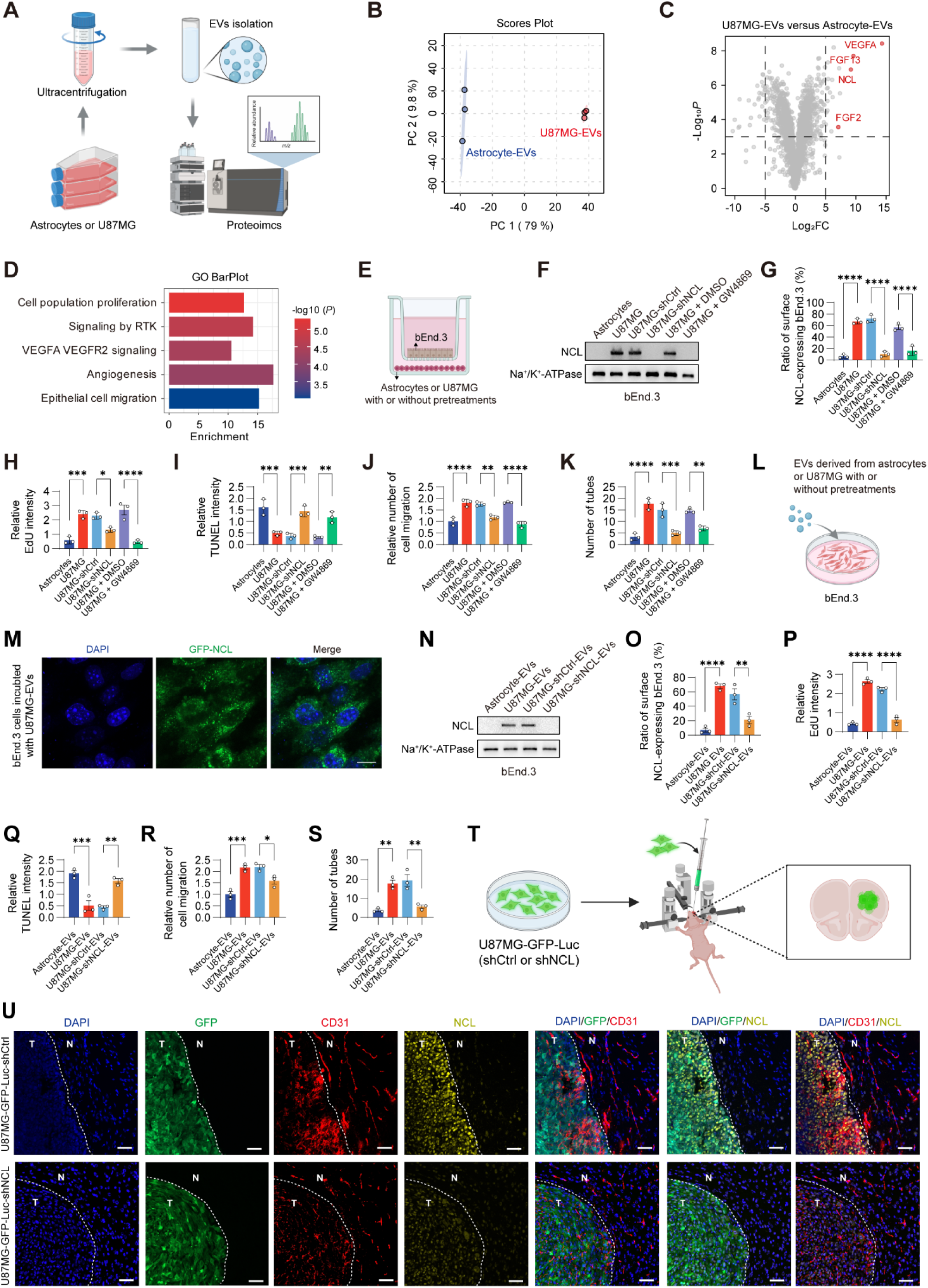
Transfer of NCL from GBM-derived EVs to the surface of brain capillary endothelial cells. **(A)** Schematic of the proteomic analysis for EVs derived from primary astrocytes and the GBM cell line U87MG. **(B)** Principal component analysis (PCA) of proteomic results from astrocyte-derived EVs and U87MG-derived EVs. **(C)** Volcano plot showing proteins with differential abundance between astrocytes-derived EVs and U87MG-derived EVs. P < 0.001, |Log_2_FC| > 5. **(D)** Functional enrichment analysis of proteins carried by U87MG-derived EVs. **(E)** Schematic of the establishment of an *in vitro* BBB or BTB Transwell model, achieved by coculturing bEnd.3 cells with astrocytes, untreated U87MG cells, U87MG cells stably transfected with plasmids encoding scrambled control shRNA (shCtrl) or NCL shRNA (shNCL), or U87MG cells preincubated with DMSO or 10 μM EV release inhibitor GW4869. After the coculture for 5 days, the surface expression of NCL on bEnd.3 cells were detected by Western blotting or flow cytometry. **(F)** Level of surface NCL on bEnd.3 cells after coculture with astrocytes, untreated U87MG cells, or U87MG cells subjected to the above pretreatments. **(G)** The ratio of surface NCL-expressing bEnd.3 cells after coculture with astrocytes, untreated U87MG cells, or U87MG cells subjected to the above pretreatments. **(H)** EdU proliferation assay for bEnd.3 cells after coculture with astrocytes, untreated U87MG cells, or U87MG cells subjected to the above pretreatments. **(I)** TUNEL assay for apoptotic bEnd.3 cells after coculture with astrocytes, untreated U87MG cells, or U87MG cells subjected to the above pretreatments. **(J)** Migration of bEnd.3 cells after coculture with astrocytes, untreated U87MG cells, or U87MG cells subjected to the above pretreatments. **(K)** Tube formation assay for bEnd.3 cells after coculture with astrocytes, untreated U87MG cells, or U87MG cells subjected to the above pretreatments. (**L**) Schematic showing the incubation of bEnd.3 cells with 10 μg/mL EVs derived from astrocytes, untreated U87MG cells, or U87MG cells stably transfected with plasmids encoding shCtrl or shNCL. After incubation for 12 h, surface expression of NCL on bEnd.3 cells were assessed using Western blotting or flow cytometry. **(M)** Confocal imaging of the surface distribution of green fluorescent protein (GFP)-tagged NCL on bEnd.3 cells after incubation with 10 μg/mL U87MG-EVs for 30 min. **(N)** Level of surface NCL on bEnd.3 cells post-incubation with EVs derived from astrocytes, untreated U87MG cells, or U87MG cells subjected to the above pretreatments. **(O)** The ratio of surface NCL-expressing bEnd.3 cells post-incubation with EVs derived from astrocytes, untreated U87MG cells, or U87MG cells subjected to the above pretreatments. (**P**) EdU proliferation assay for bEnd.3 cells after coculture with EVs derived from astrocytes, untreated U87MG cells, or U87MG cells subjected to the above pretreatments. **(Q)** TUNEL assay for apoptotic bEnd.3 cells after coculture with EVs derived from astrocytes, untreated U87MG cells, or U87MG cells subjected to the above pretreatments. **(R)** Migration of bEnd.3 cells after coculture with EVs derived from astrocytes, untreated U87MG cells, or U87MG cells subjected to the above pretreatments. **(S)** Tube formation assay for bEnd.3 cells after coculture with EVs derived from astrocytes, untreated U87MG cells, or U87MG cells subjected to the above pretreatments. (**T**) Schematic illustration of the establishment of an *in vivo* orthotopic GBM mouse model. Briefly, U87MG cells engineered to express firefly luciferase and GFP (U87MG-GFP-Luc cells) were inoculated into the brains of BALB/c nude mice and allowed to grow for 7 days. Before inoculation, U87MG-GFP-Luc cells were stably transfected with plasmids encoding shCtrl or shNCL. (**U**) Immunofluorescence (IF) staining of CD31 (red) and NCL (yellow) of brain sections from orthotopic GBM mice inoculated with U87MG-GFP-Luc cells (green). Nucleus were counterstained with DAPI (blue). N, non-tumor region; T, tumor region. Scale bars, 50 μm. *In vitro* experiments were performed in triplicate and repeated three times. n = 3 for each group of *in vivo* studies. Graphs represented means ± SEM, and statistical significance was calculated by one-way ANOVA (G, H, I, J, K, O, P, Q, R, and S). \**P* < 0.05, \*\**P* < 0.01, \*\*\**P* < 0.001 and \*\*\*\**P* < 0.0001.

To investigate whether NCL carried by GBM-derived EVs mediated the crosstalk between GBM and brain capillary endothelial cells, we established a widely used *in vitro* model of BBB or BTB^46–50^. This model involves coculture of astrocytes or U87MG cells with the brain capillary endothelial cell line bEnd.3 in a Transwell system^46, 50^. Specifically, bEnd.3 cells were grown on matrigel-coated Transwell inserts, while astrocytes or U87MG cells were plated at the bottom of the culture wells (**Fig. 1E**). Following continuous coculture, bEnd.3 cells form tight junctions, effectively mimicking the BBB or BTB^46–50^. We observed no expression of NCL protein on the surface of bEnd.3 cells cocultured with normal astrocytes by Western blotting and flow cytometric analysis. In contrast, a high level of surface NCL protein was detected on bEnd.3 cells cocultured with U87MG cells (**Fig. 1F and 1G**). However, the level of NCL mRNA in bEnd.3 cells cocultured with U87MG cells did not increase when compared to bEnd.3 cells cultured alone or cocultured with astrocytes (**Fig. S1B**). We generated U87MG cells with stable silencing of NCL using shRNA targeting NCL (shNCL) and confirmed reduced NCL level when compared to those cells transfected with scrambled control shRNA (shCtrl) (**Fig. S1C**). When bEnd.3 cells were cocultured with NCL-silenced U87MG cells, they showed a lower surface NCL protein level compared to bEnd.3 cells cocultured with shCtrl-transfected U87MG cells (**Fig. 1F and 1G**). Further, we pretreated U87MG cells with GW4869, an EV release inhibitor^51^, and demonstrated that bEnd.3 cells exhibited a decreased level of surface NCL after their coculture with the GW4869-pretreated U87MG cells (**Fig. 1F and 1G**), suggesting that GBM cells transferred NCL to the surface of capillary endothelial cells via EVs. We also found that bEnd.3 cells, when cocultured with U87MG cells, showed promoted proliferation, migration, and angiogenesis, and decreased apoptosis when compared to those cocultured with astrocytes, as evidenced by 5-ethynyl-2′-deoxyuridine (EdU), Terminal deoxynucleotidyl transferase dUTP nick-end labeling (TUNEL), Transwell and tube formation assays (**Fig. 1H-1K and Fig. S2A**), recapitulating the altered phenotypes and function of the BTB in comparison to the BBB. Conversely, bEnd.3 cells cocultured with shNCL-transfected or GW4869-pretreated U87MG cells showed reduced proliferation, migration, and tube formation, as well as increased apoptosis, compared to those cocultured with shCtrl-transfected or DMSO-pretreated U87MG cells (**Fig. 1H-1K and Fig. S2A**).

To validate the role of EVs in the transfer of NCL from GBM cells to the surface of cocultured capillary endothelial cells, we treated bEnd.3 cells with U87MG- or astrocyte-derived EVs (**Fig. 1L**). Confocal imaging revealed a clear distribution of green fluorescent protein (GFP)-tagged NCL on the surface of bEnd.3 cells following incubation with U87MG-derived EVs (**Fig. 1M**). There was a substantial increase of NCL protein on the surface of bEnd.3 cells treated with U87MG-derived EVs rather than those treated with astrocyte-derived EVs (**Fig. 1N and 1O**). However, bEnd.3 cells incubated with EVs derived from NCL-silenced U87MG cells displayed a reduced level of surface NCL (**Fig. 1N and 1O**). After treatment with U87MG-derived EVs, bEnd.3 cells demonstrated enhanced proliferation, migration, and tube formation and decreased apoptosis compared to those incubated with astrocyte-derived EVs (**Fig. 1P-1S and Fig. S2B**), mimicking the transformation of the BBB into the BTB. In contrast, bEnd.3 cells treated with EVs derived from NCL-silenced U87MG cells exhibited diminished BTB characteristics, as evidenced by reduced proliferation, migration, and tube formation and increased apoptosis of these bEnd.3 cells compared to those cells incubated with shCtrl-transfected U87MG- derived EVs (**Fig. 1P-1S and Fig. S2B**).

To explore the role of NCL transferred from GBM cells to the surface of brain capillary endothelial cells in reshaping the BTB *in vivo*, we established an orthotopic GBM mouse model by inoculating U87MG cells expressing firefly luciferase and GFP (U87MG-GFP-Luc), along with either shNCL or shCtrl, into the brains of BALB/c nude mice (**Fig. 1T**). Immunofluorescence (IF) staining revealed that mice inoculated with shCtrl- transfected U87MG-GFP-Luc cells exhibited a significant increase of CD31-positive endothelial cells within the tumor region compared to the non-tumor brain regions. Both shCtrl-transfected U87MG-GFP-Luc cells and CD31-positive endothelial cells exhibited higher levels of NCL expression than non-tumor brain cells, as evidenced by the colocalization of NCL with both GFP and CD31 (**Fig. 1U**). In contrast, when NCL-silenced U87MG-GFP-Luc cells were inoculated into the brains of nude mice, we observed a reduction in the number of CD31-positive endothelial cells in the tumor region and decreased expression of NCL in these endothelial cells when compared to the mice inoculated with shCtrl-transfected U87MG-GFP-Luc cells (**Fig. 1U**). These findings collectively indicated that NCL transferred from GBM cells to the surface of brain capillary endothelial cells facilitated the transformation of the BBB into the BTB *in vivo*.

### NCL serves as an RMT receptor, inducing the selective transcytosis of AS1411 across the BTB

We added the Cy5-labeled NCL-binding AS1411 into the Transwell inserts of the coculture system to investigate whether AS1411 could penetrate the *in vitro* BBB or BTB model (**Fig. 2A**). We found that AS1411 exhibited a strong binding affinity with bEnd.3 cells cocultured with U87MG cells, but not with bEnd.3 cells cocultured with astrocytes (**Fig. 2B**). When NCL expression was silenced in U87MG cells by shNCL, or when these cells were pretreated with the EV release inhibitor GW4869, the binding affinity of AS1411 with bEnd.3 cells were reduced following their coculture with these modified U87MG cells (**Fig. 2B**). Remarkably, we observed substantial penetration of AS1411 across the *in vitro* BTB model formed by coculturing bEnd.3 cells with U87MG cells, in contrast to the BBB model created by coculturing bEnd.3 cells with astrocytes (**Fig. 2C**). In comparison, a negative control (NC) sequence was incapable of crossing either the BTB or the BBB model (**Fig. 2C**). Silencing NCL expression in U87MG cells or pretreating these cells with GW4869 lowered the penetration ratio of AS1411 across the *in vitro* BTB model after their coculture of bEnd.3 cells, compared to the penetration observed with bEnd.3 cells cocultured with shCtrl-transfected or DMSO-pretreated U87MG cells (**Fig. 2C**).

**Fig. 2.**
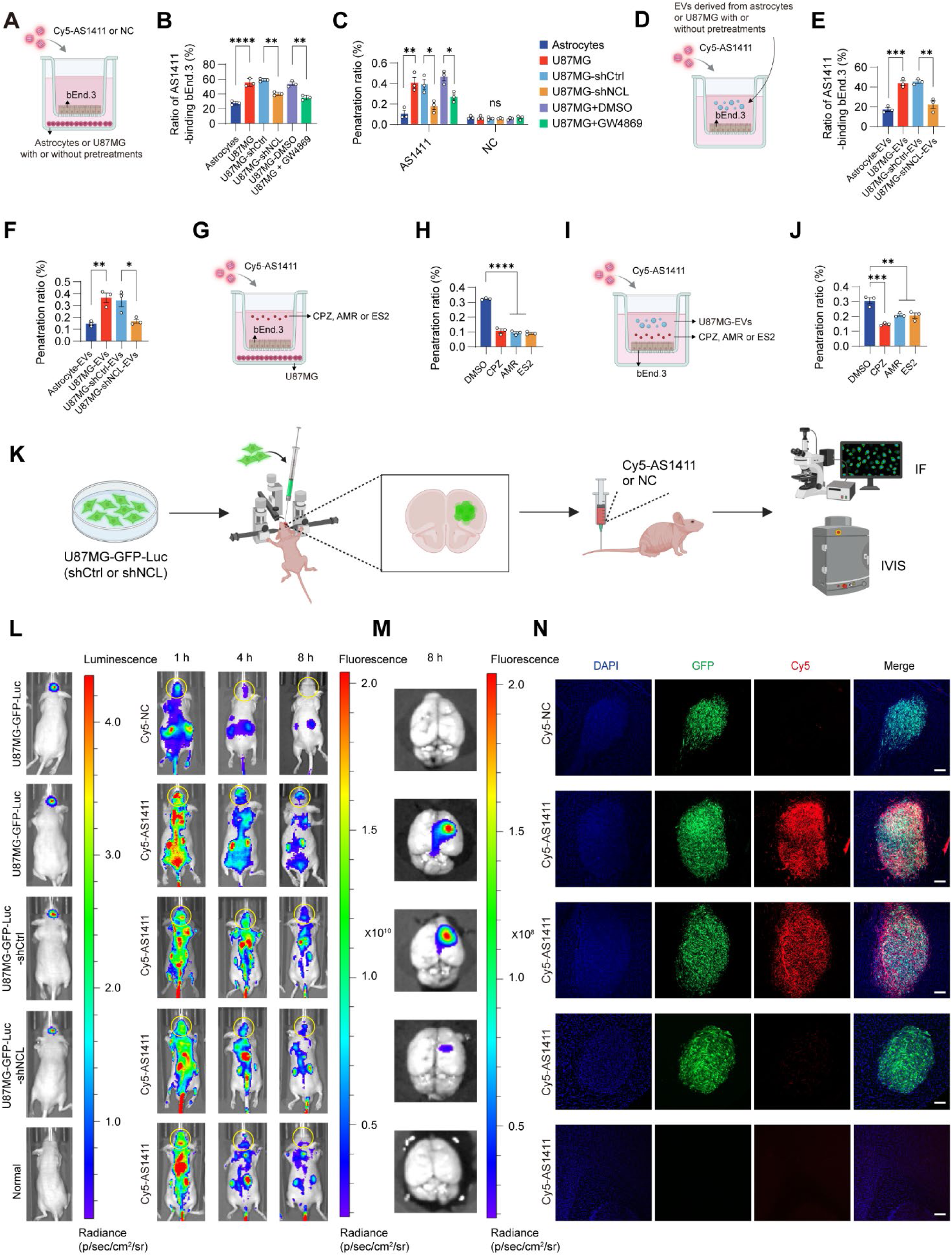
BTB-penetrating and GBM-targeting capacities of AS1411. (**A**) Schematic for evaluating penetration of AS1411 across the BBB or BTB Transwell model. Briefly, bEnd.3 cells were cocultured with astrocytes, untreated U87MG cells, U87MG cells stably transfected with plasmids encoding shCtrl or shNCL, or U87MG cells preincubated with DMSO or GW4869 for 5 days for the establishment of the BBB or BTB Transwell model. 200 nM Cy5-labeled AS1411 or negative control (NC) was added in the Transwell inserts for incubating another 6 h before the measurement of the amount of BBB- or BTB-penetrating AS1411 or NC by a microplate reader. (**B**) The ratio of AS1411-binding bEnd.3 cells post-coculture with astrocytes, untreated U87MG cells, or U87MG cells subjected to the above pretreatments. (**C**) Penetration ratio of AS1411 or NC across the BBB or BTB Transwell model. After coculture of bEnd.3 cells with astrocytes, untreated U87MG cells, or U87MG cells subjected to the above pretreatments for 5 days, the penetration ratio was calculated by measuring the amount of Cy5-labeled AS1411 or NC transferred from inside the Transwell insert to outside, normalized to the total amount of Cy5-labeled AS1411 or NC initially added to the insert. (**D**) Schematic for evaluating penetration of AS1411 across the BBB or BTB Transwell model following incubation of bEnd.3 cells with 10 μg/mL EVs derived from astrocytes, untreated U87MG cells, or U87MG cells stably transfected with plasmids encoding shCtrl or shNCL. After incubation for 12 h, Cy5-labeled AS1411 was added to the Transwell inserts for incubating another 6 h before measurement by a microplate reader. (**E**) Ratio of AS1411-binding bEnd.3 cells after incubation with EVs derived from astrocytes, untreated U87MG cells, or U87MG cells subjected to the above pretreatments. (**F**) Penetration ratio of AS1411 across the BBB or BTB Transwell model after incubating bEnd.3 cells with EVs derived from astrocytes, untreated U87MG cells, or U87MG cells subjected to the above pretreatments. (**G**) Schematic for evaluating penetration of AS1411 across the BTB Transwell model. After the coculture of bEnd.3 cells with U87MG cells for 5 days, 200 nM Cy5-labeled AS1411 was added in the Transwell inserts for incubating another 6 h, with or without the presence of 50 μM Chlorpromazine (CPZ, an inhibitor of clathrin-mediated endocytosis), 1 mM Amiloride (AMR, an inhibitor of macropinocytosis), or 250 μM exocytosis inhibitor Endosidin-2 (ES2). (**H**) Penetration ratio of AS1411 across the *in vitro* BTB model following coculture of bEnd.3 cells with U87MG cells, with or without the presence of the above treatments. (**I**) Schematic for evaluating the penetration of AS1411 across the BTB Transwell model following incubation of bEnd.3 cells with 10 μg/mL U87MG-EVs. After incubation for 12 h, Cy5-labeled AS1411 was added to the Transwell inserts for incubating another 6 h, with or without the presence of CPZ, AMR, or ES2. (**J**) Penetration ratio of AS1411 across the *in vitro* BTB model following incubation with U87MG-derived EVs, with or without the presence of the above treatments. (**K**) Schematic illustration of the establishment of an *in vivo* orthotopic GBM mouse model. Briefly, U87MG-GFP-Luc cells stably transfected with plasmids encoding shCtrl or shNCL were inoculated into the brains of BALB/c nude mice and allowed to grow for 7 days. Then, the GBM-bearing mice receive 0.2 μmol/kg Cy5-labeled NC or AS1411 via vein injection. *In vivo* imaging systems (IVIS) and IF were used to assess the brain distribution of Cy5-labeled NC or AS1411. (**L**) *In vivo* brain distribution of Cy5-labeled NC or AS1411 at 1, 4, and 8 h. (**M**) *Ex vivo* analysis of brain distribution of Cy5-labeled NC or AS1411 at 8 h. (**N**) Co-localization of Cy5-labeled NC or AS1411 (red) with U87MG-GFP-Luc cells (green) in brain sections at 8 h. Scale bars, 200 μm. *In vitro* experiments were performed in triplicate and repeated three times. n = 3 for each group of *in vivo* studies. Graphs represented means ± SEM, and statistical significance was calculated by one-way ANOVA (B, C, E, F, H, and J). \**P* < 0.05, \*\**P* < 0.01, \*\*\**P* < 0.001 and \*\*\*\**P* < 0.0001. ns, no significance.

We treated bEnd.3 cells in the Transwell inserts with EVs derived from either U87MG cells or astrocytes and then added Cy5-labeled AS1411 into the inserts (**Fig. 2D**). The results demonstrated that AS1411 exhibited a strong binding affinity with bEnd.3 cells treated with U87MG-derived EVs, whereas it showed little binding with bEnd.3 cells treated with astrocyte-derived EVs (**Fig. 2E**). Moreover, when bEnd.3 cells were treated with EVs from NCL-silenced U87MG cells, AS1411 displayed significantly reduced binding affinity compared to those treated with EVs from shCtrl-transfected U87MG cells (**Fig. 2E**). Additionally, the penetration of AS1411 across the *in vitro* BTB model was notably enhanced when bEnd.3 cells were cocultured with U87MG-derived EVs, in contrast to those cocultured with astrocyte-derived EVs (**Fig. 2F**). However, when bEnd.3 cells were treated with EVs derived from NCL-silenced U87MG cells, the penetration of AS1411 across the *in vitro* BTB model was reduced compared to those treated with EVs derived from shCtrl-transfected U87MG cells (**Fig. 2F**).

The RMT process involves a series of steps, including receptor-ligand binding, endocytosis, intracellular trafficking, and exocytosis^52^, facilitating the transport of therapeutic cargo from apical to basolateral plasma membranes of brain capillary endothelial cells^53^. To determine whether the penetration of AS1411 across the BTB occurred via an RMT-dependent pathway, we added endocytic or exocytic inhibitors into the Transwell inserts of the coculture system prior to assessing the transport of AS1411 across the *in vitro* BTB model (**Fig. 2G**). We showed that either Chlorpromazine (CPZ, an inhibitor of clathrin-mediated endocytosis)^54^, Amiloride (AMR, an inhibitor of macropinocytosis)^55^, or Endosidin-2 (ES2, an inhibitor of exocytosis)^56^ reduced the penetration of AS1411 across the *in vitro* BTB model when bEnd.3 cells were cocultured with U87MG cells in the Transwell system (**Fig. 2H**). We also incubated bEnd.3 cells with EVs derived from U87MG cells in the Transwell inserts and added AS1411 into the inserts, with or without the presence of CPZ, AMR, or ES2 (**Fig. 2I**). Our results demonstrated that the BTB-penetrating ability of AS1411 was inhibited by CPZ, AMR, or ES2 (**Fig. 2J**). These results indicate that NCL transferred from GBM-derived EVs serves as an RMT receptor on BTB capillary endothelial cells rather than on BBB endothelial cells, facilitating the selective transcytosis of AS1411 across the BTB.

We evaluated the *in vivo* BTB-penetrating capacity of AS1411 using the orthotopic GBM mouse models inoculated with U87MG-GFP-Luc cells. Cy5-labeled AS1411 or NC sequence was intravenously injected into the mice and fluorescent intensities were detected at 1, 4, and 8 h after the injection using the *in vivo* imaging system (IVIS) (**Fig. 2K**). The strong bioluminescent signal from firefly luciferase indicated the development of GBM in mice inoculated with U87MG-GFP-Luc cells (**Fig. 2L**). There was an abundant accumulation of fluorescent intensity of Cy5-labelled AS1411 in brain tissues of GBM mice at different time points, whereas the fluorescent signal of the Cy5-labelled NC sequence was not detectable in brain tissues of GBM mice (**Fig. 2L**). When inoculating NCL-silenced U87MG-GFP-Luc cells into the brains of mouse models, we observed less distribution of AS1411 in the brain tissues when compared to mice inoculated with shCtrl-transfected U87MG- GFP-Luc cells (**Fig. 2L**). Additionally, normal mice with no inoculation of U87MG-GFP-Luc cells did not exhibit the brain distribution of AS1411 (**Fig. 2L**). *Ex vivo* analysis confirmed that AS1411 could penetrate the BTB rather than the BBB, and the BTB-penetrating ability of AS1411 was dependent on the GBM-derived NCL (**Fig. 2M**).

### AS1411 exhibits GBM-targeting and -internalizing ability

We determined the GBM-targeting and -internalizing ability of AS1411 *in vitro*. Flow cytometric analysis revealed that the binding ability of Cy5-labeled AS1411 with U87MG cells was significantly higher than that of the Cy5-labeled NC sequence (**Fig. S3A**). Confocal imaging showed that Cy5-labeled AS1411 was internalized into U87MG cells, whereas no entry of Cy5-labeled NC sequence into U87MG cells was observed (**Fig. S3B**). Similarly, we also demonstrated that AS1411 exhibited stronger binding affinity with another GBM cell line, U251, compared to the NC sequence *in vitro* (**Fig. S3C**). The uptake of AS1411 into U251 cells was more pronounced than that of the NC sequence (**Fig. S3D**). In contrast, AS1411 displayed no binding ability with either astrocytes or a non-tumorigenic epithelial cell line MCF 10A, and internalization of AS1411 into these cells was not detectable *in vitro* (**Fig. S3E-S3H**).

Multicellular tumor spheroids are considered promising *in vitro* 3D models that bridge the gap between *in vitro* 2D cellular monolayers and *in vivo* animal models^57^, as they more closely mimic the characteristics of solid tumors, such as spatial distribution of tumor cells, extracellular matrix, cell adhesion, hypoxia, and acidic microenvironment^58^. We established 3D spheroids of U87MG and U251 cells and observed that Cy5-labeled AS1411 could penetrate these spheroids, whereas the NC sequence was incapable of entering the spheroids (**Fig. S4A-S4C**). When silencing NCL in spheroids of U87MG cells, AS1411 could not penetrate the spheroids, suggesting that the penetration of AS1411 into GBM spheroids is NCL-dependent (**Fig. S4D-S4F**).

To determine the GBM-targeting ability of AS1411 *in vivo*, we utilized the orthotopic GBM mouse models inoculated with U87MG-GFP-Luc cells. Our IF staining results showed that AS1411, rather than the NC sequence, was colocalized with U87MG-GFP-Luc cells, but not with non-tumor cells on brain tissue sections from GBM mice (**Fig. 2N**). Silencing of NCL decreased the *in vivo* colocalization of AS1411 with U87MG-GFP-Luc cells compared to shCtrl-transfected U87MG-GFP-Luc cells (**Fig. 2N**). No distribution of AS1411 was observed on brain tissue sections from normal mice without inoculation of U87MG-GFP-Luc cells (**Fig. 2N**). All the above results suggest that AS1411 specifically targets and enters GBM cells.

### AS1411-based PROTACs trigger the spatial proximity of the NCL-MDM2 complex with POIs

Since we previously proved that AS1411 could be repurposed as an MDM2 recruiter in PROTACs via forming an NCL-bridged complex, *i.e.*, AS1411-NCL-MDM2^36^, we aimed to develop AS1411-based PROTACs for treating GBM by degrading VEGFR2 or EGFR, as both are promising POIs for GBM^37–39^. Our co-immunoprecipitation (IP) assays confirmed that NCL was associated with MDM2, forming a complex without involving VEGFR2 and EGFR, in U87MG cells (**Fig. 3A**). We labeled AS1411 with biotin and conducted pull-down assays using streptavidin-coated magnetic beads. Our results showed that AS1411 captured both NCL and MDM2, but not VEGFR2 or EGFR, in U87MG cells (**Fig. 3B**). However, when silencing NCL in U87MG cells, AS1411 could not capture MDM2 (**Fig. 3C**), suggesting that the recruitment of MDM2 by AS1411 was dependent on NCL. We also observed the *in vivo* colocalization of AS1411 with both NCL and MDM2 in the tumor region of brain tissue sections from GBM mice (**Fig. S5A and S5B**). We then conjugated AS1411 with a VEGFR2 inhibitor Ced^40^ or an EGFR inhibitor Gef^41^ via a linear alkyl linker, resulting in the creation of hetero-bifunctional molecules termed AS1411-Ced (**Fig. 3D and Scheme 1**) or AS1411-Gef (**Fig. 3E and Scheme 2**). AS1411 is a G-rich aptamer capable of folding into a G-quadruplex structure, which contributes to its specific binding with NCL^59^. N-methylmesoporphyrin IX (NMM) is a widely utilized fluorescent probe for the easy detection of G-quadruplex^60^. After incubation with NMM, we observed the comparable fluorescent absorption spectra among AS1411, AS1411-Ced, and AS1411-Gef (**Fig. 3F**), suggesting that the conjugation of AS1411 with Ced or Gef could not compromise the NCL-binding ability of AS1411.

**Fig. 3.**
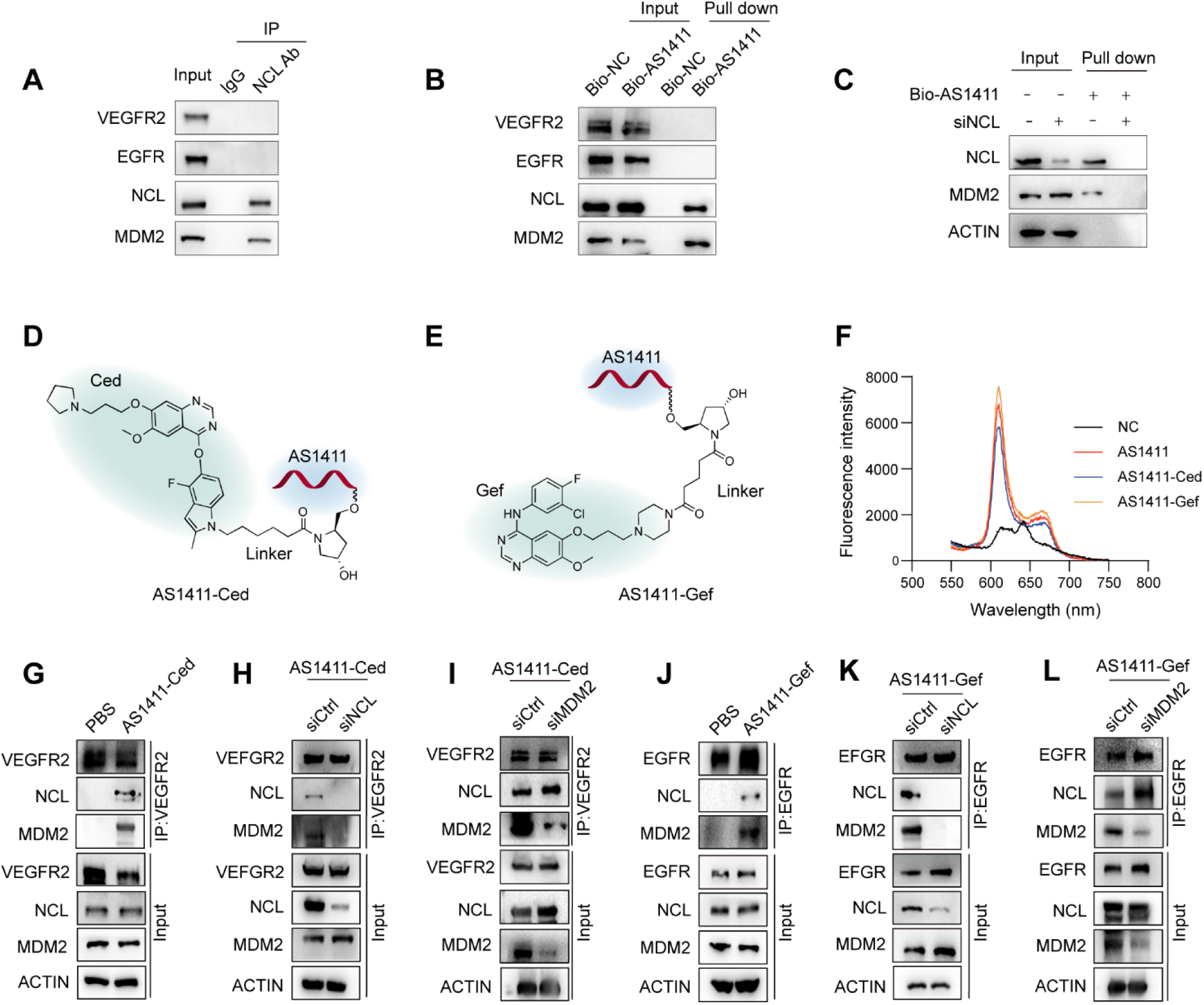
Formation of a MDM2-NCL-PROTAC-POI quaternary complex. (**A**) Co-immunoprecipitation (IP) of NCL, MDM2, VEGFR2, and EGFR in U87MG cells using isotype-matched IgG or an anti-NCL antibody. (**B**) Pull-down assay of NCL, MDM2 VEGFR2, and EGFR in U87MG cells treated with 500 nM biotin-labeled NC (Bio-NC) or AS1411 (Bio-AS1411) for 6 h by streptavidin-coated magnetic beads. (**C**) Pull-down assay of NCL and MDM2 in siNCL-transfected U87MG cells after treatment with 500 nM Bio-NC or Bio-AS1411 for 6 h. (**D**) Structure of AS1411-cediranib (AS1411-Ced). (**E**) Structure of AS1411-gefitinib (AS1411-Gef). (**F**) Absorption spectra of N-methylmesoporphyrin IX (NMM) in the presence of 10 µM NC, AS1411, AS1411-Ced, and AS1411-Gef. The concentration of NMM was 1 µM. (**G**) Co-IP of NCL, MDM2, and VEGFR2 in U87MG cells treated with PBS or 500 nM AS1411-Ced for 6 h using an anti-VEGFR2 antibody in the presence of 10 µM MG132. (**H**) Co-IP of NCL, MDM2, and VEGFR2 in scrambled control siRNA (siCtrl)- or NCL siRNA (siNCL)- transfected U87MG cells after treatment with PBS or 500 nM AS1411-Ced for 6 h using an anti-VEGFR2 antibody in the presence of 10 µM MG132. (**I**) Co-IP of NCL, MDM2, and VEGFR2 in siCtrl or siMDM2- transfected U87MG cells after treatment with PBS or 500 nM AS1411-Ced for 6 h using an anti-VEGFR2 antibody in the presence of 10 µM MG132. (**J**) Co-IP of NCL, MDM2, and EGFR in U87MG cells treated with PBS or 500 nM AS1411-Gef for 6 h using an anti-EGFR antibody in the presence of 10 µM MG132. (**K**) Co-IP of NCL, MDM2, and EGFR in siCtrl- or siNCL-transfected U87MG cells after treatment with PBS or 500 nM AS1411-Gef for 6 h using an anti-EGFR antibody in the presence of 10 µM MG132. (**L**) Co-IP of NCL, MDM2, and EGFR in siCtrl- or siMDM2-transfected U87MG cells after treatment with PBS or 500 nM AS1411-Gef for 6 h using an anti-EGFR antibody in the presence of 10 µM MG132. Experiments were performed in triplicate and repeated three times.

To test whether AS1411-based PROTACs could induce spatial proximity between the NCL-MDM2 complex and POIs, respectively, we treated U87MG cells with PBS or AS1411-based PROTACs and performed co-IP assays. Our results demonstrated that NCL and MDM2 were coprecipitated with VEGFR2 in U87MG cells incubated with AS1411-Ced, but not in cells treated with PBS (**Fig. 3G**), indicating that AS1411-Ced enabled the formation of a quaternary complex involving MDM2, NCL, VEGFR2 and itself. When NCL was silenced in U87MG cells, we observed that VEGFR2 failed to coprecipitate MDM2 in the presence of AS1411-Ced (**Fig. 3H**). However, VEGFR2 still was coprecipitated with NCL in MDM2-silenced U87MG cells treated with AS1411-Ced (**Fig. 3I**). These findings suggest that NCL acts as a molecular bridge between MDM2 and VEGFR2, facilitating the assembly of the MDM2-NCL-PROTAC-VEGFR2 quaternary complex. Similarly, NCL and MDM2 were effectively coprecipitated with EGFR in U87MG cells incubated with AS1411-Gef (**Fig. 3J**). Silencing of NCL destroyed the coprecipitation of MDM2 with EGFR in U87MG cells (**Fig. 3K**), while silencing of MDM2 did not affect the coprecipitation of NCL with EGFR (**Fig. 3L**), suggesting that AS1411-Gef induced the formation of a quaternary complex involving MDM2, NCL, EGFR and itself, with NCL acting as a molecular bridge between MDM2 and EGFR.

### AS1411-based PROTACs induce degradation of POIs in both NCL- and MDM2-dependent manners

To assess whether AS1411-Ced could induce VEGFR2 degradation, we treated U87MG or U251 cells with increasing concentrations and varied incubation times of AS1411-Ced. Our results showed that AS1411-Ced decreased the VEGFR2 protein level in both cell lines in both concentration- and time-dependent manners (**Fig. 4A and 4B**). We treated U87MG or U251 cells with PBS, AS1411, Ced, AS1411 + Ced (a physical combination of AS1411 and Ced), or AS1411-Ced. In contrast to PBS, AS1411-Ced induced the degradation of VEGFR2, whereas AS1411, Ced, or AS1411 + Ced had no effect on VEGFR2 level (**Fig. 4C**), suggesting that the spatial proximity of the NCL-MDM2 complex with VEGFR2 induced by AS1411-Ced was responsible for the degradation of VEGFR2. IF staining results confirmed that AS1411-Ced reduced the VEGFR2 protein level in U87MG cells (**Fig. 4D**). Proteomic analysis validated that AS1411-Ced specifically degraded VEGFR2 in U87MG cells (**Fig. 4E**). Pathway enrichment analysis revealed that AS1411-Ced remarkably downregulated VEGFR2-mediated signaling pathways, including those involved in cell migration, cell cycle, blood vessel development, and epithelial cell differentiation and upregulated signaling pathways related to cell apoptosis and growth inhibition (**Fig. 4F**). We also evaluated whether AS1411-Gef were effective PROTACs for degrading EGFR. Treatment of U87MG and U251 cells with AS1411-Gef resulted in a concentration- and time-dependent decrease in EGFR protein level (**Fig. 4G and 4H**). AS1411-Gef, rather than AS1411, Gef, or AS1411 + Gef, induced the degradation of EGFR compared to PBS (**Fig. 4I**). IF staining and proteomic results confirmed the reduction of EGFR protein level in U87MG cells treated with AS1411-Gef (**Fig. 4J and Fig. 4K**). Pathway enrichment analysis revealed that AS1411-Gef downregulated EGFR-mediated signaling pathways involved in cell proliferation, epithelial cell migration, and differentiation, while promoted signaling pathways related to cell death and survival suppression (**Fig. 4L**).

**Fig. 4.**
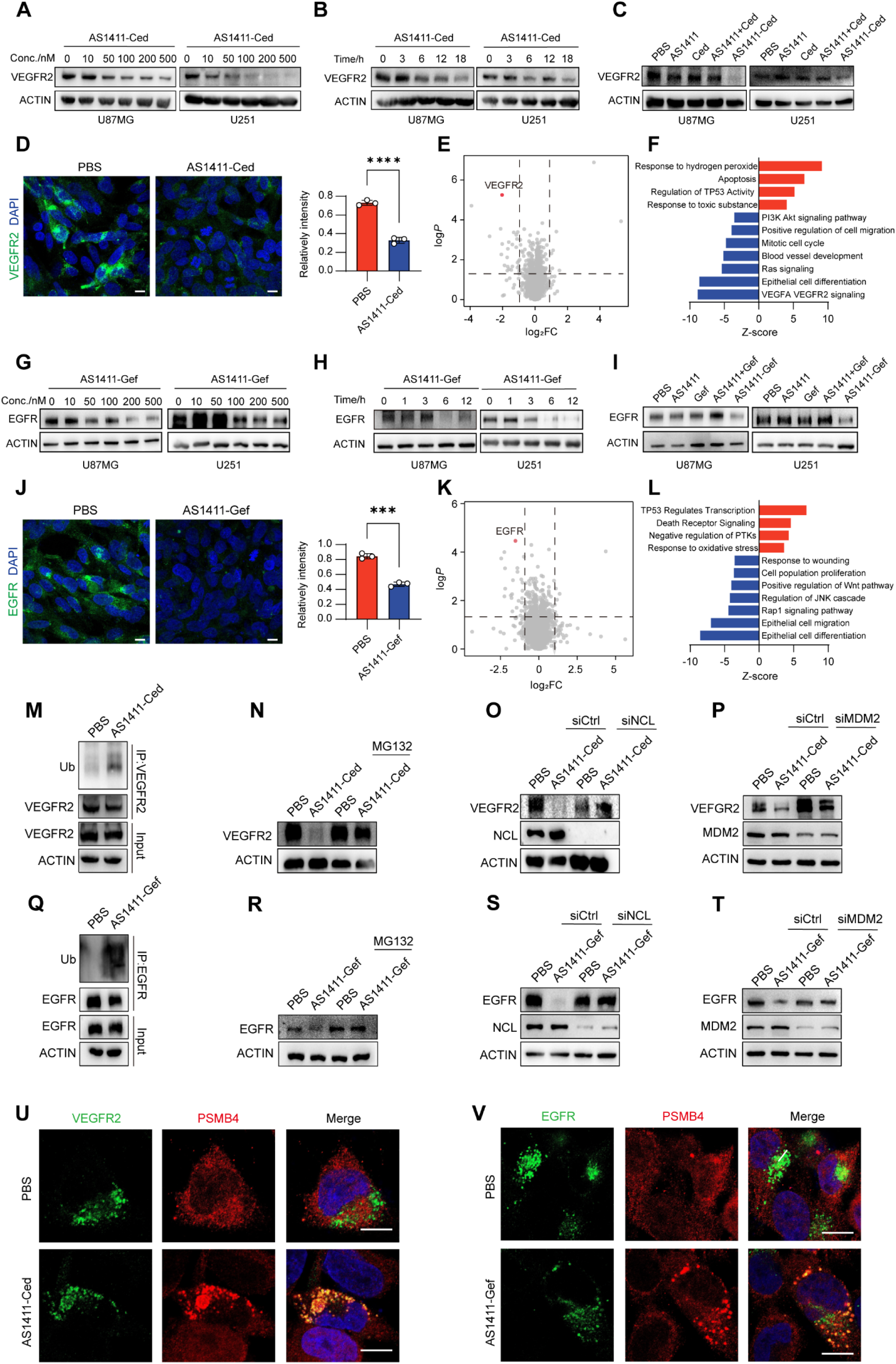
Degradation of VEGFR2 or EGFR by AS1411-based PROTACs. (**A**) Level of VEGFR2 in U87MG and U251 cells after treatment with increasing concentrations of AS1411-Ced for 18 h. (**B**) Level of VEGFR2 in U87MG and U251cells after treatment with 200 nM AS1411-Ced at increasing incubation times. (**C**) Level of VEGFR2 in U87MG and U251 cells after treatment with PBS, AS1411, Ced, AS1411+Ced, or AS1411-Ced at a concentration of 200 nM for 18 h. (**D**) IF staining of VEGFR2 in U87MG cells after treatment with PBS or 200 nM AS1411-Ced for 18 h. Scale bars, 10 µm. (**E**) Volcano plot displaying differentially abundant proteins in U87MG cells treated with PBS or 200 nM AS1411-Ced for 18 h, as determined by proteomics. P value < 0.05, Fold change > 2. **(F)** Functional enrichment analysis of the differentially abundant proteins in U87MG cells treated with PBS or 200 nM AS1411-Ced for 18 h. (**G**) Level of EGFR in U87MG and U251 cells after treatment with increasing concentrations of AS1411-Gef for 12 h. (**H**) Level of EGFR in U87MG and U251 cells after treatment with 200 nM AS1411-Gef at increasing incubation times. (**I**) Level of EGFR in U87MG and U251 cells after treatment with PBS, AS1411, Gef, AS1411+Gef, or AS1411-Gef at a concentration of 200 nM for 12 h. (**J**) IF staining of EGFR in U87MG cells after treatment with PBS or 200 nM AS1411-Gef for 12 h. (**K**) Volcano plot displaying the differentially abundant proteins in U87MG cells treated with PBS or 200 nM AS1411-Gef for 12 h, as determined by proteomics. P value < 0.05, Fold change > 2. **(L)** Functional enrichment analysis of the differentially abundant proteins in U87MG cells treated with PBS or 200 nM AS1411-Gef for 12 h. (**M**) Level of VEGFR2 ubiquitination in U87MG cells after treatment with PBS or 200 nM AS1411-Ced for 6 h in the presence of 10 µM MG132. (**N**) Level of VEGFR2 in U87MG cells after treatment with PBS or 200 nM AS1411-Ced for 18 h, in the presence or absence of 10 μM MG132. (**O**) Level of VEGFR2 in siCtrl- or siNCL-transfected U87MG cells after treatment with PBS or 200 nM AS1411-Ced for 18 h. (**P**) Level of VEGFR2 in siCtrl- or siMDM2-transfected U87MG cells after treatment with PBS or 200 nM AS1411-Ced for 18 h. (**Q**) Level of EGFR ubiquitination in U87MG cells after treatment with PBS or 200 nM AS1411-Gef for 6 h in the presence of 10 µM MG132. (**R**) Level of EGFR in U87MG cells after treatment with PBS or 200 nM AS1411-Gef for 12 h, in the presence or absence of 10 μM MG132. (**S**) Level of EGFR in siNC- or siNCL-transfected U87MG cells after treatment with PBS or 200 nM AS1411-Gef for 12 h in the presence of 10 µM MG132. (**T**) Level of EGFR in siCtrl- or siMDM2-transfected U87MG cells after treatment with PBS or 200 nM AS1411-Gef for 12 h. (**U**) Confocal analysis of the co-localization of VEGFR2 with proteasome subunit PSMB4 in U87MG cells treated with PBS or 200 nM AS1411-Ced for 18 h. Scale bars, 10 μm. (**V**) Confocal analysis of the co-localization of EGFR with proteasome subunit PSMB4 in U87MG cells after treatment with PBS or 200 nM AS1411-Gef for 12 h. Scale bars, 10 μm. Experiments were performed in triplicate and repeated three times. Graphs represented means ± SEM and statistical significance was calculated by Student’s *t*-test (D and J). \*\*\**P* < 0.001 and \*\*\*\**P* < 0.0001.

We investigated whether AS1411-Ced and AS1411-Gef utilized the ubiquitin-proteasome pathway to trigger the degradation of VEGFR2 and EGFR, respectively. We showed that AS1411-Ced led to an increase in the ubiquitination of VEGFR2 in U87MG cells (**Fig. 4M**). The addition of the proteasome inhibitor MG132 inhibited the degradation of VEGFR2 induced by AS1411-Ced (**Fig. 4N**). Furthermore, silencing of NCL or MDM2 abolished the AS1411-Ced-mediated degradation of VEGFR2 in U87MG cells (**Fig. 4O and 4P**). In parallel, we also demonstrated that AS1411-Gef promoted the ubiquitination of EGFR in U87MG cells (**Fig. 4Q**). The presence of MG132 prevented the degradation of EGFR induced by AS1411-Gef (**Fig. 4R**). Treatment of NCL- or MDM2-silenced U87MG cells with AS1411-Gef could not trigger the efficient degradation of EGFR (**Fig. 4S and 4T**). Proteasome subunit beta type 4 (PSMB4) is a non-catalytic component of the 20S core proteasome complex^61^. IF imaging showed the colocalization of PSMB4 with VEGFR2 or EGFR in U87MG cells following treatment with AS1411-Ced or AS1411-Gef (**Fig. 4U and 4V**). These results indicate that AS1411-Ced and AS1411-Gef induce the proteasomal degradation of VEGFR2 and EGFR in both NCL- and MDM2-dependent manners, respectively.

### AS1411-based PROTACs display anti-GBM effects *in vitro*

We evaluated the anti-GBM potential of AS1411-Ced *in vitro* using U87MG and U251. Both cell lines were treated with PBS, AS1411, Ced, AS1411 + Ced, and AS1411-Ced, respectively. Cell proliferation was measured using the Cell Counting Kit-8 (CCK-8) assay and EdU staining, which revealed that AS1411-Ced significantly inhibited the proliferation of both U87MG and U251 cells compared to PBS, AS1411, Ced, or AS1411 + Ced (**Fig. 5A, 5B, S6A, and S6B**). Flow cytometric analysis demonstrated that AS1411-Ced markedly increased the apoptotic rate in both cell lines (**Fig. 5C and S6C**). To assess the impact of AS1411- Ced on cell invasion, we employed a 3D spheroid invasion assay, which mimics the native 3D tumor microenvironment by embedding tumor spheroids in an extracellular matrix^62, 63^. Notably, AS1411-Ced treatment significantly attenuated the invasion of U87MG cell spheroids (**Fig. 5D**). Similarly, we evaluated the *in vitro* effects of AS1411-Gef on proliferation, apoptosis, and invasion of GBM cells. AS1411-Gef demonstrated high efficacy in reducing the proliferation of U87MG and U251 cells compared to other treatments, as evidenced by CCK-8 and EdU staining assays (**Fig. 5E, 5F, S6D, and S6E**). Flow cytometric assays showed that AS1411-Gef significantly increased the apoptotic rate in both cell lines (**Fig. 5G and S6F**). AS1411-Gef decreased the invasion of U87MG cell spheroids in the 3D spheroid invasion assays (**Fig. 5H**).

**Fig. 5.**
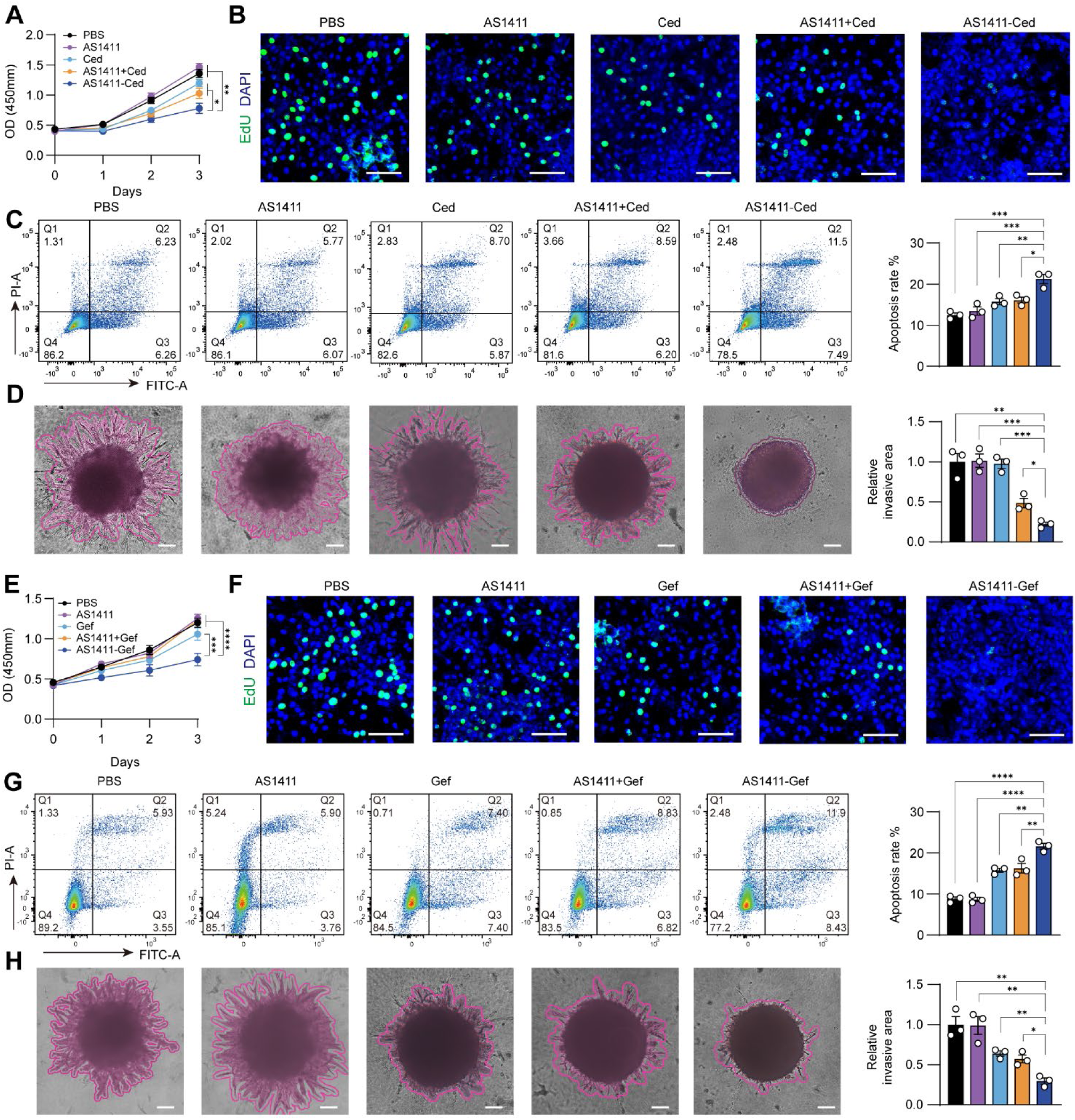
Anti-GBM activity of AS1411-based PROTACs *in vitro*. (**A**) CCK-8 assay for the viability of U87MG cells after 3-day treatment with PBS, AS1411, Ced, AS1411+Ced, or AS1411-Ced at a concentration of 200 nM every day. (**B**) EdU staining of U87MG cells after 3-day treatment with PBS, AS1411, Ced, AS1411+Ced, or AS1411-Ced at a concentration of 200 nM every day. Scale bars, 50 μm. (**C**) Flow cytometry analysis of apoptosis of U87MG cells after 3-day treatment with PBS, AS1411, Ced, AS1411+Ced, or AS1411-Ced at a concentration of 200 nM every day. (**D**) 3D spheroid cell invasion assay of U87MG cells after 1-day treatment with PBS, AS1411, Ced, AS1411+Ced, or AS1411-Ced at a concentration of 200 nM. Scale bars, 200 μm. (**E**) CCK-8 assay for the viability of U87MG cells after 3-day treatment with PBS, AS1411, Gef, AS1411+Gef, or AS1411-Gef at a concentration of 200 nM every day. (**F**) EdU staining of U87MG cells after 3-day treatment with PBS, AS1411, Gef, AS1411+Gef, or AS1411-Gef at a concentration of 200 nM every day. Scale bars, 50 μm. (**G**) Flow cytometry analysis of apoptosis of U87MG cells after 3-day treatment with PBS, AS1411, Gef, AS1411+Gef, or AS1411-Gef at a concentration of 200 nM every day. (**H**) 3D spheroid cell invasion assay of U87MG cells after 1-day treatment with PBS, AS1411, Gef, AS1411+Gef, or AS1411-Gef at a concentration of 200 nM. Scale bars, 200 μm. Graphs represented means ± SEM and statistical significance was calculated by two-way ANOVA (A and E) and one-way ANOVA (C, D, G, and H). \**P* < 0.05, \*\**P* < 0.01, \*\*\**P* < 0.001 and \*\*\*\**P* < 0.0001.

### AS1411-based PROTACs exhibit robust antitumor potency in orthotopic GBM xenografts

We further evaluate the *in vivo* therapeutic effects of AS1411-Ced in the orthotopic GBM mouse models with inoculation of U87MG-GFP-Luc cells. Seven days post-xenograft, the mice were randomly grouped and intravenously received PBS, AS1411, Ced, AS1411 + Ced, and AS1411-Ced every other day for 8 days (**Fig. 6A**). Mice were monitored daily for GBM progression indicators, including seizures, ataxia, and weight loss, and were euthanized if symptomatic tumor burden developed^64^. Brain tumor growth was assessed using bioluminescent imaging via IVIS and survival rates were analyzed by Kaplan-Meier curves^65^. Mice treated with AS1411-Ced exhibited remarkable tumor growth inhibition and extended median survival compared to those receiving PBS, AS1411, Ced, or AS1411 + Ced (**Fig. 6B-6D**). Histological examinations of whole-brain sections by hematoxylin and eosin (H&E) staining revealed smaller GBM tumors from the mice treated with AS1411- Ced compared to those from the mice in other treatment groups (**Fig. 6E**). IF analysis of GBM tumor sections showed a significant reduction in VEGFR2 protein level and fewer Ki-67-positive proliferative cells in the mice treated with AS1411-Ced (**Fig. 6F and 6G)**. TUNEL assays demonstrated that AS1411-Ced treatment induced substantial apoptosis in GBM tissues (**Fig. 6F and 6G**). Notably, AS1411-Ced did not significantly affect the body weight (**Fig. S7A**), or alter levels of hepatic and renal function markers, including alanine aminotransferase (ALT), aspartate aminotransferase (AST), albumin (ALB) and alkaline phosphatase (ALP), as confirmed by serum biochemical assays (**Fig. S7B**). H&E staining of non-tumor organs (heart, liver, spleen, lung, and kidney) showed no obvious systemic toxicity or abnormality in AS1411-Ced-treated mice (**Fig. S7C**). To determine the therapeutic potential of AS1411-Gef for GBM, the orthotopic GBM mouse models were administered with PBS, AS1411, Gef, AS1411 + Gef, and AS1411-Gef via tail vein injection every other day for 8 days (**Fig. S8A**). Mice treated with AS1411-Gef showed reduced tumor size and extended median survival when compared to other treatment groups, as evidenced by bioluminescent imaging, Kaplan-Meier curve, and HE staining (**Fig. S8B-S8E**). IF and TUNEL staining of GBM tissues from AS1411-Gef-treated mice showed decreased EGFR protein levels, reduced Ki-67-positive cells, and increased apoptotic activity (**Fig. S8F and S8G**). Furthermore, AS1411-Gef treatment did not significantly alter body weight, serum hepatic and renal function parameters (ALT, AST, ALB, and ALP), or the histology of non-tumor organs (heart, liver, spleen, lung, and kidney) compared to other treatment groups (**Fig. S9A-S9C**).

**Fig. 6.**
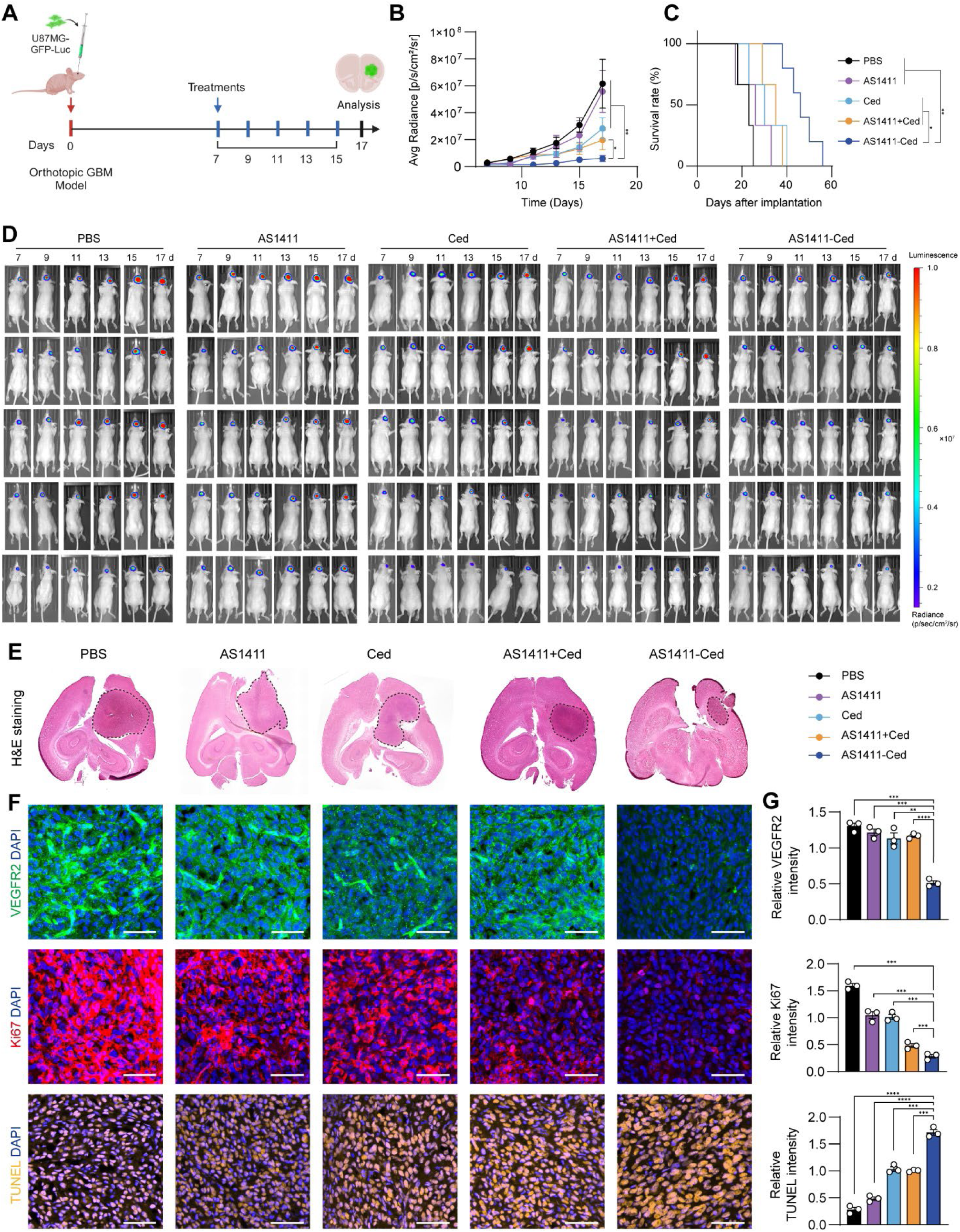
Anti-GBM efficacy of AS1411-Ced *in vivo*. (**A**) Schematic illustration for determining anti-GBM efficacy of AS1411-Ced *in vivo*. Briefly, U87MG-GFP-Luc cells were inoculated into the brains of BALB/c nude mice for 7 days. The orthotopic GBM mice were systemically administrated with PBS, AS1411, Ced, AS1411+Ced, or AS1411-Ced every other day for 8 days. The dose of AS1411-Ced was 4.5 μmol/kg. (**B**) The bioluminescent signal intensity of U87MG-GFP-Luc cells in the mice following treatment with PBS, AS1411, Ced, AS1411+Ced, or AS1411-Ced. (**C**) Kaplan-Meier survival curves of mice after each treatment. (**D**) The bioluminescent images of orthotopic GBM mice after each treatment. n = 5 for each treatment group. (**E**) H&E staining of brain tissues from orthotopic GBM mice after each treatment. (**F**) IF staining of VEGFR2 (green) and Ki67 (red), and TUNEL staining (orange) of GBM sections in different treatment groups. Scale bars, 50 μm. **(G)** Quantification of fluorescence intensities of VEGFR2, Ki67, and TUNEL on GBM sections in different treatment groups. n = 3 for each treatment group. Graphs represented means ± SEM, and statistical significance was calculated by two-way ANOVA (B), log-rank (Mantel-Cox) test (C), and One-way ANOVA (G). \**P* < 0.05, \*\**P* < 0.01, \*\*\**P* < 0.001 and \*\*\*\**P* < 0.0001.

## DISCUSSION

The development of efficient and safe therapies for GBM remains an arduous journey as reflected by the largely negative outcome of most clinical trials^66^. This dilemma is primarily attributed to the presence of biological barriers, including the BBB and its modified form, the BTB, which restricts the targeted delivery of therapeutic agents to tumor sites^67^. The distinction between the BBB and BTB, however, remains a subject of ongoing debate^68^. While some studies suggest that the BTB retains certain features of the BBB, such as the expression of specific transporters and efflux pumps, others argue that it is markedly compromised in GBM, exhibiting increased vascular leakiness and aberrant signaling pathways^69^. This ongoing controversy underscores the complexity of the BTB and highlights the need for a more nuanced understanding of its biological and functional properties^11^. Resolving the discrepancies between BBB and BTB is critical for the development of targeted therapeutic strategies that can exploit the unique properties of the BTB to enhance drug delivery to GBM cells^70^.

EVs, loaded with diverse cargoes such as proteins, lipids, and nucleic acids^71^, play a vital role in maintaining GBM by reprogramming the adjacent brain microenvironment into a tumor-promoting system^72^. EVs are released from donor cells through the outward budding of the plasma membrane and can subsequently fuse with recipient cells^73^. Recent studies have emphasized the crucial involvement of GBM-derived EVs in facilitating crosstalk between GBM and brain capillary endothelial cells, driving the transformation of the BBB into BTB^16^. For example, numerous microRNAs carried within GBM-derived EVs have been reported to participate in this process by modulating the permeability and integrity of the BBB^74, 75^. In our study, we revealed that a shuttling protein, NCL, was transferred from GBM cells, rather than astrocytes, to the surface of capillary endothelial cells via EVs (**Fig. 7A and 7B**). Considering that NCL is overexpressed in GBM cells and can be partially translocated to their surface^24, 45^, we speculate that NCL carried by EVs likely originates from the surface of GBM cells and anchors on endothelial cells through direct membrane fusion. In addition to NCL, we demonstrated that GBM-derived EVs also carried several well-established pro-angiogenic factors, including FGF2, FGF13, and VEGFA, which have been implicated in GBM-derived EVs-mediated BTB formation^22^.

**Fig. 7.**
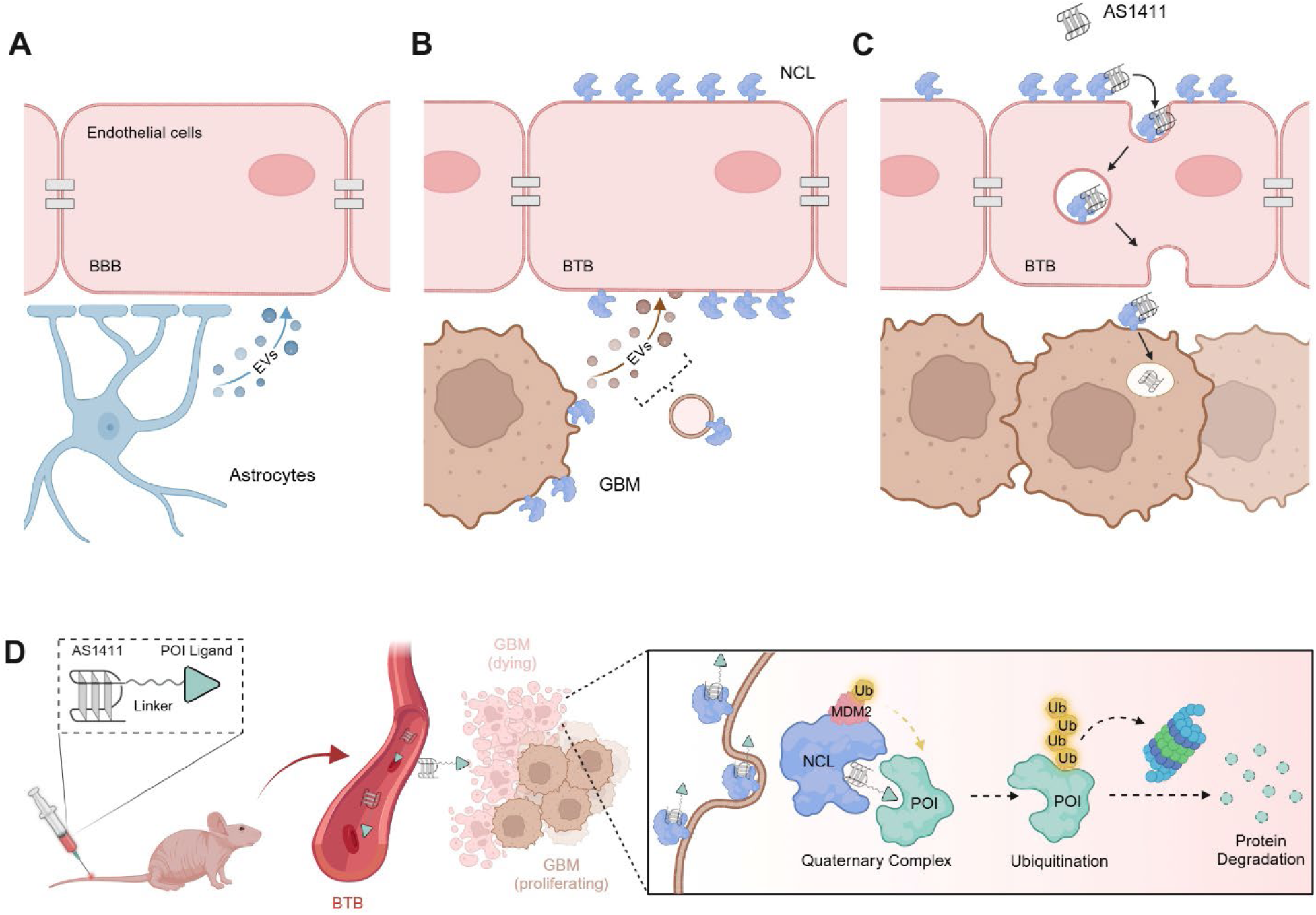
Targeting NCL for the development of the BTB-penetrating and GBM-specific PROTACs. (**A**) No transfer of NCL from astrocytes to the surface of brain capillary endothelial cells via EVs, maintaining the normal function of BBB. (**B**) EVs-mediated transfer of NCL from GBM cells to the surface of brain capillary endothelial cells, facilitating the transformation of BBB into BTB. (**C**) Transcytosis of AS1411 across the BTB in an NCL-dependent manner, enabling the targeting of GBM cells. (**D**) Development of BTB-penetrating and GBM-targeting PROTACs via conjugating AS1411 with ligands of POIs in GBM cells. Briefly, the AS1411-based PROTACs can transverse the BTB, selectively enter GBM cells, and induce the spatial proximity of POIs with the NCL-MDM2 complex. This will facilitate the ubiquitination and proteasomal degradation of POIs, leading to targeted therapy of GBM.

Regarding the role of NCL in BTB formation, we found that EVs-carried NCL transferred from GBM cells to the surface of capillary endothelial cells enhanced the survival and activities of BTB capillary endothelial cells. This observation aligns with previous reports showing that GBM-associated capillary endothelial cells exhibited reduced apoptosis and increased resistance to cytotoxic treatments^76, 77^, supporting the active participation of NCL in the pathological reorganization of the BTB, similar to the roles of VEGFA and FGFs^22^. Importantly, we demonstrated that normal astrocytes did not express NCL on their surface, which was consistent with the previous opinion that NCL is specifically expressed on the surface of cancer cells, but could not be translocated on the surface of normal cells^78^. Moreover, we found that astrocytes were incapable of delivering NCL to the surface of capillary endothelial cells via EVs, suggesting the unique contribution of NCL carried by GBM-derived EVs in reshaping the BBB into BTB. These findings elucidate the critical role of EVs-loaded NCL in mediating communication between GBM cells and capillary endothelial cells and provide a strong foundation for developing NCL-targeting strategies to enhance drug delivery across the BTB for targeted treatment of GBM.

RMT has been extensively explored for delivering drugs into the brain to access intracranial targets^79–82^. AS1411, a well-characterized aptamer, was initially developed as an anticancer agent for metastatic renal cell carcinoma but failed in phase 1/2 clinical trials due to limited efficacy^83^. Since then, AS1411 has been widely used as a delivery ligand for anticancer drugs because of its high affinity for NCL overexpressed on the cancer cell surface^84^. Despite its widespread use, there has been no evidence demonstrating that AS1411 can cross the BTB via the RMT pathway. In this study, we revealed that AS1411 could specifically traverse the BTB, but not the BBB, both *in vitro* and *in vivo*. This capacity of AS1411 is mediated by its recognition of NCL transferred from GBM cells to the surface of BTB capillary endothelial cells via EVs, which was supported by our findings that silencing NCL in GBM cells or blocking EV secretion from GBM cells diminished the BTB penetration of AS1411. It has been reported that cell surface NCL facilitates the internalization of AS1411 via clathrin-mediated endocytosis and macropinocytosis^85, 86^, and both internalizing pathways, as well as exocytosis, are critical steps involved in transcytosis^52^. We found that inhibition of either clathrin-mediated endocytosis, macropinocytosis, or exocytosis in capillary endothelial cells abolished the ability of AS1411 to cross the BTB, proving that the penetration of AS1411 across BTB is dependent on the RMT pathway. These findings highlight the critical role of NCL transferred from GBM cells to the surface of brain capillary endothelial cells via EVs in enabling AS1411 to specifically penetrate the BTB through transcytosis (**Fig. 7C**).

The development of PROTACs capable of effectively targeting intracranial lesions in GBM faces significant challenges^87^. Recently, a few PROTACs, such as C004091, XL01126, and ARV-102, have been developed with the ability to penetrate the BBB^88–90^. However, these PROTACs are designed for neurogenerative diseases, targeting tau and leucine-rich repeat kinase 2 (LRRK2), and their mechanisms for BBB-penetration remain unclear^88, 89^. To date, no PROTACs capable of crossing BTB to target POIs in GBM cells have been reported. As an alternative, researchers have proposed encapsulating PTOTACs within nanoparticles that can penetrate the BBB for targeted GBM therapy^91–93^. In our previous work, we repurposed AS1411 as a recruiter of E3 ligase MDM2 since it selectively entered cancer cells via targeting surface NCL and then recruited MDM2 using intracellular NCL as a molecular bridge^36^. By incorporating AS1411 into PROTACs, we successfully modulated the expression of various POIs, such as STAT3, c-Myc, c-MET, and VEGF165, achieving targeted therapy for numerous solid cancers^25, 27, 36^. In this study, we discovered that AS1411 also possesses the ability to penetrate BTB, in addition to its well-established cancer-targeting and MDM2-recruiting capacities, allowing us to design BTB-penetrating and GBM-targeting PROTACs for GBM (**Fig. 7D**). Using VEGFR2 and EGFR as examples of POIs^37–39^, we demonstrated that AS1411-based PROTACs induced their ubiquitination and proteasomal degradation in GBM cells rather than in normal astrocytes. This led to the inhibition of tumor cell growth and induction of apoptosis *in vitro*, as well as significant shrinkage of GBM tissues in mouse models. Moreover, these PROTACs exhibited minimal toxicity, likely due to their BTB-specific penetration and precise GBM targeting facilitated by AS1411. Notably, AS1411-based PROTACs offer significant advantages over the above-mentioned nanoparticle-loaded PROTACs, as they integrate the triple capabilities of BTB penetration, GBM targeting, and MDM2 recruitment into a single hetero-bifunctional molecule.

Despite the promising advancements demonstrated in our study, several limitations warrant further investigation to fully boost the efficacy of AS1411-based PROTACs. First, it has been well-established that both the length and the composition of the linkers play critical roles in determining the physicochemical properties and efficacy of PROTACs^94, 95^. AS1411-based PROTACs are hybrid molecules that consist of oligonucleotides and small molecules connected by chemical linkers. In this study, we only evaluated the effectiveness of AS1411-based PROTACs containing the most commonly used alkyl linker with a single length. However, we did not explore the broader impact of linker variability, such as different lengths or compositions, on the performance of AS1411-based PROTACs. Second, while our study provides a proof-of-concept demonstration of the potential of AS1411-based PROTACs for targeted therapy of GBM, several critical aspects remain to be fully elucidated, including the long-term stability, pharmacokinetics, and immunogenicity of these PROTACs *in vivo*. Future studies should address these challenges to optimize the therapeutic potential of AS1411-based PROTACs for the targeted treatment of GBM.

In conclusion, NCL transferred from GBM cells to the surface of brain capillary endothelial cells via EVs acts as a key contributor in driving BTB formation. The NCL-binding AS1411 is a versatile aptamer with three distinct and valuable capabilities: (1) penetrating the BTB, (2) selectively targeting GBM, and (3) recruiting the E3 ligase MDM2. By harnessing these properties, AS1411-based PROTACs emerge as a novel and highly promising strategy for the targeted treatment of GBM, offering a balance between therapeutic efficacy and safety.

## MATERIALS AND METHODS

### Cell culture

U87MG, U251, MCF 10A, HEK293T and bEnd.3 cell lines were obtained from the American Type Culture Collection (ATCC, USA). The U87MG-GFP-Luc cell lines were obtained by transduction using a viral vector expressing firefly luciferase and GFP protein (pCH-EF1a-GFP-T2A-Luc2-Ires-puro). The transduction was carried out as previously described^96^. After transduction, cells were treated with puromycin and sorted, and single-cell cloning was performed. Astrocytes were isolated according to the previous protocol and seeded on a T75 flask coated with poly-L-lysine (0.005 mg/ml, Beyotime, CHN)^97^. These cells were cultured in Dulbecco’s Modified Eagle’s Medium (DMEM; Corning, USA) supplemented with 10% FBS (Corning, USA) and 1% penicillin-streptomycin (Gibco, USA). Cell cultures were maintained at a standard condition of 37°C in a humidified environment containing 5% CO_2_.

### Plasmid constructs, lentivirus packaging, and transduction

To achieve overexpression of GFP-tagged NCL, cells were stably transfected with pCDH constructs (IGE BIO). To silence NCL gene expression, cells were stably transfected with Plko.1 construct (IGE BIO). All plasmids were confirmed by DNA sequencing. For lentivirus packaging and infection, HEK293T cells were seeded onto a 10 cm dish in a quantity of 1 × 10^6^. The transfection procedure was conducted after a duration of 24 h, utilizing 1 mg of target plasmids, 0.75 mg of psPAX2, and 0.25 mg of pMD2.G. The media of the culture was replaced at 6 h following the transfection process. The initial batch of material was collected at 48 h after transfection, without further changes to the medium. After medium refreshment, an additional collection of media took place at 72 h after the transfection. The two medium batches were combined and subjected to centrifugation at 1250 r/min for five min to remove any cell debris. The resulting supernatant was then kept in 1 mL portions at a temperature of −80 °C. 1 × 10^6^ U87MG cells were seeded onto a 6-well plate for each well and, after 24 h, were given 1 mL of the thawed lentivirus solution, which was gently stirred to ensure thorough mixing. The medium was substituted 24 h after the introduction of the virus. The process of antibiotic selection began 48 h after transfection, using a concentration of 2 mg/mL puromycin in the media. This selection continued until most cells died, leaving only the resistant cells, which finally attained confluency. Following that, the cells were prepped for extended preservation in liquid nitrogen.

### EV isolation

Cells were left in culture until they had reached 70–80% confluency, washed three times with PBS, and further cultured in FBS-free medium for 24 h before collection of the conditioned medium for EV purification^98^. EVs were isolated by differential ultracentrifugation as previously described with some modifications. Briefly, conditioned medium (100 mL; 10 × 10-cm dishes per replicate) was centrifuged at 300 × g for 10 min to pellet cells. Then, the supernatant was centrifuged for 30 min at 10,000 × g in a SW 28 rotor to pellet apoptotic bodies and cellular debris. The collected media was ultracentrifuged at 100,000 × g for 70 min to pellet EVs. Finally, the EV pellet was resuspended in PBS, carefully washed, and centrifuged at 100,000 × g for 70 min to collect the final EV pellets. All centrifugation steps were performed at 4 °C. The total EV protein concentrations were determined by the BCA Protein Assay kit (Thermo Fisher Scientific, USA). Proteins from isolated EVs were prepared for proteomics.

### Sample preparation for proteomics

Protein extracts from EVs or U87MG cells were prepared using the EasyPep Mini MS Sample Prep Kit (Thermo Fisher Scientific, USA)^99^. Initially, EVs or cells were lysed with the provided lysis buffer to extract proteins. A quantity of 100 µg of the protein extract was then transferred to a fresh microcentrifuge tube, and the volume was brought up to 100 µL using the same lysis buffer. Subsequent steps involved the addition of reduction and alkylation solutions to the protein sample. The mixture was gently mixed and subjected to incubation at 95°C for 10 min to facilitate the reduction and alkylation processes. Post-incubation, the sample was allowed to cool to room temperature. The reconstituted Trypsin/Lys-C Protease Mix was then introduced to the prepared protein sample, followed by incubation with shaking at 37°C for 2 h, to achieve protein digestion. Upon completion of the digestion, the digestion stop solution was added to terminate the enzymatic reaction. Peptides were then purified using a peptide clean-up column. The samples were dried via vacuum centrifugation and subsequently reconstituted in a 0.1% formic acid aqueous solution, preparing them for LC- MS/MS analysis.

### Nanoflow LC-MS/MS analysis

The Orbitrap Fusion mass spectrometer (Thermo Fisher Scientific, USA) was used in conjunction with an Easy-nLC 1000 ultrahigh-pressure liquid chromatography pump (Thermo Fisher Scientific, USA) for the LC- MS/MS analysis. Separation was achieved using a trap column and an analytic column with a spray tip, both filled with 3 µm/120 Å ReproSil-Pur C18 resins (Dr. Maisch GmbH, DE). The employed separation buffers were comprised of 0.1% formic acid in both water and acetonitrile. A fraction of the collected samples was initially introduced into the trap column with a 2 µL/min flow rate, and then, it was separated via the analytical column at a flow rate of 300 nl/min. The separation gradient was established as starting with 3% to 7% acetonitrile over 2 minutes, increasing to 22% acetonitrile over the next 50 minutes, then to 35% acetonitrile in 10 minutes, surging to 90% acetonitrile within 2 minutes, maintaining at 90% for 6 minutes, dropping back to 3% acetonitrile in 2 minutes, and finally stabilizing at 3% acetonitrile for a duration of 13 minutes. Full MS scans were performed in an Orbitrap mass analyzer over m/z range of 395-1205 with a mass resolution of 60,000. Data was processed and analyzed for DIA-Based proteomics using Spectronaut version 14.9 (Biognosys, CH)^100^.

### AS1411 transwell assay

For the *in vitro* BBB model, bEnd.3 were plated onto the apical side of transwell inserts (0.4 µm pores, 12 mm diameter) at 1 × 10^5^ cells/cm^2^ per well alone or in coculture with astrocytes, which were seeded on the basolateral side at 1 × 10^5^ cells/cm^2^ per well. For the *in vitro* BTB model, the U87MG cells were seeded into the basolateral chambers of the transwell plates at a density of 1 × 10^6^ cells/cm^2^ per well, and bEnd.3 were seeded onto the upper inserts of the transwell plates at a density of 1 × 10^5^ cells/cm^2^ per well. Medium was refreshed every 48 h, and cultures were maintained for 5 days until a continuous monolayer developed. Paracellular integrity was then verified by quantifying the flux of 70 kDa TMR-dextran; fluorescence was read at 555/580 nm (ex/em) on multi-mode microplate reader (PerkinElmer Enspire 2300, USA). After confirming barrier restriction, 1 µM AS1411 was introduced into the upper chamber. At designated intervals, the entire basolateral volume was collected, and AS1411 accumulation was determined via fluorescence^46–49^.

### Western blotting

Western blot analysis was conducted as previously described^101^. For the extraction of total protein, cells were lysed using the RIPA lysis buffer (Beyotime, CHN), and supernatants were collected. For membrane protein, extraction was performed by using the Membrane and Cytosol Protein Extraction Kit (Beyotime, CHN), following the manufacturer’s instructions. Then the protein levels were quantified and loaded onto an SDS- PAGE gel. The separated proteins were then transferred onto a PVDF membrane (Millipore, MA, USA) via the Bio-Rad Trans-Blot Turbo™ transfer apparatus (USA). Following blocking with 5% non-fat dry milk in TBST, the membrane was incubated overnight at 4°C with primary antibodies against VEGFR2 (1:1000, CST, USA), EGFR (1:1000, CST, USA), NCL (1:1000, Proteintech, CHN), MDM2 (1:1000, Proteintech, CHN), Na^+^/K^+^- ATPases (1:1000, Proteintech, CHN), and ACTIN (1:2000, Abclonal, CHN). Following this, the membrane was rinsed thrice using TBS-T and then incubated with suitable HRP-linked secondary antibodies for one hour at room temperature. Chemiluminescent detection was subsequently carried out using the enhanced chemiluminescence kit (Abclonal, CHN), and the blots were visualized using the Tanon Multi5200 chemiluminescence imaging system (Tanon, Multi5200, CHN).

### Membrane NCL detection via flow cytometry

Cells were enzymatically detached and washed three times with PBS. After fixation with 10% formalin, the cells were blocked with a blocking buffer for 30 min. The cells were then incubated with 488-conjugated NCL antibody (1:200, Proteintech, CHN) for 1 h at room temperature. After three times of washes with PBS, the cells were re-suspended in 500 μL PBS and analyzed by a BD flow cytometer (BiotreeDB, USA). Data acquired from the flow cytometer were analyzed with the FlowJo software (BiotreeDB, USA).

### IF staining

For IF staining of cells, 2 x 10^3^ cells were plated in cell culture slides. After fixation with 10% formalin in PBS, the cells were permeabilized with 0.1% Triton X-100 and blocked with 3% bovine serum albumin. The cells were then incubated with primary antibodies overnight at 4°C, followed by incubation with secondary antibodies conjugated to a fluorescent dye (Alexa Fluor 488, 555, or 594, Abcam, USA). After counterstaining the nuclei with DAPI (Beyotime, CHN), coverslips are mounted onto microscope slides using an anti-fade mounting medium. For IF staining of tissue sections, the sections were subjected to antigen retrieval and permeabilized with 0.2 % Triton X-100. Blocking was carried out with 5% BSA for an hour before incubating overnight at 4°C with primary antibodies. The sections were then rinsed and incubated with either anti-mouse or anti-rabbit Alexa Fluor 488, 555, or 594 secondary antibodies (1:200, Abcam, UK) for one hour at room temperature. Finally, the sections were counterstained with DAPI (Beyotime, CHN) and images were acquired using a confocal fluorescence microscope (Zeiss 980, DE).

### Cellular binding and uptake of AS1411 *in vitro*

U87MG, U251, astrocytes, or MCF 10A cells were seeded at a concentration of 1 × 10^6^ cells per well into 6- well plates. After overnight incubation in 2 mL of DMEM with 10% FBS for adherence, the cells were incubated with Cy5-labeled NC or AS1411. After washing three times with PBS, a flow cytometer (BD, FACSCanto SORP, USA) was used to determine the binding affinity of NC or AS1411 with the cells. Analysis was performed using the FlowJo software package. For confocal observations, cells were plated at a density of 2 × 10^3^ cells per well in cell culture slides. Following overnight incubation, the cells were treated with Cy5-labeled NC or AS1411. The cells were washed with PBS three times and fixed with 10% formalin. After counterstaining with DAPI (Beyotime, CHN), confocal images were acquired using a confocal fluorescence microscope (Zeiss 980, DE).

### 3D spheroid tumor model

The 3D tumor spheroids of U87MG were established according to the following steps. Briefly, U87MG cells were plated at a density of 2 x 10^3^ cells per well in a 96-well spheroid microplate (Corning, USA). After 4 days, the tumor spheroids were treated with Cy5-labeled NC or AS1411. Then, tumor spheroids were washed and fixed with 10% formalin. The permeability of Cy5-NC or Cy5-AS1411 into tumor spheroids was investigated by a confocal fluorescence microscope (Zeiss 980, DE).

### Detection of the G-quadruplexes

AS1411, AS1411-Ced, AS1411-Gef, and NC were diluted to 10 µM in cell-cultured medium and transferred to a black 96-well plate and then co-incubated with 1 μM NMM (MCE, USA) for 1 h. Fluorescence spectra from 550 to 750 nm were collected using a multi-mode microplate reader (PerkinElmer Enspire 2300, USA). Excitation was at 399 nm, and slit width was set to 1 nm for both excitation and emission.

### Gene silencing by siRNA

The siRNAs were purchased from OBiO (CHN). Cells were cultivated on 6-well plates up to 50-70% confluency for siRNA transfection. The siRNA at a final concentration of 160 nM was diluted in 150 μL Opti-MEM medium (Gibco, USA). 9 μL Lipofectamine RNAiMAX reagent (Invitrogen, USA) was diluted in 150 μL of Opti-MEM medium for 5 min at room temperature. The two solutions were mixed and incubated for 10 min at room temperature. After adding the siRNA-lipid complex to cells, cells were incubated for 2 days at 37 °C in a humidified environment containing 5% CO_2_.

### Cell viability assay

Cells were plated at a density of 2 x 10^3^ cells per well in a 96-well plate and allowed to adhere overnight. After treatment, cell viability was determined using the CCK-8 assays. This involved washing the cells with PBS, adding 10 µL of CCK-8 reagent to each well along with 100 µL of medium, and then incubating the plates for 1 h. Absorbance was measured at 450 nm using a multi-mode microplate reader (PerkinElmer, USA) at one-day intervals to monitor cell viability^102^.

### Apoptosis assay by flow cytometry

Cells were plated at a density of 1 x 10^6^ cells per well into 6-well plates and allowed to adhere overnight. After treatment, cells were enzymatically detached and washed twice with PBS to ensure purity. Subsequently, cells were stained utilizing an Annexin V-FITC Apoptosis Detection Kit (Beyotime, CHN) to label apoptotic cells. Flow cytometric analysis was carried out using a BD flow cytometer (BiotreeDB, USA) to quantify the proportion of apoptotic cells. Data acquired from the flow cytometer were analyzed with the FlowJo software (BiotreeDB, USA) to interpret the apoptotic profiles^101^.

### EdU cell proliferation assay

The proliferation of cells was detected using EdU cell proliferation assays according to the manufacturer’s instructions (Beyotime, CHN). Cells were plated at a density of 2 x 10^3^ cells in cell culture slides and incubated overnight for adherence. After treatment, a total of 500 µL EdU (10 µM) reagent was added to each well and incubated for 2 h to label the cells. After three times of washing with PBS, cells were fixed in a 10% formalin for 15 min, permeabilized with 0.3% Triton X-100 (Beyotime, CHN) for another 15 min, and then incubated with the click-reaction reagent for 30 min at room temperature in the dark environment. After counterstaining with DAPI (Beyotime, CHN), confocal images were acquired using a confocal fluorescence microscope (Zeiss 980, DE).

### 3D tumor spheroid invasion assay

Cells were plated at a density of 2 × 10^3^ per well in 96-well spheroid microplates (Corning, USA). After 4 days, 100 µl Matrigel Basement membrane matrix (Corning, USA) was added into the well. The plate was transferred to an incubator at 37 °C and allow the Matrigel Basement membrane matrix to solidify. After treatment, digital images were obtained using a confocal fluorescence microscope (Zeiss 980, DE) and quantitative analysis of the area of invasion was performed utilizing ImageJ software^62^.

### TUNEL assay

After different treatments, the cells and tissue sections were incubated with terminal deoxynucleotidyl transferase enzyme and Cy3-labeled dUTP mixture from the One Step TUNEL Apoptosis Assay Kit (Beyotime, CHN) for 1 h at 37 °C. Finally, the nucleus was counterstained by DAPI (Beyotime, China). Images were taken with a confocal fluorescence microscope (Zeiss, LSM980, DE).

### Tube formation assay

Tube formation assay was performed as previously described with some modifications^103^. In total, 100 μL Matrigel (Corning, USA) was added into 48-well plates and incubated at 37°C for 30 min. bEnd.3 cells were counted and diluted into a solution containing 2.5 × 10^5^ cells/mL. Mixed cell suspension (100 μL; 25,000 cells) was seeded on the Matrigel in 24-well plates and incubated for 12 h. bEnd.3 were stained with Calcein AM (Beyotime, CHN) and then imaged using a fluorescent inverted microscope equipped with a 485 nm excitation filter and a 520 nm emission filter. The vascular network was quantified using the ImageJ software (NIH, USA) with the angiogenesis analyzer plug-in.

### Transwell migration assay

Transwell migration assay was performed as previously described with some modifications^104^. Cells were seeded at a density of 5 × 10^4^ cells in 100 µL of serum-free medium into the upper chamber of a Transwell polycarbonate culture insert, 6.5 mm in diameter with an 8 µm pore size (BIOFIL, USA). These inserts were placed into 24-well plates containing 600 µL of medium supplemented with 10% FBS in the lower chamber. The assay plates were incubated for 12 h. Post-incubation, the Transwell inserts were carefully removed, and the upper chamber was cleared of non-migratory cells using a cotton swab. The cells that had migrated to the underside of the membrane were fixed with 10% formalin for 20 min, stained with 0.2% crystal violet for 30 min, and subsequently visualized under a Nikon Eclipse Ts2 microscope. To quantify the migratory cells, three distinct non-overlapping fields were counted using ImageJ software.

### IP and ubiquitination assay

The IP procedure was performed as previously described^105^. 5 × 10^4^ cells were lysed in lysis buffer (Thermo Scientific, USA) with a proteinase inhibitor cocktail. The lysate was centrifuged at 13,000 × g at 4 °C, and the resulting supernatant was incubated overnight at 4 °C with an anti-EGFR primary antibody (1:50, CST, USA), anti-VEGFR2 primary antibody (1:50, CST, USA), or anti-NCL primary antibody (1:100, CST, USA). Subsequently, the mixture was attached to Protein A/G Magnetic Beads (Thermo Scientific, USA) at room temperature for 2 h. The beads are then washed five times extensively to remove non-specifically bound proteins by using DynaMag™-2 Magnet (Invitrogen, USA). The immunoprecipitated proteins were then prepared in loading buffer, heated at 100 °C for 10 min, and subjected to SDS-PAGE and western blot analysis using primary antibodies against ubiquitin (1:2000, CST, USA), VEGFR2 (1:1000, CST, USA), EGFR (1:1000, CST, USA), NCL (1:1000, CST, USA), MDM2 (1:1000, CST, USA) and ACTIN (1:2000, CST, USA).

### Mice

Male BALB/c nude mice aged 6-8 weeks were obtained from GemPharmatech (CHN). The study conducted on these animals took place at the Experimental Animal Center of the Southern University of Science and Technology. This facility guarantees a controlled environment free from pathogens, offers unrestricted access to food and water, maintains a consistent temperature of 22 °C, and follows a 12-hour light/dark cycle. All animal handling protocols and experiments were approved by the Institutional Animal Care and Use Committees (IACUC) at Southern University of Science and Technology (SUSTech-JY202402003), guaranteeing adherence to recognized animal care criteria.

### Animal models of orthotopic GBM

An Orthotopic GBM mouse model was established through stereotactical inoculation of U87MG-GFP-Luc cells (3×10^5^ cells per mouse in 7 µL of PBS) into the brain (1.8 mm lateral and 1 mm anterior to the bregma; 3 mm of depth) of the mice that were anesthetized using 2.5% avertin (15 µL/g). After one week of tumor inoculation, mice were recruited into different treatment groups. The growth of the tumor was monitored by bioluminescence using an IVIS (PerkinElmer; IVIS Spectrum, USA), starting 10 min after the mice were injected with chloral hydrate (5% w/v) at a dose of 62.5 mg/kg combined with the luciferase substrate D-luciferin potassium (15 mg/mL in PBS) at 75 mg/kg^106^.

### Histological analysis

Brain or organs were dissected and fixed overnight in 10% formalin at 4°C. For the brain, a daily sucrose gradient (15 and then 30% sucrose in PBS) was used to wash out the formalin. Fixed brains were then frozen on dry ice and cryosectioned axially into 20 μm slices (Leica, CM1950, DE) for IF analysis. Organs were embedded in paraffin and sectioned to a thickness of 5 µm for staining with H&E. Images of the entire ankles were acquired using a NanoZoomer S60 Digital slide scanner (HAMAMATSU, JAN) and analyzed using NDP.view2 software (HAMAMATSU, JAN).

### Serum biochemical assays

Blood samples were collected from mice after treatment via hepatic portal vein puncture. Serum was collected via centrifugation of blood samples in 4000 × g at 4 °C for 10 min. Serum biochemical parameters, including ALT, AST, ALB, and ALP, were analyzed by an MS-480 Automatic Biochemistry Analyzer (Medicalsystem Biotechnology, CHN).

### Statistical analysis

Data analysis was conducted using GraphPad Prism software. For assessing differences among multiple independent groups, one-way ANOVA followed by a post-hoc test was employed. Comparisons between two independent groups were made using the Student’s t-test. For examining differences among groups categorized by two factors, two-way ANOVA with a subsequent post-hoc test was utilized. The statistical significance of the difference for survival analysis was analyzed by using the log-rank (Mantel-Cox) test. Data in the figures are represented as mean ± SEM, with levels of statistical significance denoted by \**P* < 0.05, \*\**P* < 0.01, \*\*\**P* < 0.001, and \*\*\*\**P* < 0.0001. Sample sizes for *in vivo* studies were established based on power analysis beforehand. Mice were assigned to groups in a random and blinded manner, and any mice in poor health prior to the start of the studies were not included.

## ACKNOWLEDGMENTS

The authors acknowledge the assistance of the Southern University of Science and Technology Core Research Facilities, the Microscope and Imaging Center, and the Experimental Animal Center of the Southern University of Science and Technology.

## AUTHOR CONTRIBUTIONS

Conceptualization: C.L. and A.L. Methodology: H.C. and J.Z. Software: H.C. Validation: H.C. and J.Z. Formal analysis: H.C. and J.Z. Investigation: H.C., J.Z., F.Q. and S.Y. Resources: F.Q. and Z.W. Data Curation: H.C. Writing original draft: H.C., J.Z., and C.L. Writing, review and editing: C.L., J.Z. and H.C. Visualization: H.C. and J.Z. Supervision: C.L. and A.L. Project administration: C.L. Funding acquisition: C.L. and A.L.

## DATE AVAILABILITY STATEMENT

The data that support the findings of this study are available from the corresponding author upon reasonable request.

## Supplementary Information

**Fig. S1.**
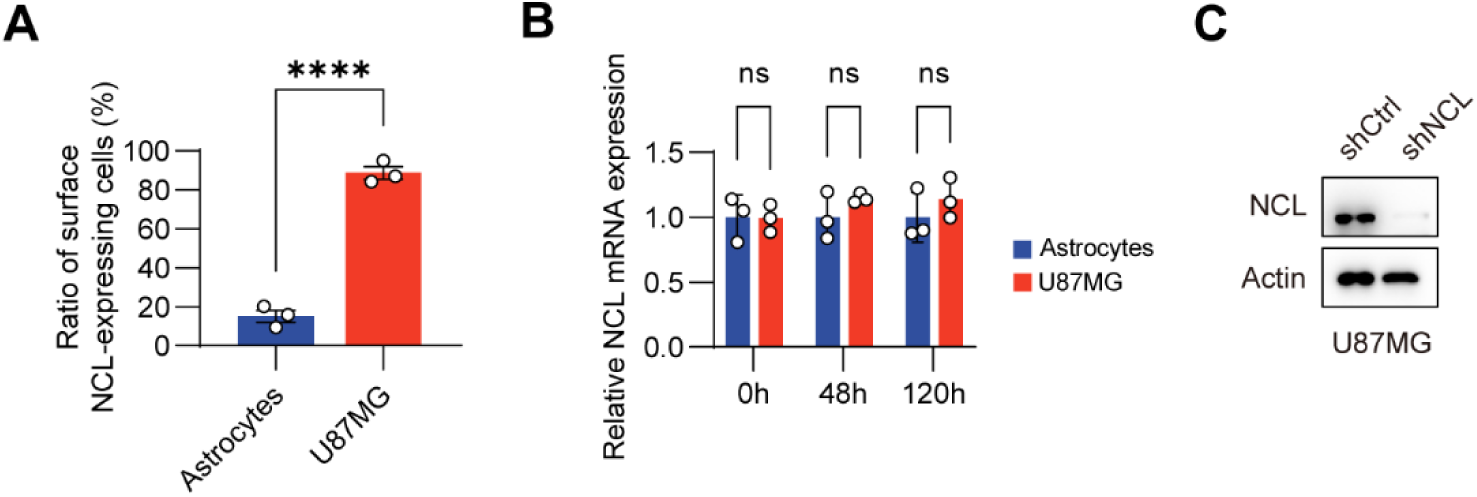
Expression of NCL on the surface of astrocytes and U87MG cells, or in bEnd.3 cells. (**A**) Flow cytometric analysis of surface NCL expression on astrocytes and U87MG cells using an anti-NCL antibody. (**B**) Relative mRNA expression of NCL in bEnd.3 cells after coculture with astrocytes or U87MG cells at 0, 48, and 120 h. (**C**) Protein expression of NCL in U87MG cells stably transfected with plasmids encoding shCtrl or shNCL. Experiments were performed in triplicate and repeated three times. Graphs represented means ± SEM, and statistical significance was calculated by Student’s *t*-test (A) and one-way ANOVA (B). \*\*\*\**P* < 0.0001. ns, no significance.

**Fig. S2.**
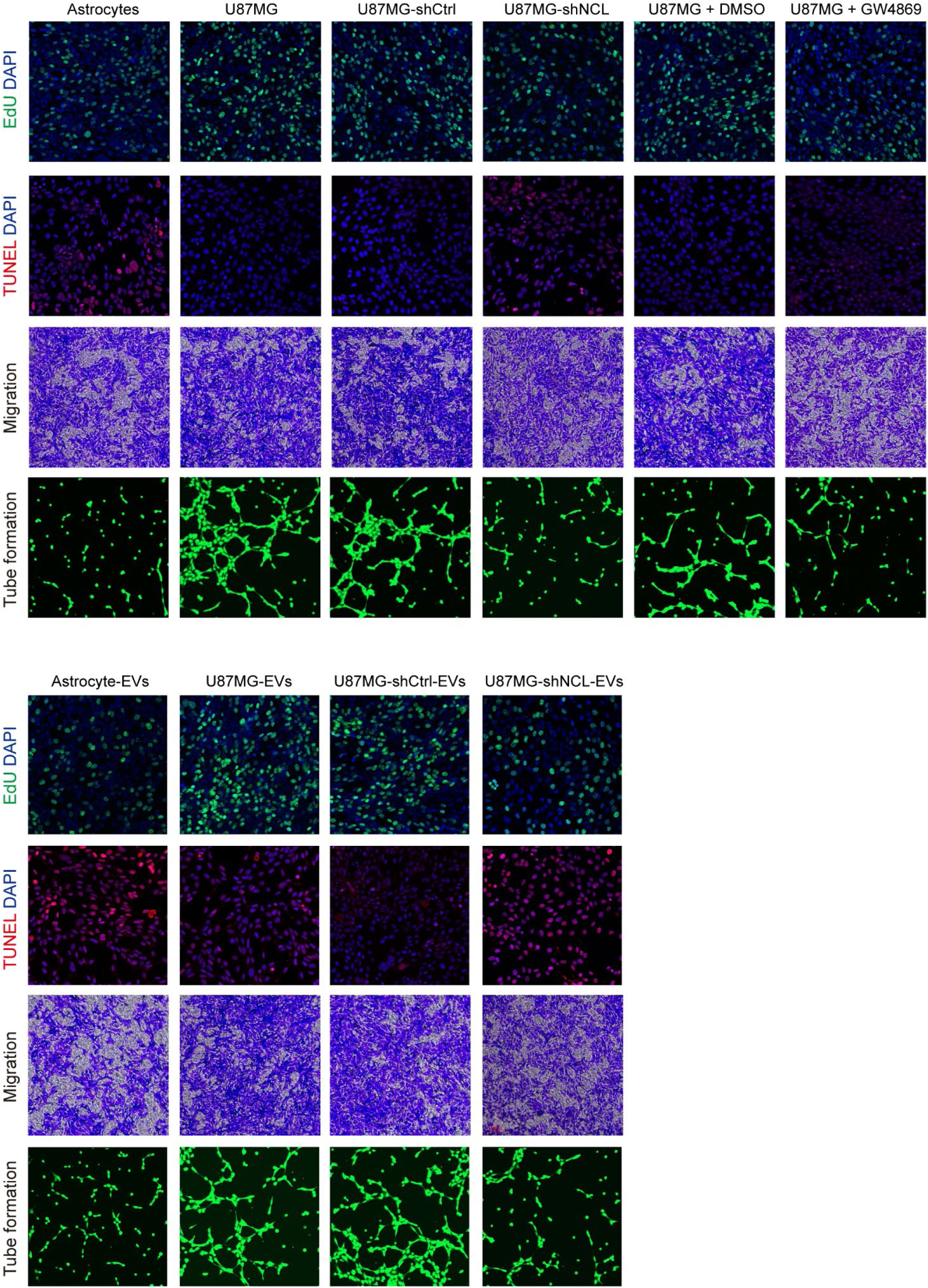
Proliferation, apoptosis, migration, and tube formation of bEnd.3 cells. (**A**) EdU and TUNEL staining, migration, and tube formation of bEnd.3 cells after coculture with astrocytes, untreated U87MG cells, or U87MG cells with or without different pretreatments. (**B**) EdU and TUNEL staining, migration, and tube formation of bEnd.3 cells after incubating bEnd.3 cells with EVs derived from astrocytes, untreated U87MG cells, or U87MG cells with or without different pretreatments. Scale bars for EdU and TUNEL staining, 100 μm. Scale bars for migration and tube formation, 200 μm. Experiments were performed in triplicate and repeated three times

**Fig. S3.**
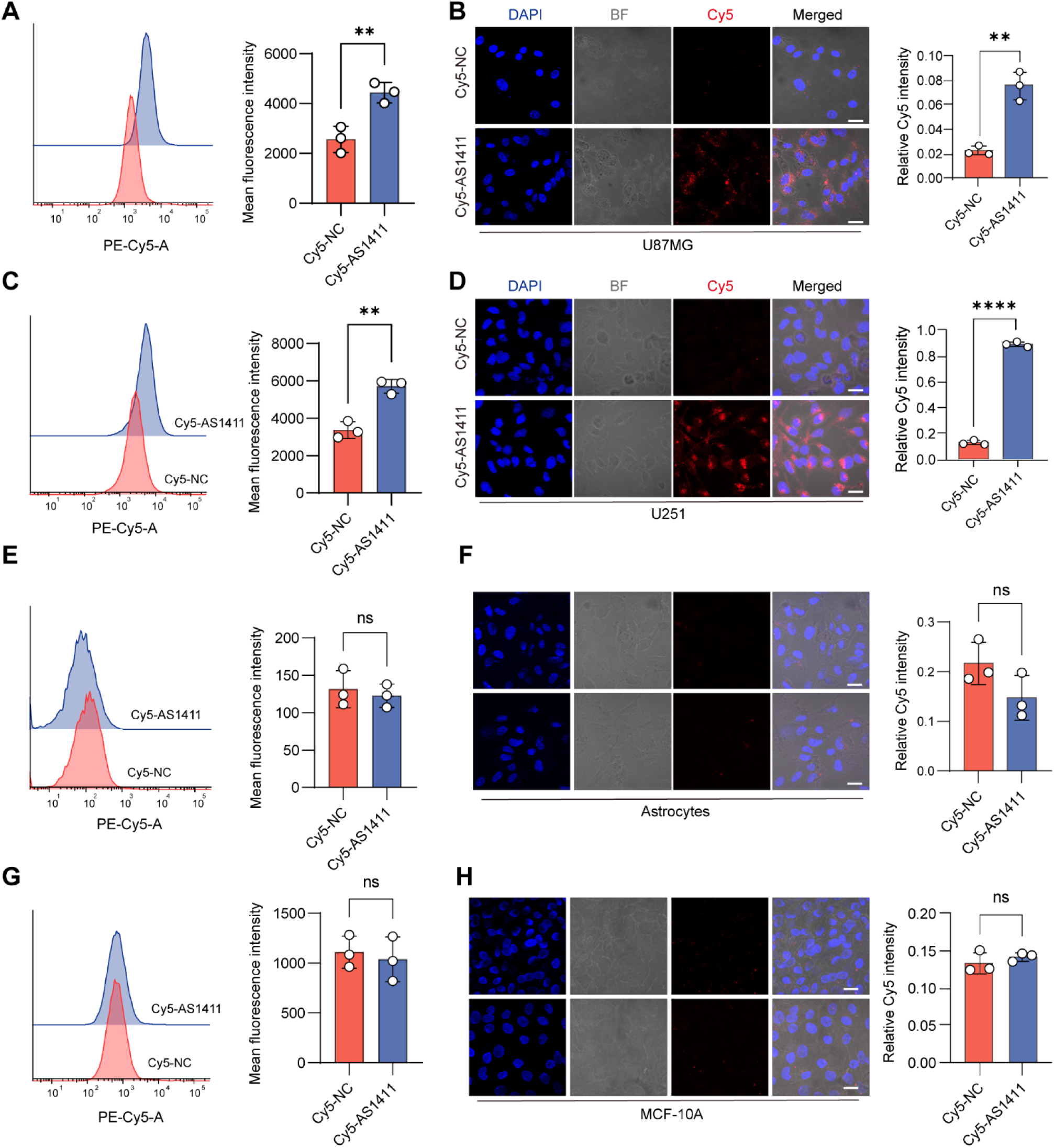
GBM-targeting and -internalizing ability of AS1411 *in vitro*. (**A**) Flow cytometric analysis of U87MG cells after incubation with 200 nM Cy5-labeled NC or AS1411 for 6 h. (**B**) Confocal analysis of U87MG cells after incubation with 200 nM Cy5-labeled NC or AS1411 for 6 h. Scale bars, 10 μm. (**C**) Flow cytometric analysis of U251 cells after incubation with 200 nM Cy5-labeled NC or AS1411 for 6 h. (**D**) Confocal analysis of U251 cells after incubation with 200 nM Cy5-labeled NC or AS1411 for 6 h. Scale bars, 10 μm. (**E**) Flow cytometric analysis of astrocytes after incubation with 200 nM Cy5-labeled NC or AS1411 for 6 h. (**F**) Confocal analysis of astrocytes after incubation with 200 nM Cy5-labeled NC or AS1411 for 6 h. (**G**) Flow cytometric analysis of the non-tumorigenic epithelial cell line MCF 10A after incubation with 200 nM Cy5-labeled NC or AS1411 for 6 h. (**H**) Confocal analysis of MCF 10A cells after incubation with 200 nM Cy5-labeled NC or AS1411 for 6 h. Scale bars, 10 μm. Graphs represented means ± SEM, and statistical significance was calculated by Student’s *t*-test (A, B, C, D, E, and F). \**P* < 0.05, \*\**P* < 0.01 and \*\*\*\**P* < 0.0001. ns, no significance.

**Fig. S4.**
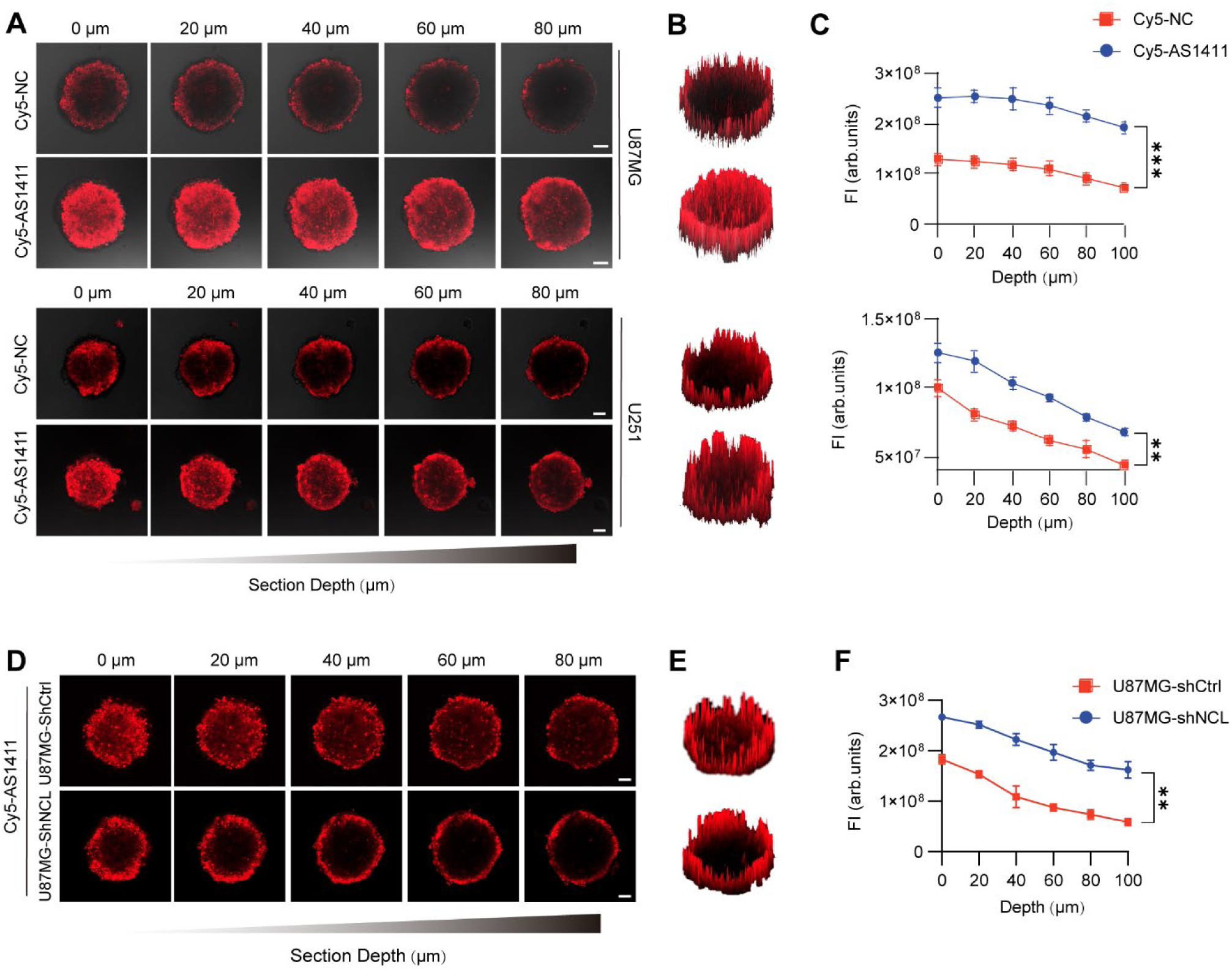
Internalization of AS1411 in 3D GBM spheroid models. (**A**) Confocal examination of 3D-cultured U87MG (upper) and U251 (bottom) cell spheroids treated with 200 nM Cy5-labeled NC or AS1411 for 6 h at scanning depths of 0, 20, 40, 60 and 80 μm. Scale bars, 50 μm. **(B)** 2.5-D reconstruction of U87MG (upper) and U251 (bottom) cell spheroids treated with 200 nM Cy5-labeled NC or AS1411 for 6 h at a scanning depth of 60 μm. **(C)** Fluorescence intensity of the central region versus Z-axis depth for Cy5-labeled NC or AS1411 in U87MG (upper) and U251 (bottom) cell spheroids. (**D**) Confocal examination of 3D-cultured shCtrl- or shNCL-transfected U87MG cell spheroids treated with 200 nM Cy5-labeled AS1411 for 6 h at scanning depths of 0, 20, 40, 60 and 80 μm. Scale bars, 50 μm. **(E)** 2.5-D reconstruction of shCtrl- or shNCL-transfected U87MG cell spheroids treated with 200 nM Cy5-labeled AS1411 for 6 h at a scanning depth of 60 μm. **(F)** Fluorescence intensity of the central region versus Z-axis depth for Cy5-labeled AS1411 in shCtrl- or shNCL-transfected U87MG cell spheroids. Experiments were performed in triplicate and repeated three times. Graphs represented means ± SEM, and statistical significance was calculated by two-way ANOVA (C and F). \*\**P* < 0.01 and \*\*\**P* < 0.001.

**Fig. S5.**
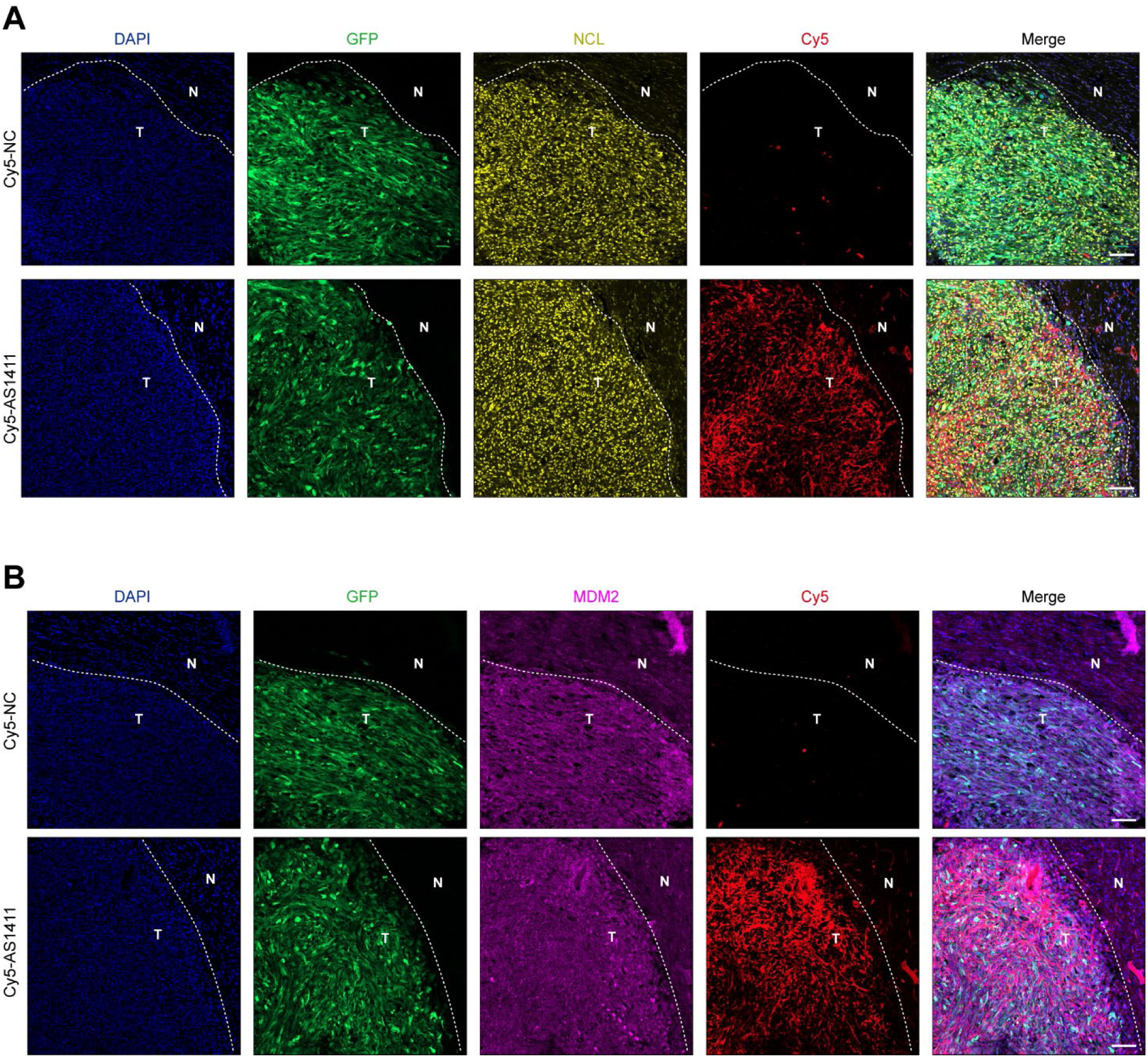
Co-localization of AS1411 with NCL and MDM2 *in vivo*. (**A**) Co-localization of Cy5-labeled NC or AS1411 (red) with NCL (yellow) in brain sections from orthotopic GBM mice inoculated with U87MG-GFP-Luc cells (green) after intravenous injections of 0.2 µmol/kg Cy5-labeled NC or AS1411 for 1 h. Scale bars, 100 μm. (**B**) Co-localization of Cy5-labeled NC or AS1411 (red) with MDM2 (purple) in brain sections from orthotopic GBM mice inoculated with U87MG-GFP-Luc cells (green) after intravenous injections of 0.2 µmol/kg Cy5-labeled NC or AS1411 for 1 h. Nuclei were stained with DAPI (blue). N, non-tumor region; T, tumor region. Scale bars, 100 μm. n = 3 for each group.

**Fig. S6.**
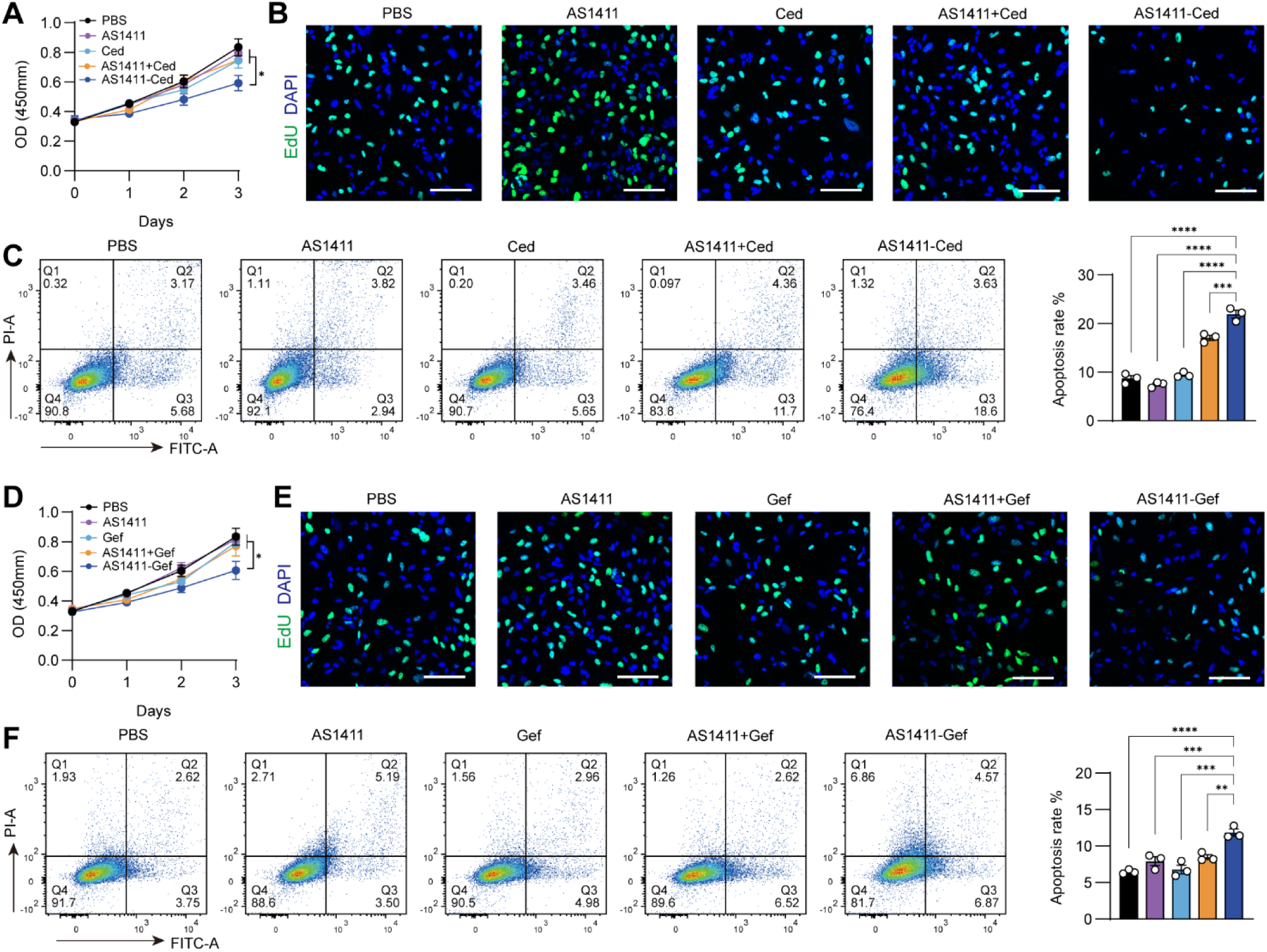
Anti-GBM activity of AS1411-based PROTACs *in vitro*. (**A**) CCK-8 assay for the viability of U251 cells after 3-day treatment with PBS, AS1411, Ced, AS1411+Ced, or AS1411-Ced at a concentration of 200 nM every day. (**B**) EdU staining of U251 cells after 3-day treatment with PBS, AS1411, Ced, AS1411+Ced, or AS1411-Ced at a concentration of 200 nM every day. Scale bars, 50 μm. (**C**) Flow cytometry analysis of apoptosis of U251 cells after 3-day treatment with PBS, AS1411, Ced, AS1411+Ced, or AS1411-Ced at a concentration of 200 nM every day. (**D**) CCK-8 assay for the viability of U251 cells after 3-day treatment with PBS, AS1411, Gef, AS1411+ Gef, or AS1411-Gef at a concentration of 200 nM every day. (**E**) EdU staining of U251 cells after 3-day treatment with PBS, AS1411, Gef, AS1411+ Gef, or AS1411-Gef at a concentration of 200 nM every day. Scale bars, 50 μm. (**F**) Flow cytometry analysis of apoptosis of U251 cells after 3-day treatment with PBS, AS1411, Gef, AS1411+ Gef, or AS1411-Gef at a concentration of 200 nM every day. Scale bars, 50 μm. Experiments were performed in triplicate and repeated three times. Graphs represented means ± SEM, and statistical significance was calculated by two-way ANOVA (A and D) and one-way ANOVA (C and F). \**P* < 0.05, \*\**P* < 0.01, \*\*\**P* < 0.001.

**Fig. S7.**
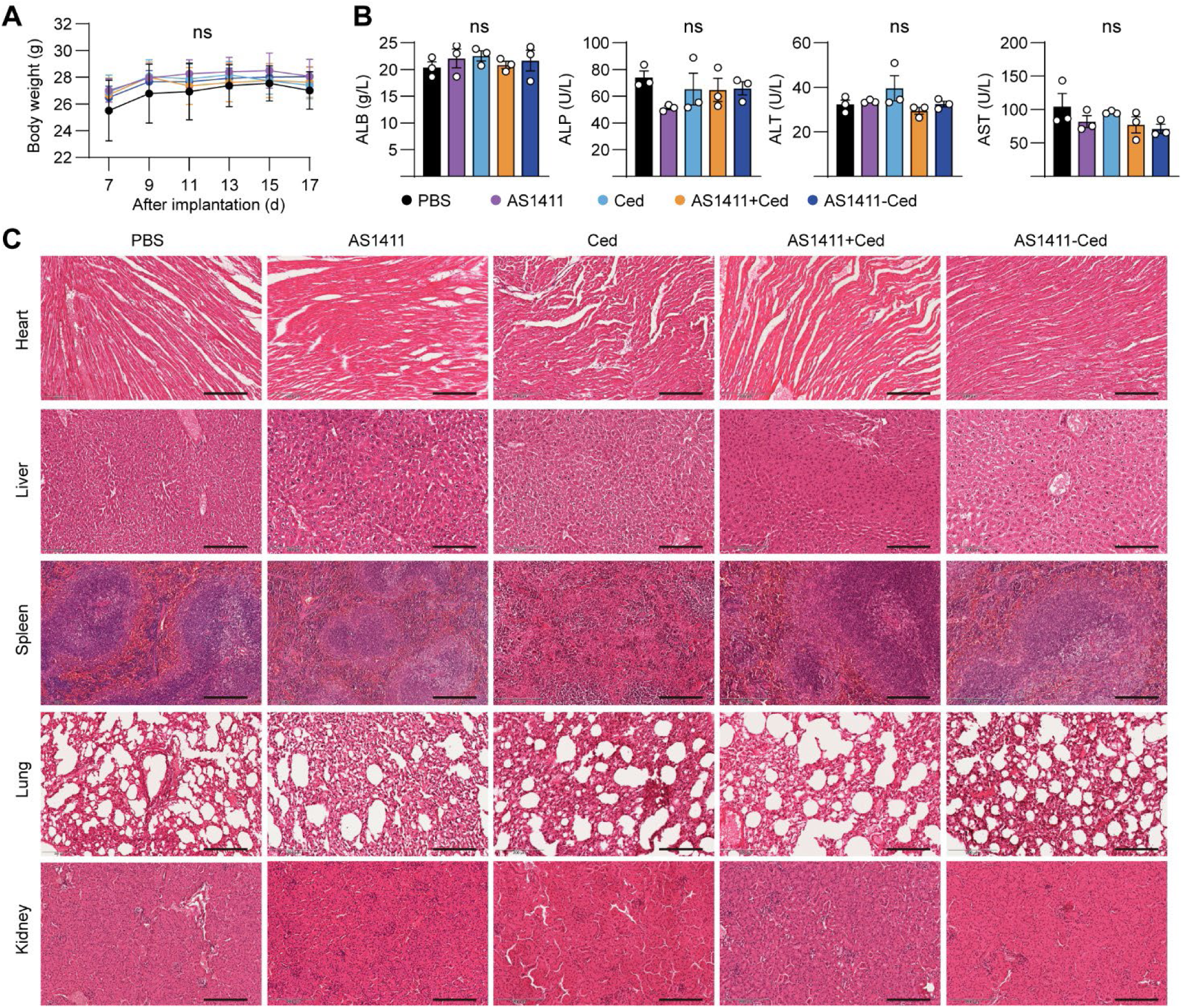
*In vivo* safety of AS1411-Ced. (**A**) Changes in body weight of the orthotopic GBM mice after each treatment. (**B**) Levels of ALB, ALT, AST, and ALP of the orthotopic GBM mice in each treatment group as determined by the automated hematology analyzer. (**C**) Histological assessments of non-tumor organs (heart, liver, spleen, lung, and kidney) of the orthotopic GBM mice in each treatment group by H&E staining. Scale bars, 200 μm. Data were presented as mean ± SEM. n = 3 for each group. ns, no significance.

**Fig. S8.**
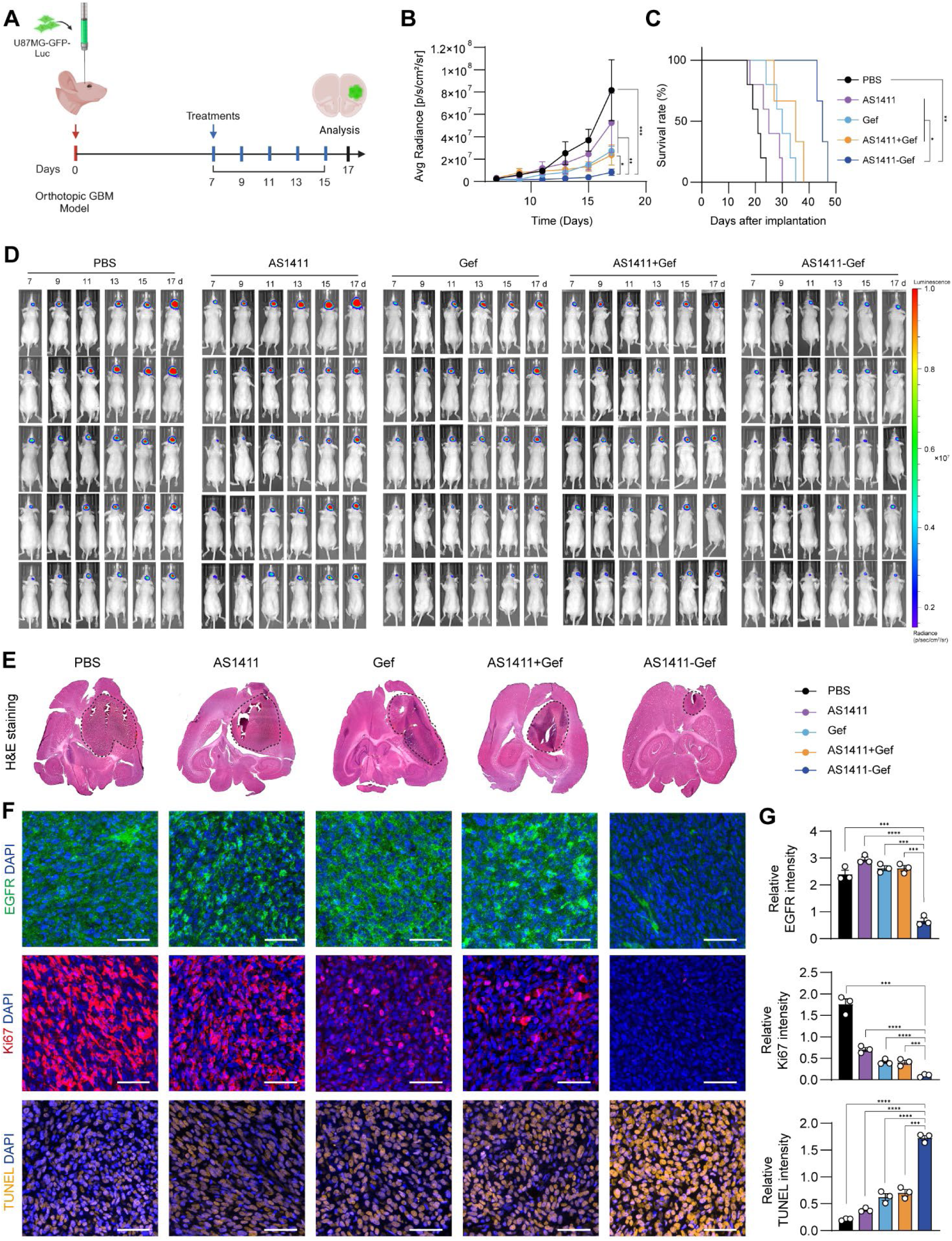
Anti-GBM efficacy of AS1411-Gef *in vivo*. (**A**) Schematic illustration for determining anti-GBM efficacy of AS1411-Gef *in vivo*. Briefly, U87MG-GFP-Luc cells were inoculated into the brains of BALB/c nude mice for 7 days. The orthotopic GBM mice were systemically administrated with PBS, AS1411, Gef, AS1411+Gef, or AS1411-Gef every other day for 8 days. The dose of AS1411-Gef was 6 μmol/kg. (**B**) The bioluminescent signal intensity of U87MG-GFP-Luc cells in the mice following treatment with PBS, AS1411, Gef, AS1411+Gef, or AS1411-Gef. (**C**) Kaplan-Meier survival curves of mice after each treatment. (**D**) The bioluminescent images of orthotopic GBM mice after each treatment. n = 5 for each treatment group. (**E**) H&E staining of brain tissues from orthotopic GBM mice after each treatment. (**F**) IF staining of EGFR (green) and Ki67 (red), and TUNEL staining (orange) of GBM sections in different treatment groups. Scale bars, 50 μm. **(G)** Quantification of fluorescence intensities of EGFR, Ki67, and TUNEL on GBM sections in different treatment groups. n = 3 for each treatment group. Graphs represented means ± SEM, and statistical significance was calculated by two-way ANOVA (B), log-rank (Mantel-Cox) test (C), and One-way ANOVA (G). \**P* < 0.05, \*\**P* < 0.01, \*\*\**P* < 0.001 and \*\*\*\**P* < 0.0001.

**Fig. S9.**
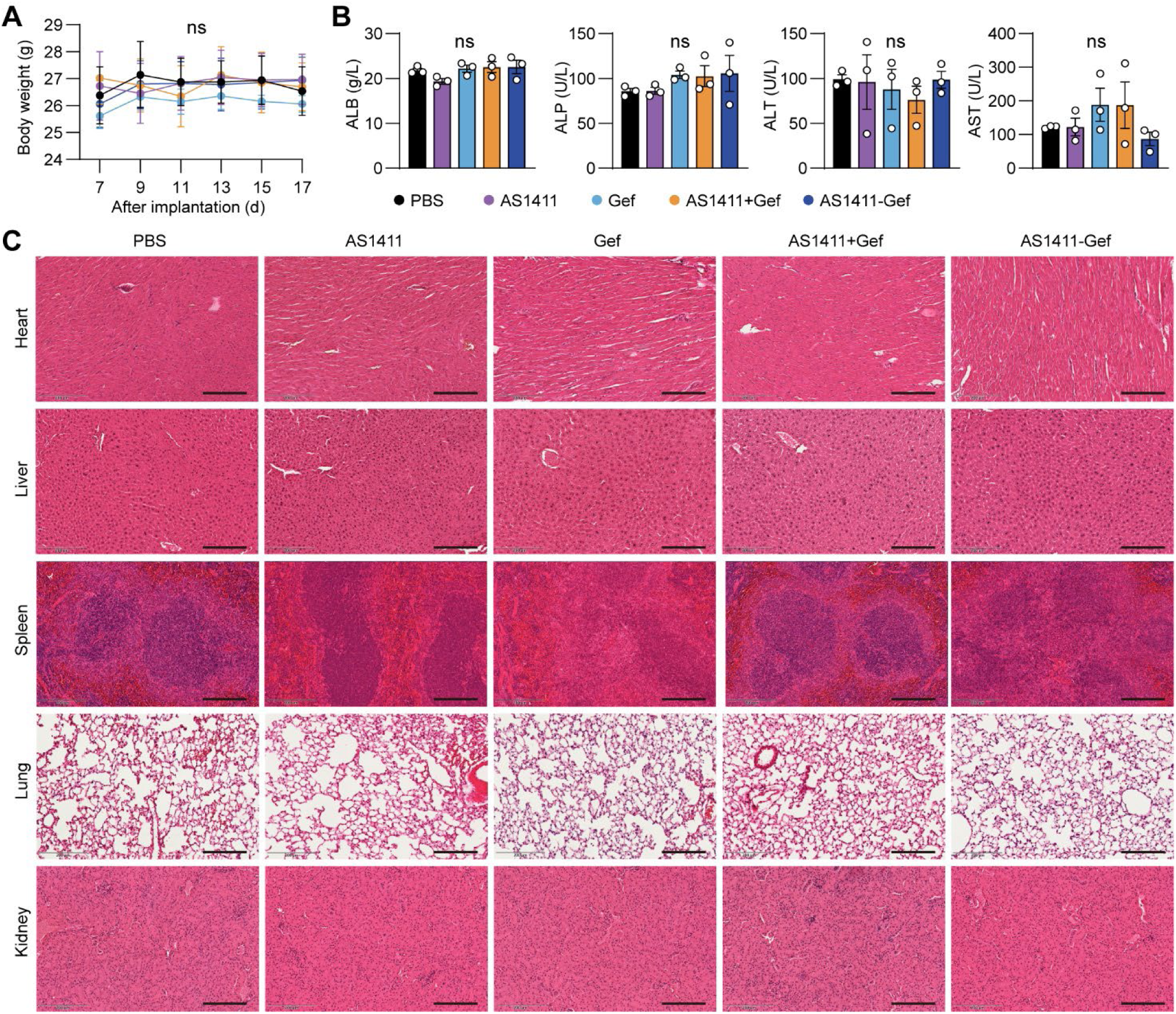
*In vivo* safety of AS1411-Gef. (**A**) Changes in body weight of the orthotopic GBM mice after each treatment. (**B**) Levels of ALB, ALT, AST, and ALP of the orthotopic GBM mice in each treatment group as determined by the automated hematology analyzer. (**C**) Histological assessments of non-tumor organs (heart, liver, spleen, lung, and kidney) of the orthotopic GBM mice in each treatment group by H&E staining. Scale bars, 200 μm. Data were presented as mean ± SEM. n = 3 for each group. ns, no significance.

**Table S1.** Biomarkers of EVs in proteomics.

|  | U87MG-EVs |  |  | Astrocyte-EVs |  |  |
| --- | --- | --- | --- | --- | --- | --- |
|  | Sample 1 | Sample 2 | Sample 3 | Sample 1 | Sample 2 | Sample 3 |
| CD63 | Positive | Positive | Positive | Positive | Positive | Positive |
| ARF6 | Positive | Positive | Positive | Positive | Positive | Positive |
| TSG101 | Positive | Positive | Positive | Positive | Positive | Positive |
| ANXA6 | Positive | Positive | Positive | Positive | Positive | Positive |
| GM130 | Negative | Negative | Negative | Negative | Negative | Negative |

## Supplementary Methods

### Scheme 1. Synthetic route for AS1411-Ced

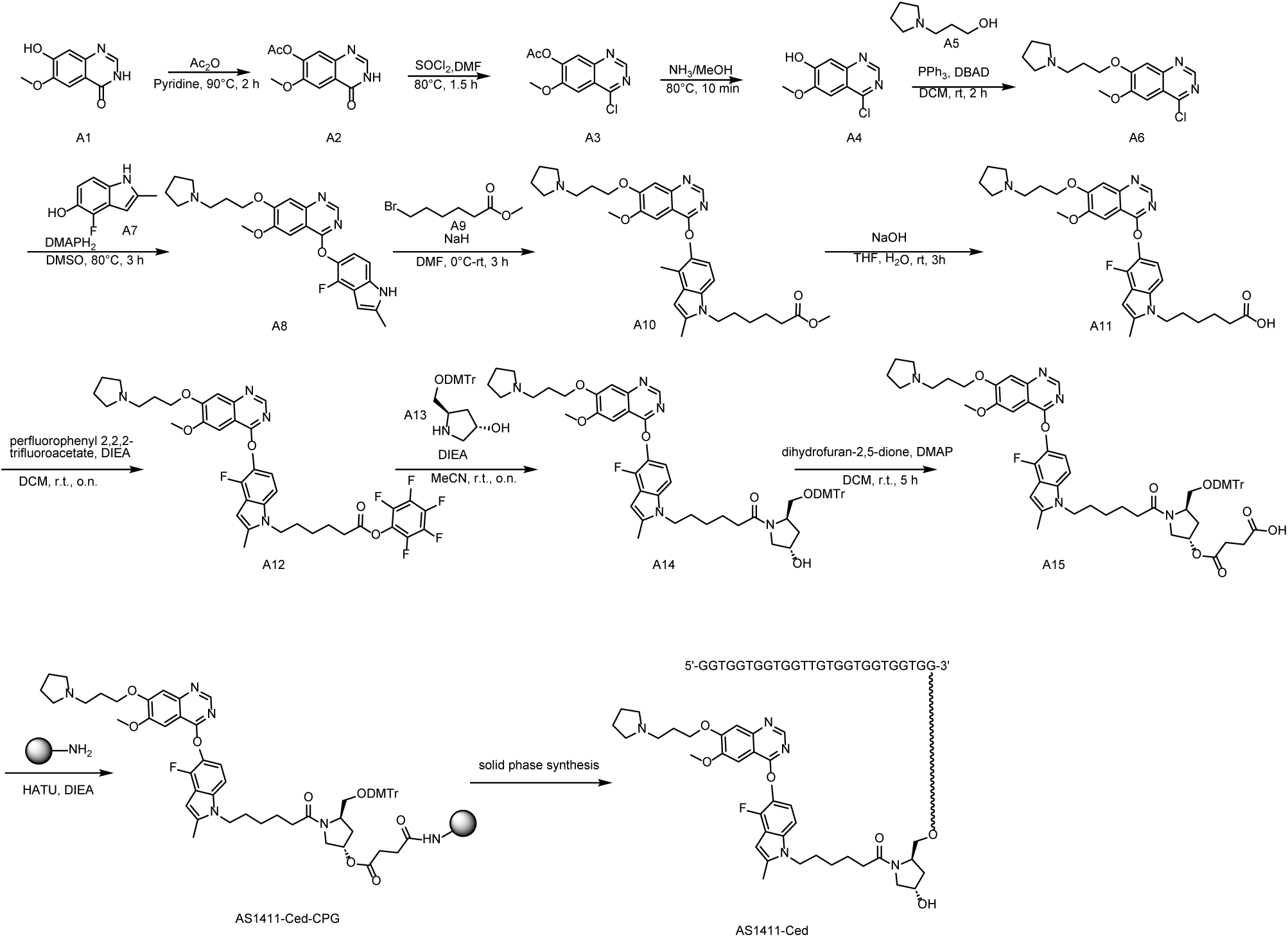

**(1) Synthesis of 6-methoxy-4-oxo-3,4-dihydroquinazolin-7-yl acetate (A2)**

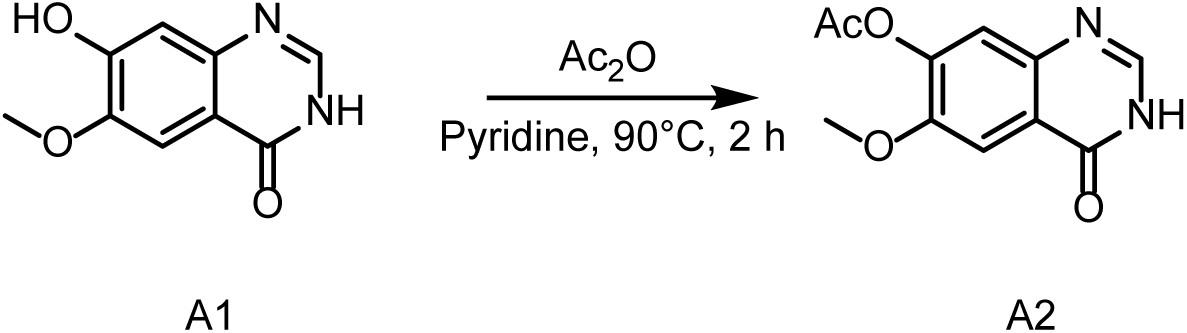

To a solution of **A1** (10.2 g, 156 mmol) in Pyridine (8.3 mL) was added Ac_2_O (33 mL, 1060.8 mmol) at 25 °C under N2 and the mixture was stirred at 90 °C for 2 hours. LCMS showed the reaction was complete. The reaction mixture was triturated with water (500 mL) at 25 °C for 10 min, filtered. The filter cake was dried to give **A2** (10.2 g, 83.69%) as a white solid.

**LCMS:** *m/z* = 235.2 [M+H]^+^, *t*_R_ = 1.024 min. **Purity:** 84.47% (254 nm).

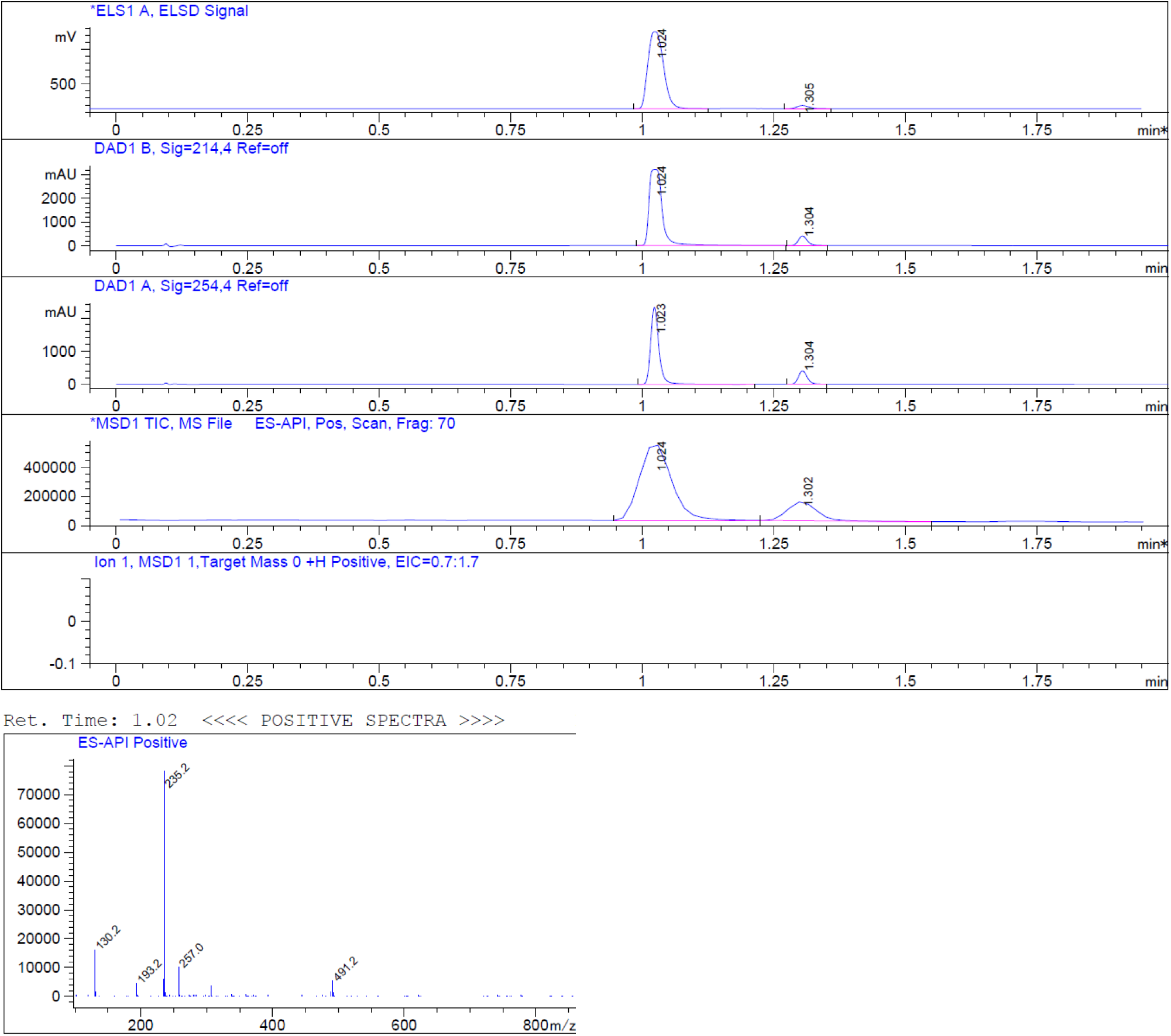

**(2) Synthesis of 4-chloro-6-methoxyquinazolin-7-yl acetate (A3)**

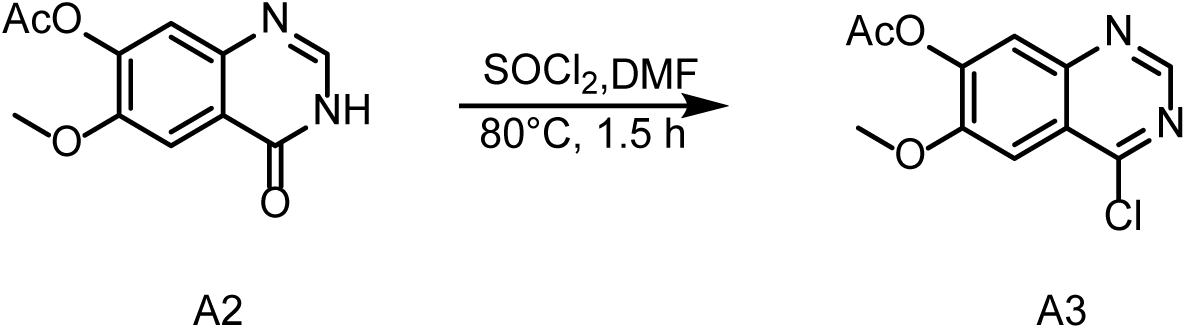

To a solution of **A2** (10.2 g, 43.55 mmol, 1.0 eq) in SOCl_2_ (100 mL) was added DMF (3 mL) at 25 °C under N_2_. Then the mixture was stirred at 80 °C for 2 hours. LCMS showed the reaction was complete. The reaction mixture was filtered and concentrated under reduced pressure to give the **A3** (10.5 g, crude) as a yellow solid. **LCMS:** *m/z* = 253.0[M+H]^+^, *t*_R_ = 1.340 min. **Purity:** crude

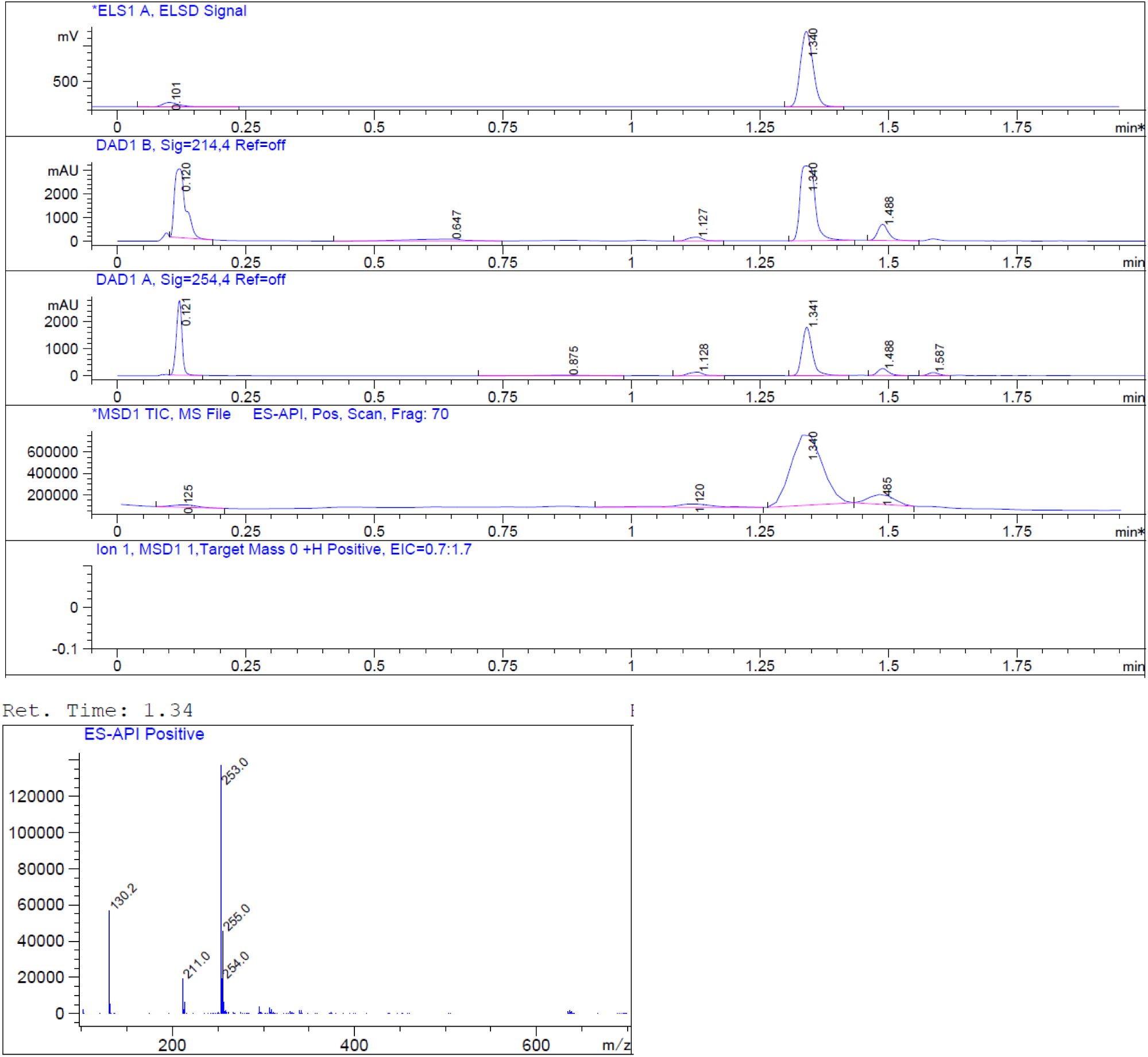

**(3) Synthesis of 4-chloro-6-methoxy-7-(3-(pyrrolidin-1-yl)propoxy)quinazoline (A6)**

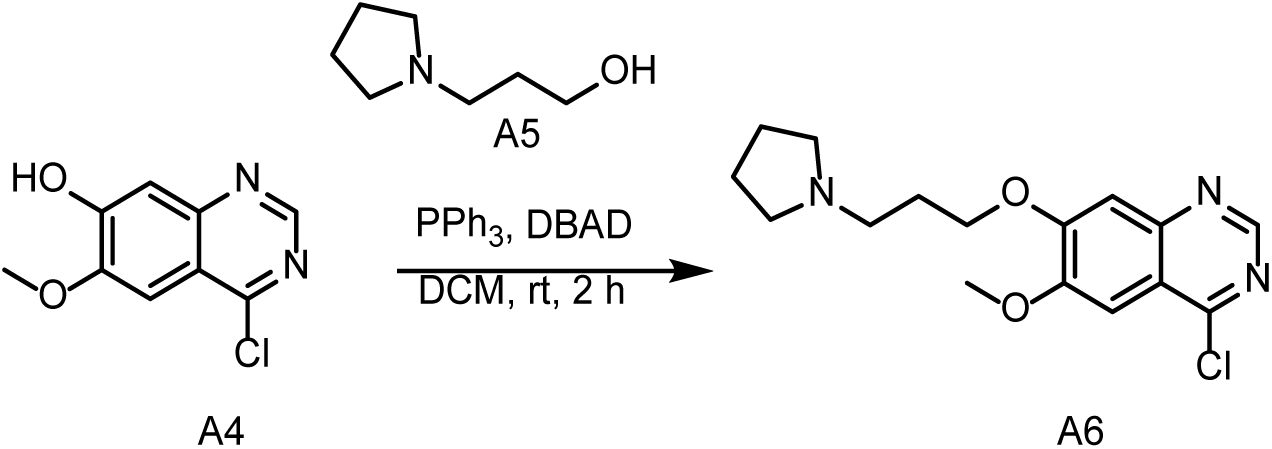

To a solution of **A4** (4.3 g, 20.43 mmol), **A5** (2.77 g, 21.44 mmol) and PPh_3_ (6.9 g, 26.55 mmol) in DCM (70 mL) was added DBAD (1.84 mg, 6.18 mmol) at 0 °C under N_2_. Then the mixture was stirred at 25 °C for 2 hours. LCMS showed the reaction was complete. The mixture was concentrated. The residue was quenched with water (200 mL) and extracted with EA (200 mL×2). The organic layers were washed with brine (200 mL), dried over anhydrous Na_2_SO_4_, filtered and concentrated in vacuo. The crude product was purified by column chromatography on silica gel (EA in PE=0 to 100%) to give **A6** (4.5 g, 63.8%) as a yellow solid.

**LCMS:** *m/z* = 322.2[M+H]^+^, *t*_R_ = 1.085 min. **Purity:** 100% (254 nm).

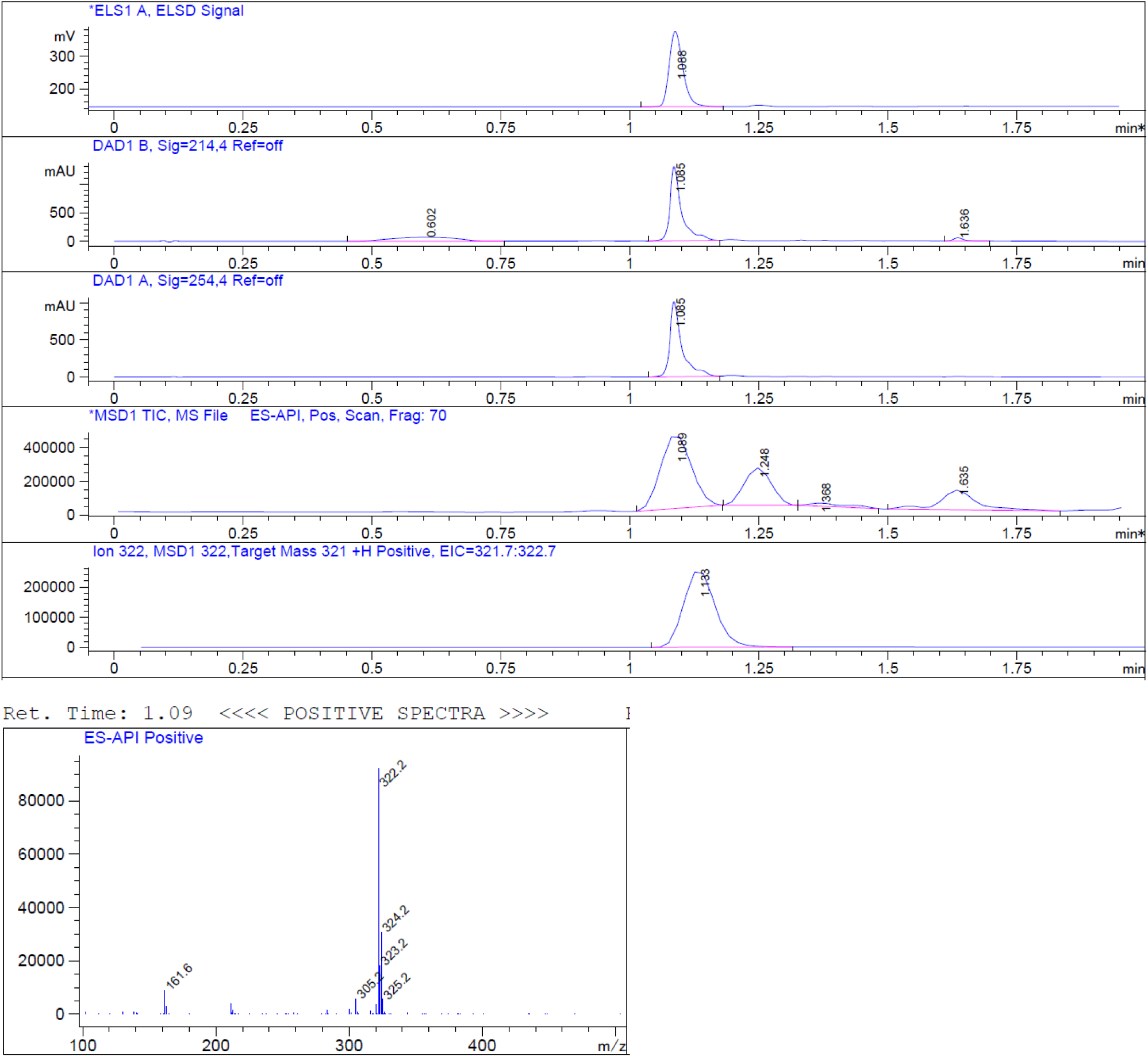

**(4) Synthesis of Synthesis of 4-((4-fluoro-2-methyl-1H-indol-5-yl)oxy)-6-methoxy-7-(3-(pyrrolidin-1-yl)propoxy)quinazoline (A8)**

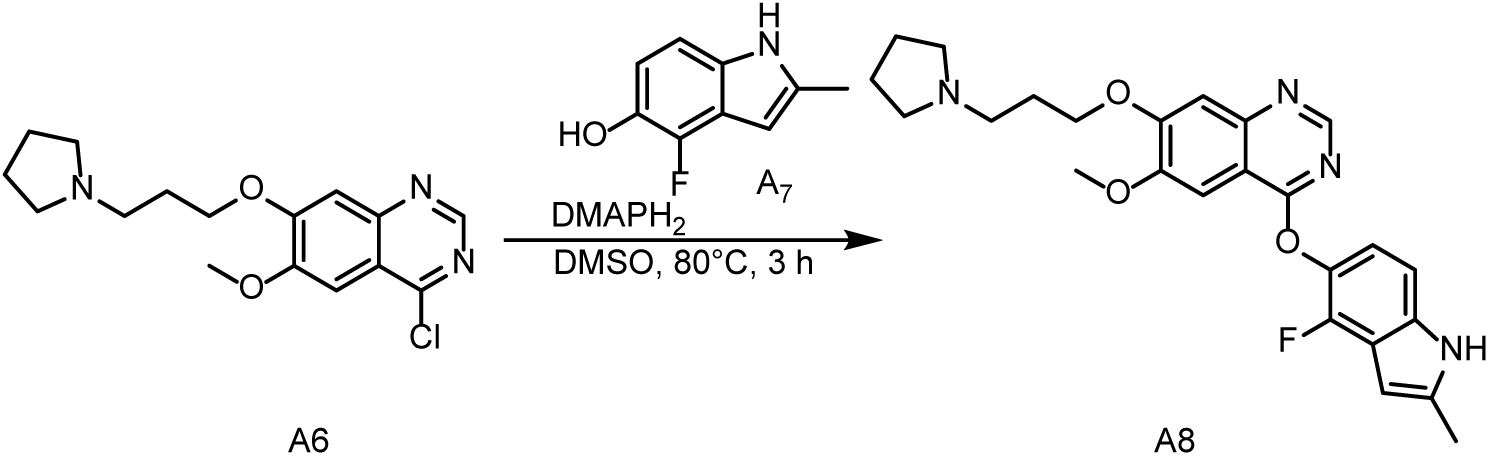

To a solution of **A6** (4.5 g, 14.29 mmol) in DMSO (50 mL) was added **A7** (2.6 g, 15.72 mmol,) and DMAP (3.9 g, 28.58 mmol) at 25 °C under N_2_. The mixture was stirred at 80 °C for 3 hours. LCMS showed the reaction was complete. The mixture was concentrated and the residue was quenched with water (200 mL) and extracted with EA (200 mL×2). The organic layers were washed with brine (200 mL × 1), dried over anhydrous Na_2_SO_4_, filtered and concentrated in vacuo. The crude product was purified by column chromatography on silica gel (EA in PE=0 to 100%) to give the **A8** (2 g, crude) as a yellow solid.

**LCMS:** *m/z* = 451.4[M+H]^+^, *t*_R_ = 1.085 min. **Purity:** 88.20% (254 nm).

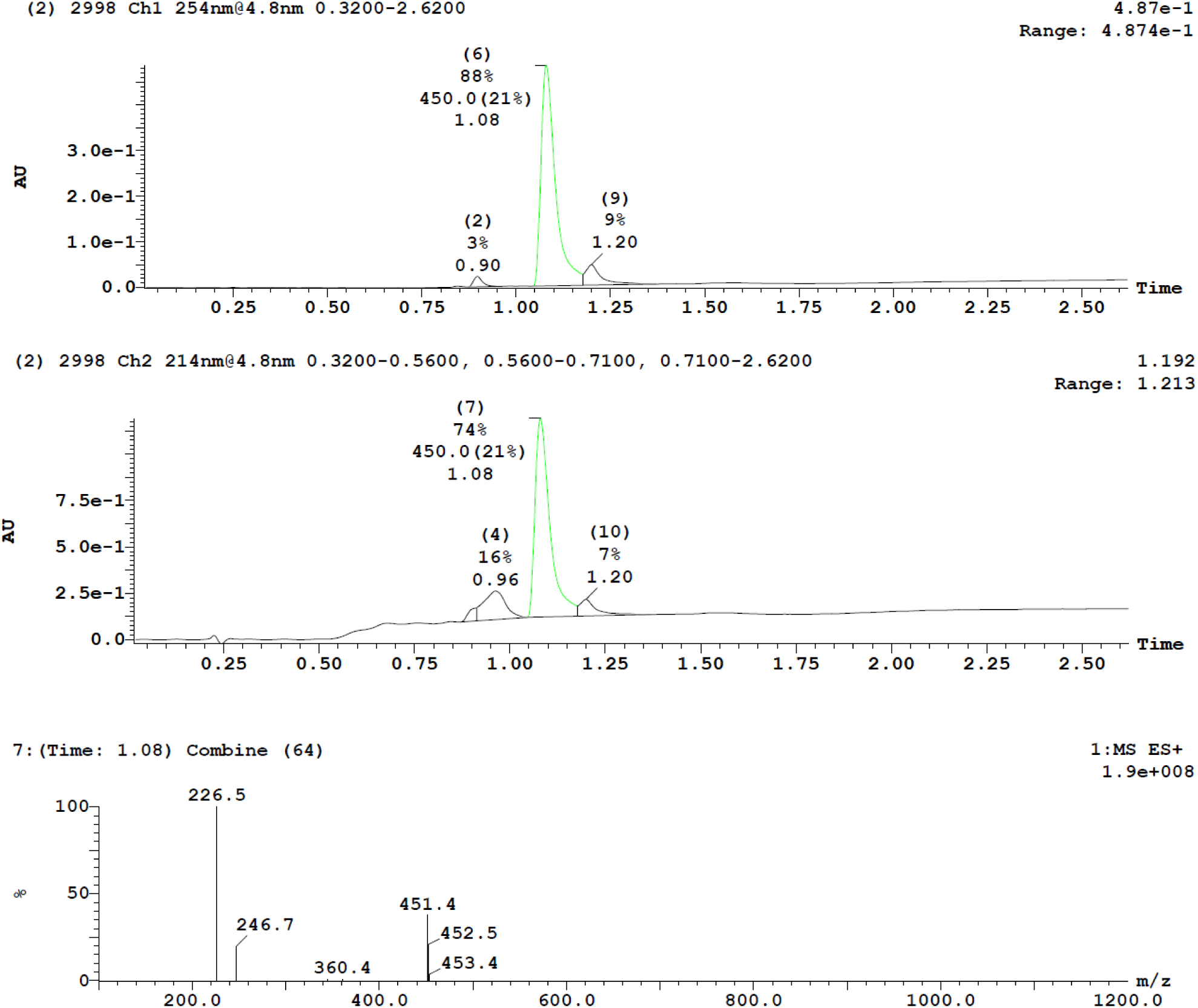

**(5) Synthesis of Synthesis of methyl 6-(5-((6-methoxy-7-(3-(pyrrolidin-1-yl)propoxy)quinazolin-4-yl)oxy)-2,4-dimethyl-1H-indol-1-yl)hexanoate (A10)**

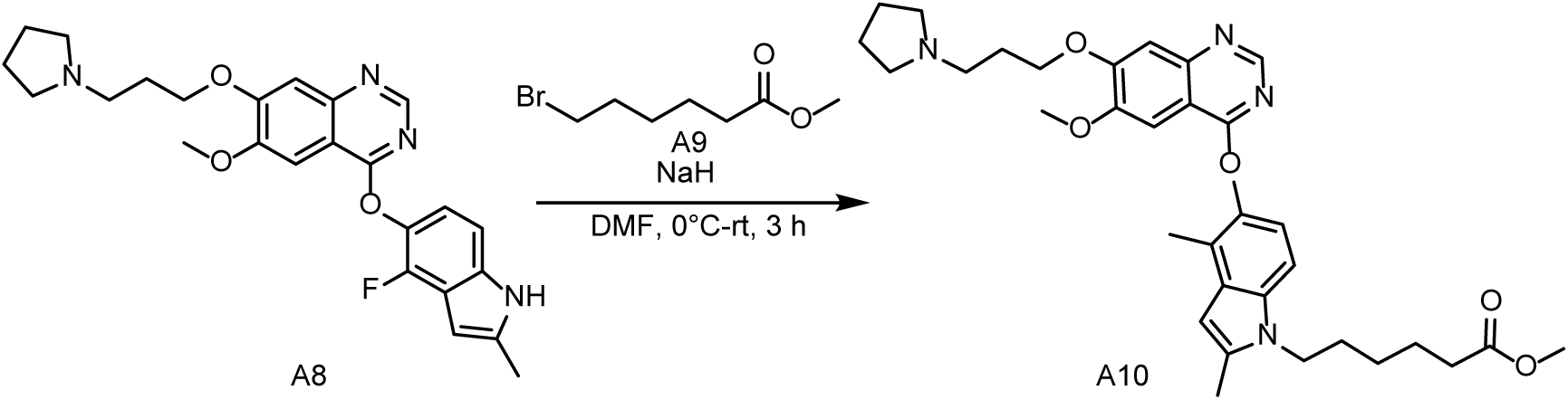

To a solution of **A8** (2 g, 4.44 mmol, 1.0 eq) in DMF (20 mL) was added NaH (266 mg, 6.66 mmol, 60% dispersion in mineral oil) at 0 °C under N_2_. After 30 min, **A9** (1.4 g, 6.66 mmol) was added and the mixture was stirred at 25 °C for 3 hours. LCMS showed the reaction was complete. The mixture was concentrated. The residue was quenched with water (200 mL) and extracted with EA (200 mL×2). The organic layers were washed with brine (200 mL), dried over anhydrous Na_2_SO_4_, filtered and concentrated in vacuo. The crude product was purified by column chromatography on silica gel (EA in PE=0∼100%) to give **A10** (3.3 g, crude) as a yellow solid.

**LCMS:** *m/z* = 579.4[M+H]^+^, *t*_R_ = 1.440 min. **Purity:** 74.40% (254 nm).

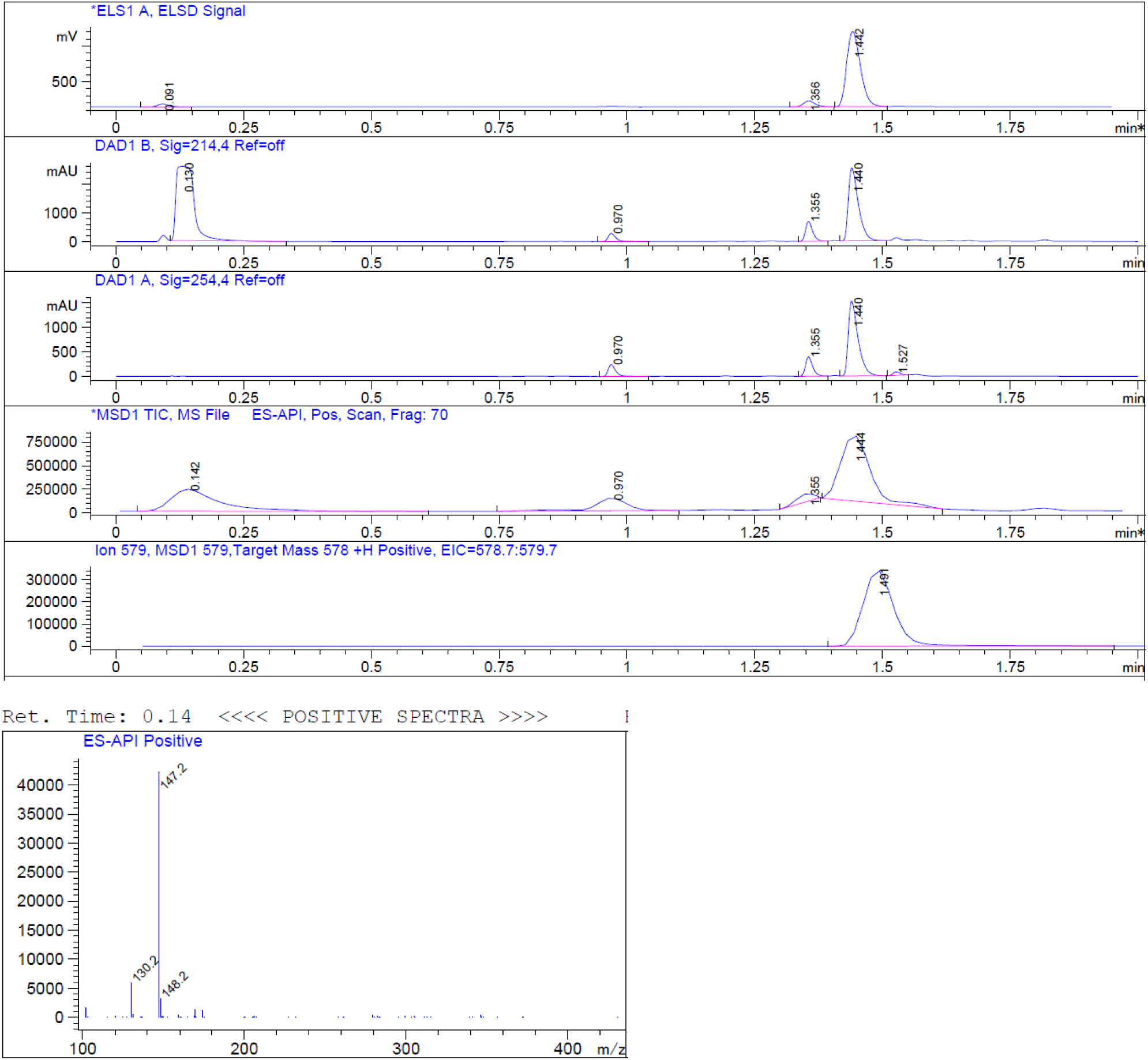

**(6) Synthesis of Synthesis of 6-(4-fluoro-5-((6-methoxy-7-(3-(pyrrolidin-1-yl)propoxy)quinazolin-4-yl)oxy)-2-methyl-1H-indol-1-yl)hexanoic acid (A11)**

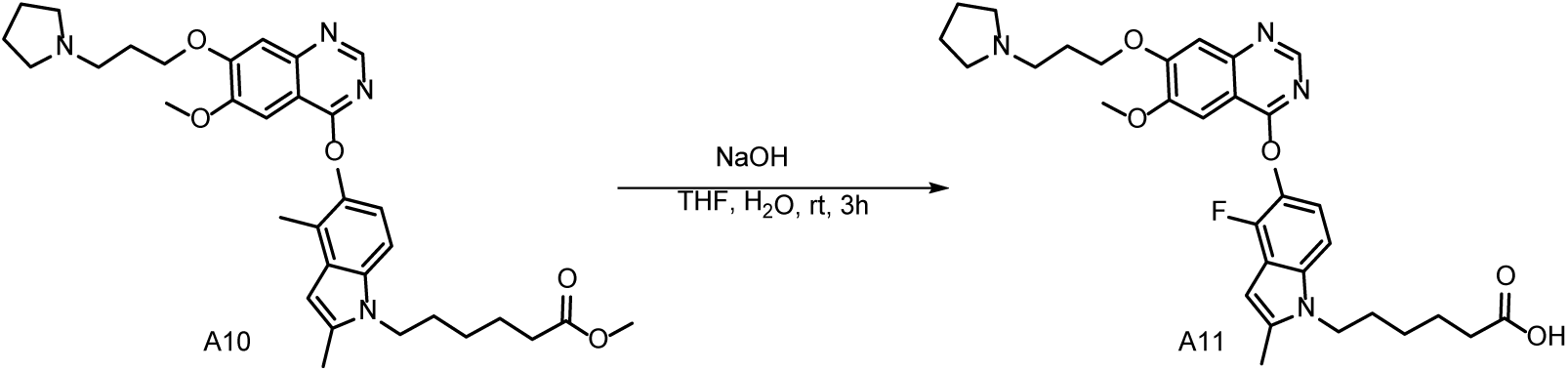

To a solution of **A10** (3.3 g, crude) in THF (20 mL) and H_2_O (10 mL) was added NaOH (230 mg, 5.8 mmol) at 0 °C under N_2_. The mixture was stirred at 25 °C for 3 hours. LCMS showed the reaction was complete. The mixture was concentrated. The residue was extracted with EA (200 mL×2). The organic layers were washed with brine (100 mL), dried over anhydrous Na_2_SO_4_, filtered and concentrated in vacuo. The crude product was purified by column chromatography on silica gel (EA in PE=0 to 100%) to give **A11** (1 g, 31.06%) as a yellow solid.

**LCMS:** *m/z* = 565.2[M+H]^+^, *t*_R_ = 1.362 min. **Purity:** 100% (254 nm).

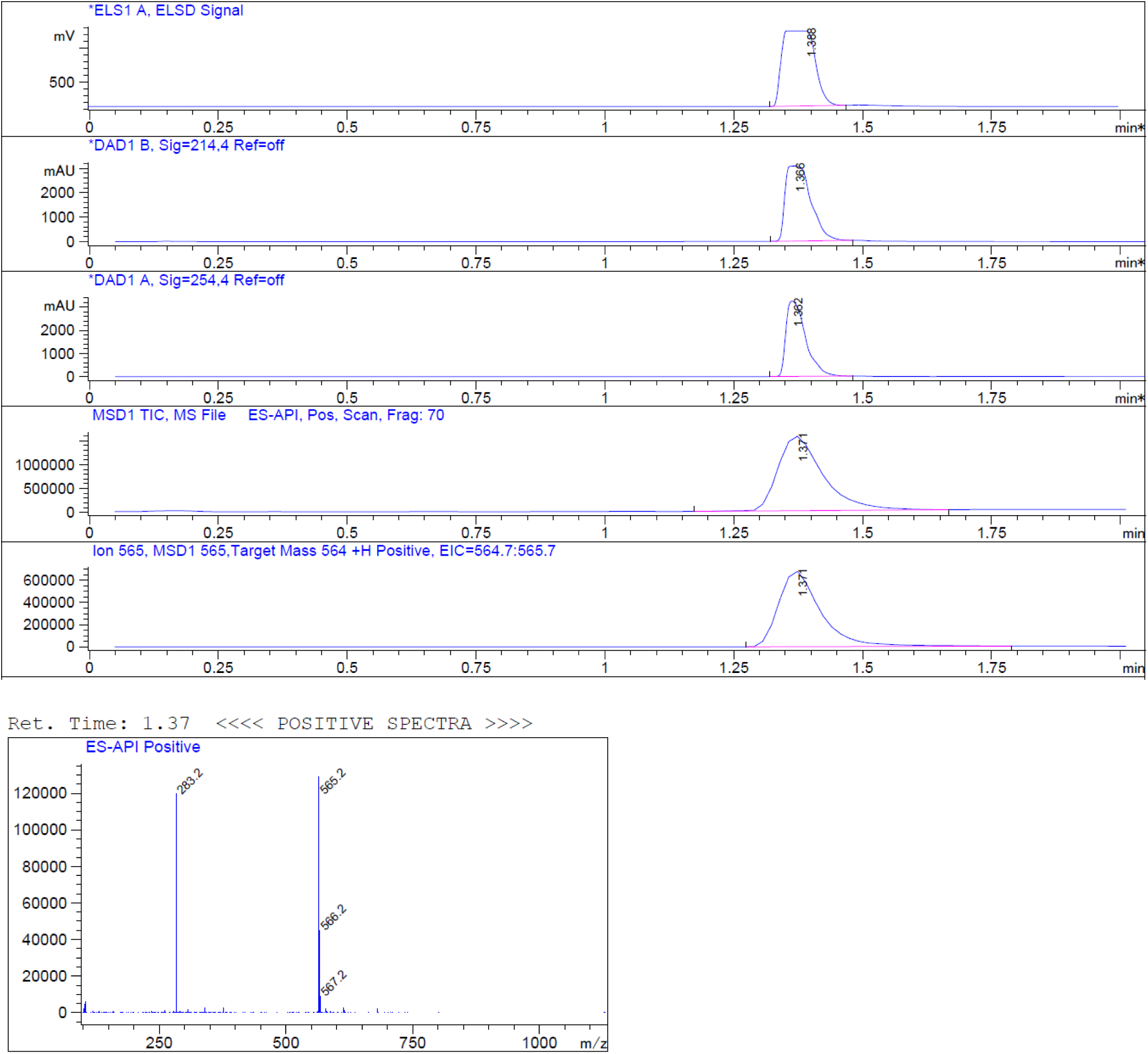

**(7) Synthesis of perfluorophenyl 6-(4-fluoro-5-((6-methoxy-7-(3-(pyrrolidin-1-yl)propoxy)quinazolin-4-yl)oxy)-2-methyl-1H-indol-1-yl)hexanoate (A12)**

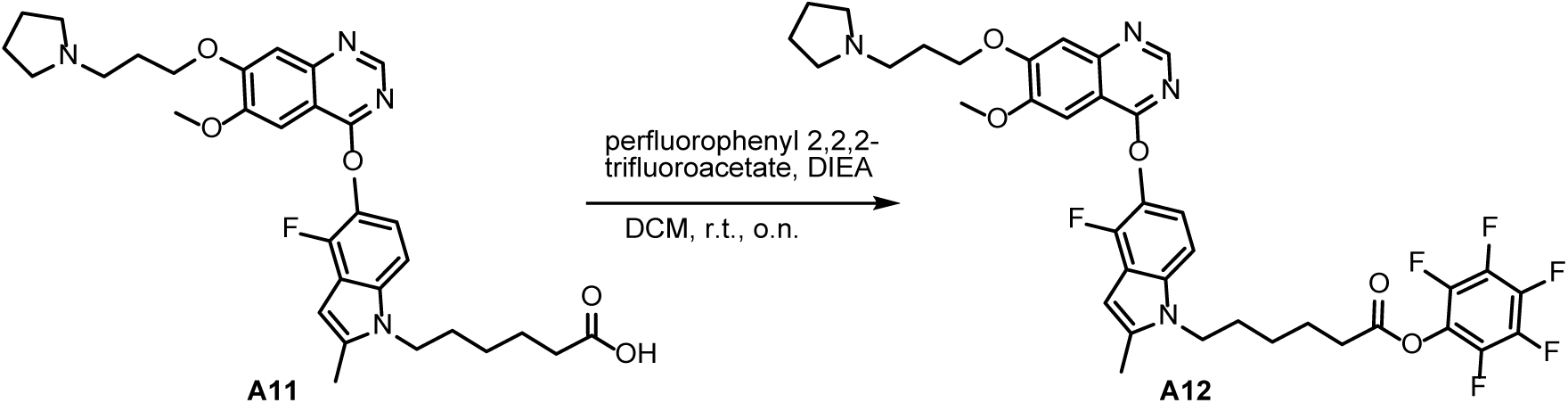

The solution of **A11** (950 mg, 1.68 mmol) in DCM (15 mL) was added DIEA (869 mg, 6.73 mmol). The mixture was stirred under Ar atmosphere at 25 °C for 10 min. Then perfluorophenyl 2,2,2-trifluoroacetate (941 mg, 3.36 mmol) was added and then the mixture was stirred under Ar atmosphere at 25 °C overnight. LCMS showed the reaction was completed. The mixture was extracted with DCM and diluted with water. The organic layers were washed with NaCl saturation and dried over anhydrous Na_2_SO_4_. After filtration, the organic layers were concentrated under reduced pressure. The residue was purified by silica gel column chromatograph, eluted with MeOH: DCM= 15% to afford product **A12** (white solid, 1100 mg, 1.51 mmol, yield: 89.7%).

**LCMS:** *m/z* = 731.2 [M+H]^+^ *t*_R_ = 1.638 min. **Purity:** 77.8% (254 nm).

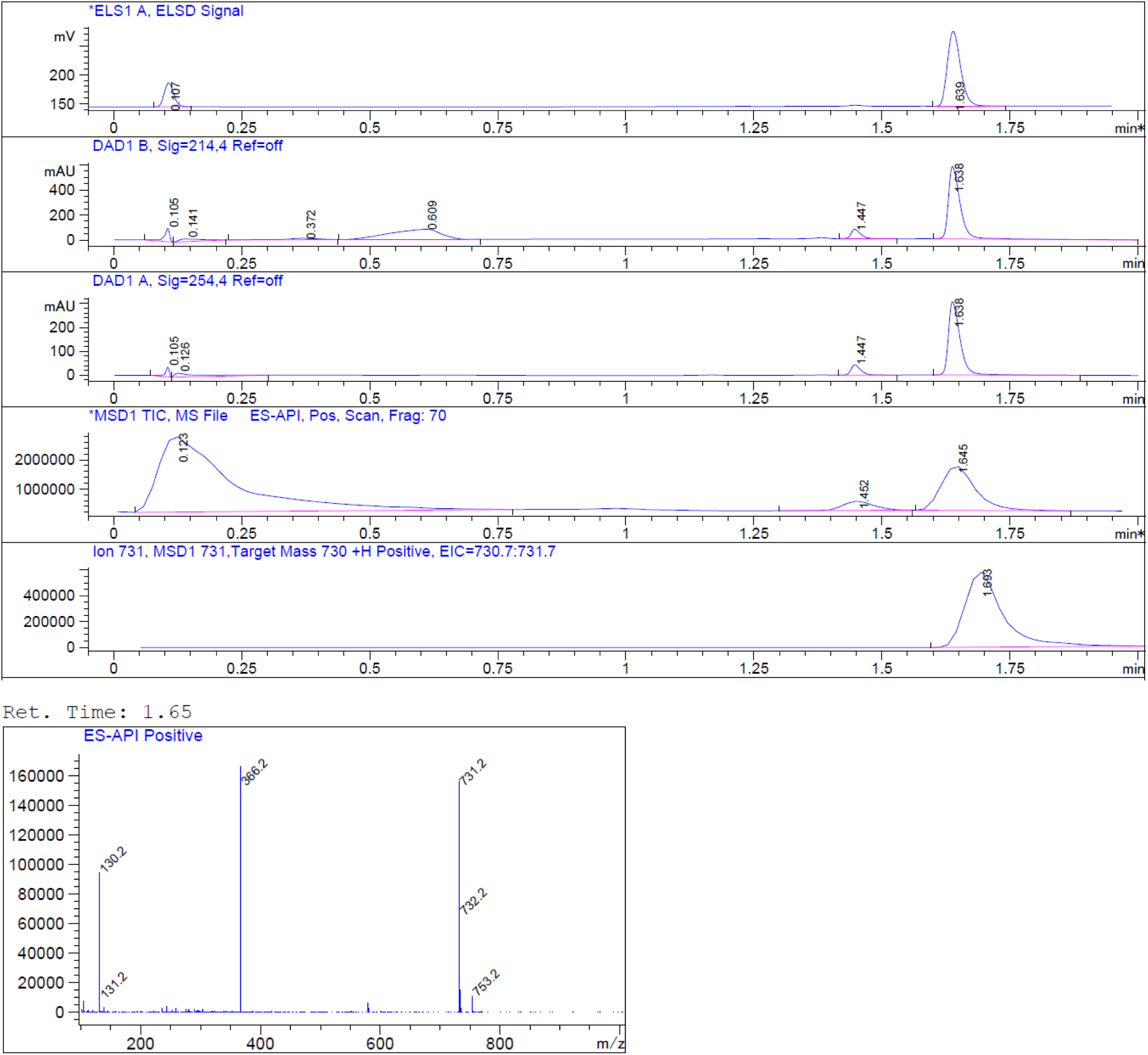

**(8) Synthesis of 1-((2R,4S)-2-((bis(4-methoxyphenyl)(phenyl)methoxy)methyl)-4-hydroxypyrrolidin-1-yl)-6-(4-fluoro-5-((6-methoxy-7-(3-(pyrrolidin-1-yl)propoxy)quinazolin-4-yl)oxy)-2-methyl-1H-indol-1-yl)hexan-1-one (A14)**

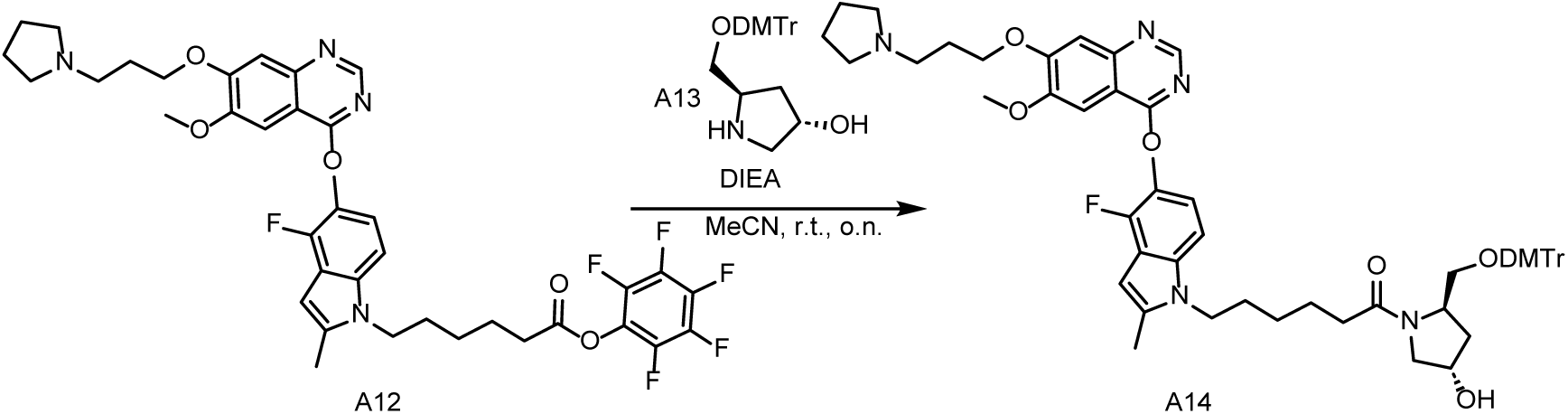

To a solution of **A12** (1100 mg, 1.51 mmol) in MeCN (25 mL) was added DIEA (780 mg, 6.04 mmol). The mixture was stirred under Ar atmosphere at 25 °C for 10 min. Then **A13** (3*S*,5*R*)-5-((bis(4-methoxyphenyl) (phenyl) methoxy) methyl) pyrrolidin-3-ol (949 mg, 2.27 mmol) was added. The mixture was stirred under Ar atmosphere at 25 °C overnight. LCMS showed the reaction was complete. The mixture was diluted with water (100 mL) and extracted with DCM (100 mL*3). The organic layers were wash with brine (100 mL), dried over anhydrous Na_2_SO_4_. After filtration, the organic layer was concentrated under reduced pressure. The residue was purified by silica gel column chromatograph, eluted with MeOH: DCM= 15% to afford product **A14** (white solid, 800 mg, 0.83 mmol, yield: 54.8 %).

**LCMS:** *m/z* = 966.5 [M+H]^+^ *t*_R_ =1.438 min. **Purity:** 95 % (254 nm).

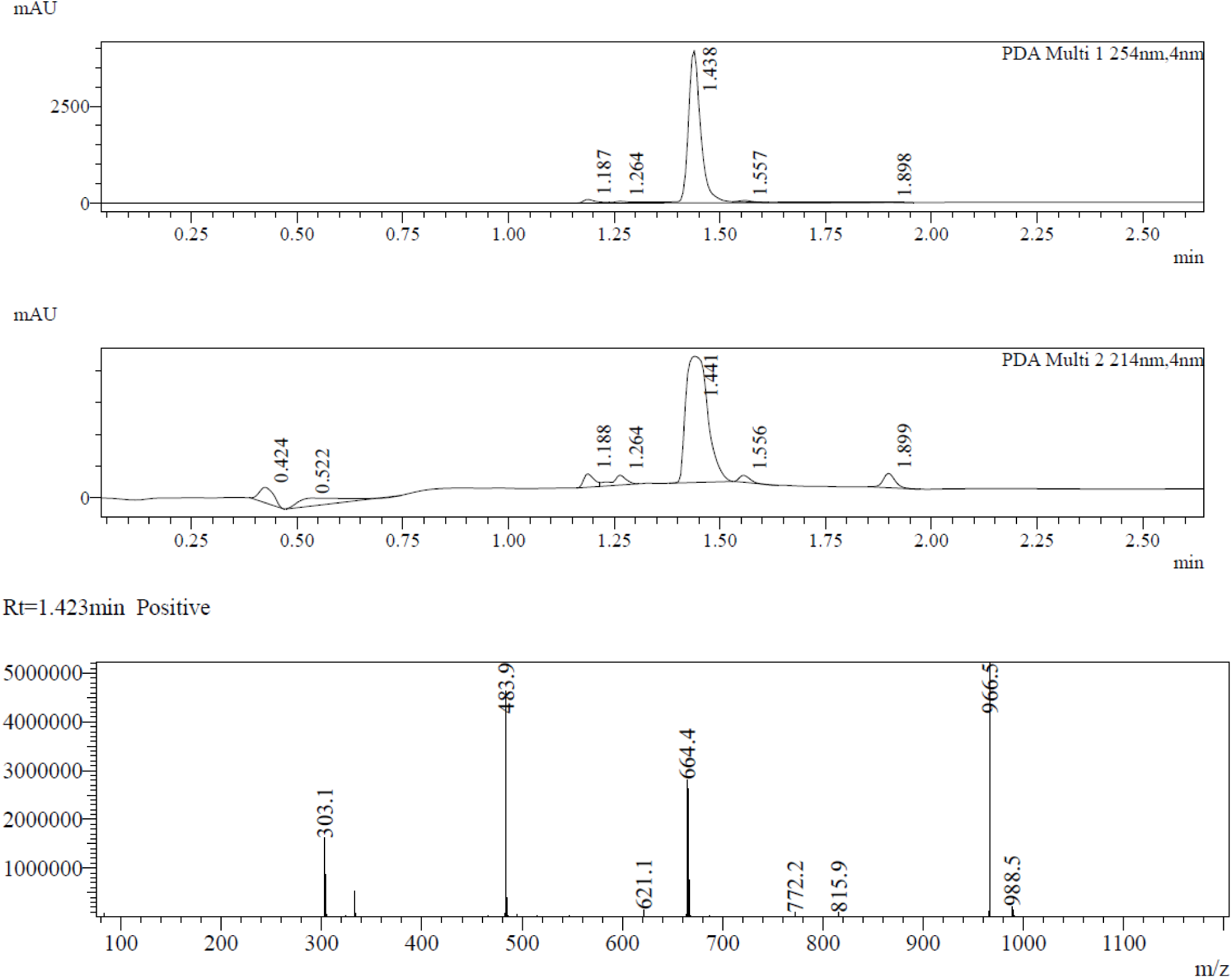

**(9) Synthesis of 4-(((3S,5R)-5-((bis(4-methoxyphenyl)(phenyl)methoxy)methyl)-1-(6-(4-fluoro-5-((6-methoxy-7-(3-(pyrrolidin-1-yl)propoxy)quinazolin-4-yl)oxy)-2-methyl-1H-indol-1- yl)hexanoyl)pyrrolidin-3-yl)oxy)-4-oxobutanoic acid (A15)**

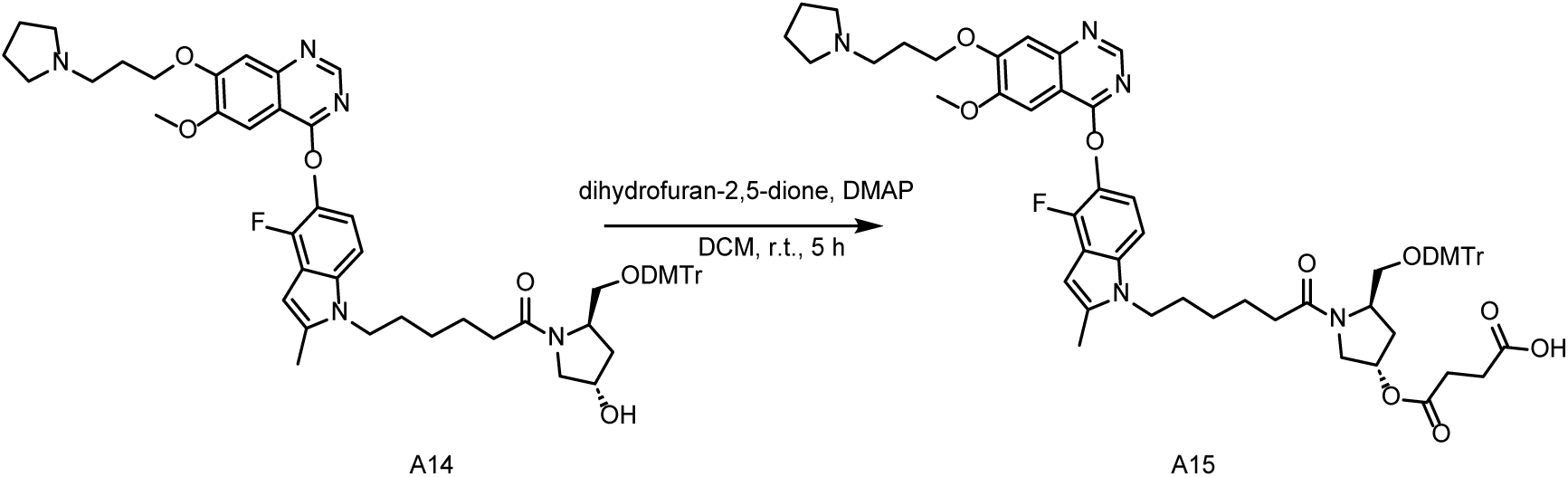

To a solution of **A14** (800 mg, 0.83 mmol) in DCM (15 mL) was added DMAP (405 mg, 3.32 mmol). The mixture was stirred under Ar atmosphere at 25 °C for 10 min. Then dihydrofuran-2,5-dione (124 mg, 1.24 mmol) was added to the system and the mixture was stirred under Ar atmosphere at 25 °C for 5 hours. LCMS showed the reaction was complete. The mixture was diluted with water (100 mL) and extracted with DCM (100 mL*3). The organic layers were wash with brine (100 mL) and dried over anhydrous Na_2_SO_4_. After filtration, the organic layer was concentrated under reduced pressure. The residue was purified by reverse chromatography [ACN/water (TEA: 0.02%): 5%∼30%] to afford the desired product (white solid **A15**, 700 mg, 0.657 mmol, yield: 79%).

**LCMS:** *m/z* = 1064[M-H]^-^ *t*_R_ =2.015 min. **Purity:** 100% (214 nm).

**^1^HNMR** (400 MHz, DMSO-d6) δ 8.48 (s, 1H), 7.60 (s, 1H), 7.39 (s, 1H), 7.36 – 7.14 (m, 10H), 7.09 – 6.98 (m, 1H), 6.88-6.85 (m, 4H), 6.31-6.33 (m, 1H), 5.35-5.25 (m, 1H), 4.26 (t, *J* = 6.4 Hz, 2H), 4.16-4.22 (m, 1H), 4.07-4.14 (m, 2H), 3.99 (s, 3H), 3.79 – 3.66 (m, 7H), 3.55 (d, *J* = 11.6 Hz, 1H), 3.20-3.32 (m, 1H), 3.02-3.07 (m, 2H), 2.59 (t, *J* = 7.2 Hz, 2H), 2.46-2.37 (m, 9H), 2.27-2.18 (m, 3H), 2.03-1.96 (m, 4H), 1.72-1.62 (m, 6H), 1.58-1.48 (m, 2H), 1.43-1.33 (m, 2H).

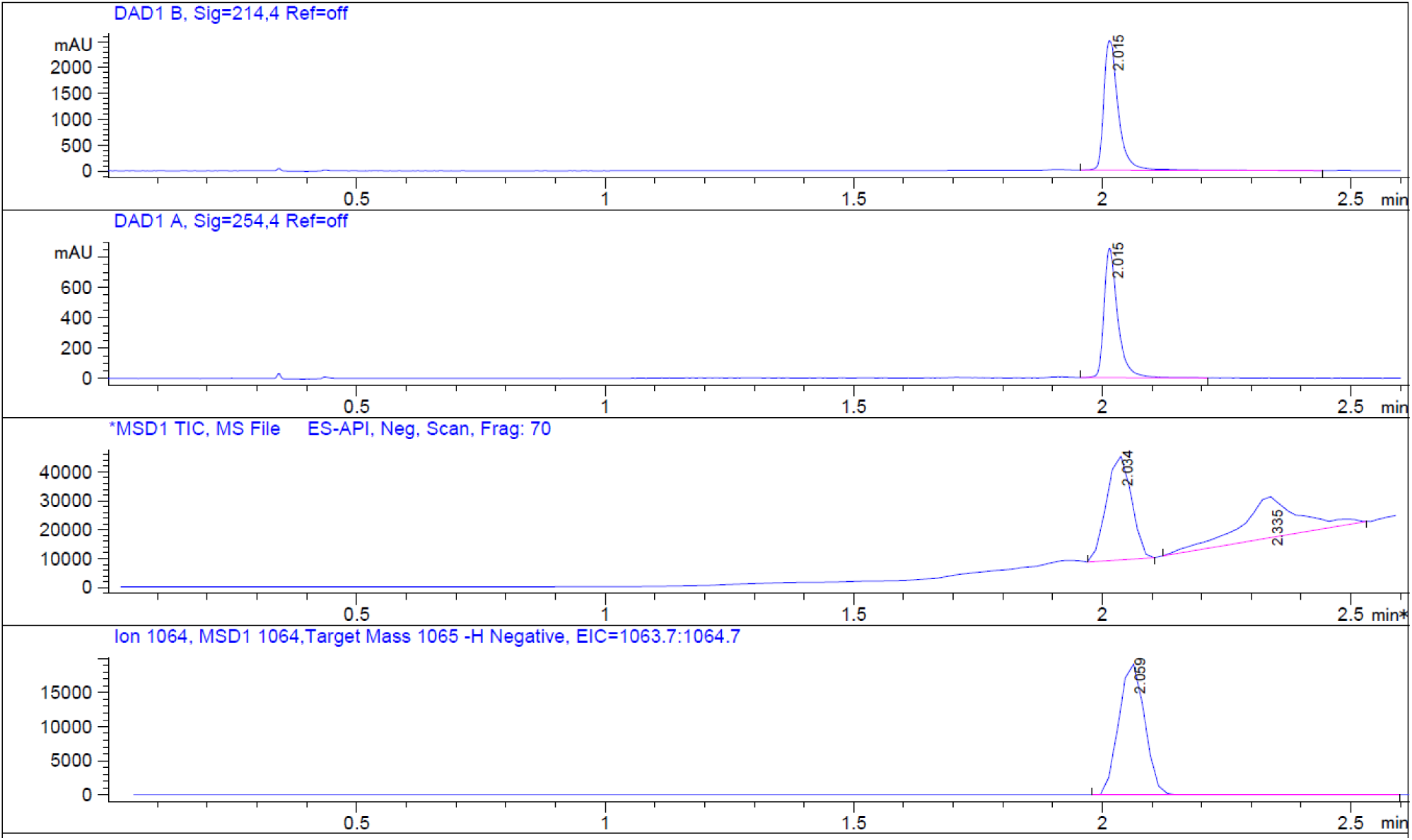
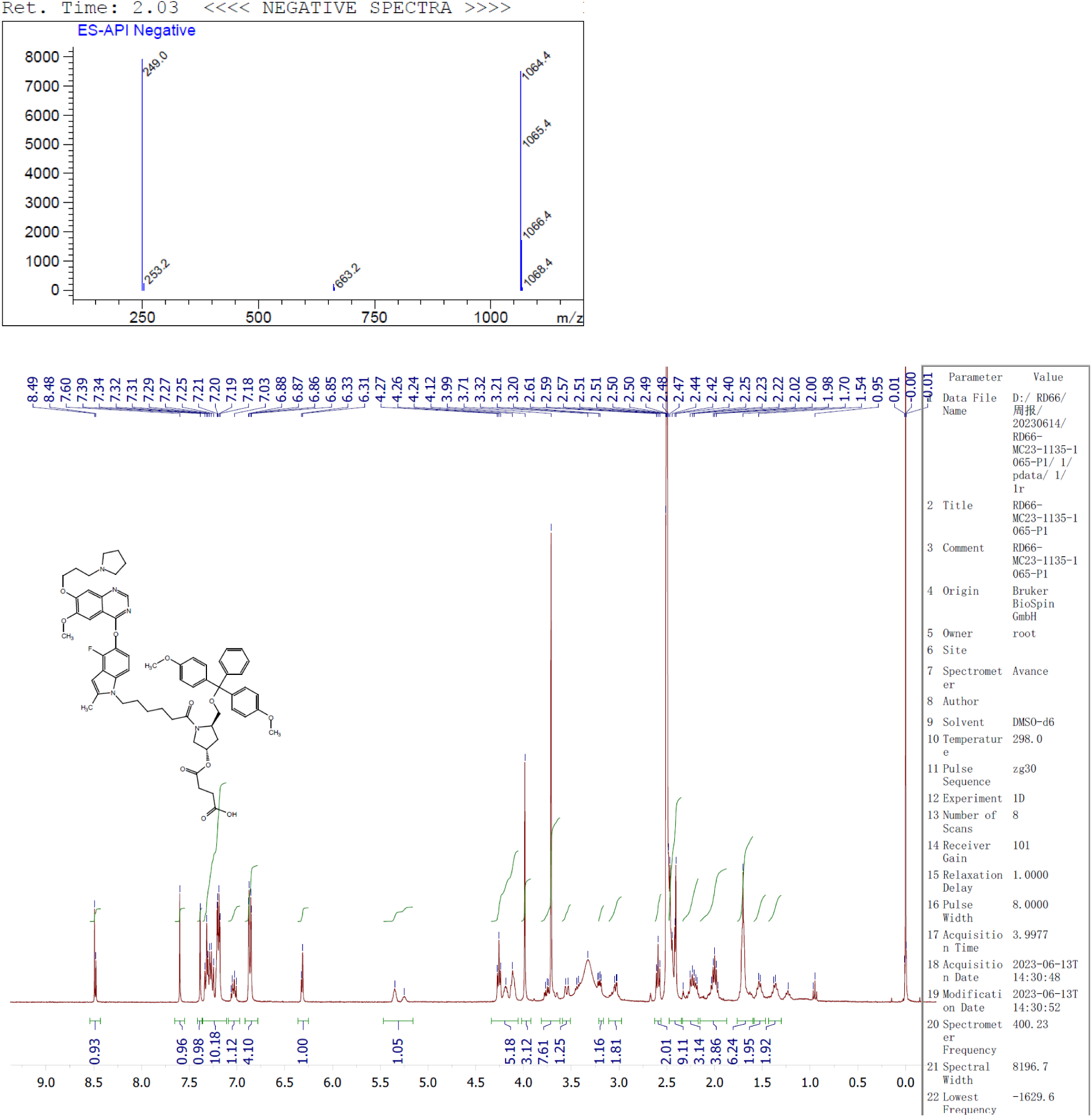

**(10) Synthesis of AS1411-Ced-CPG**

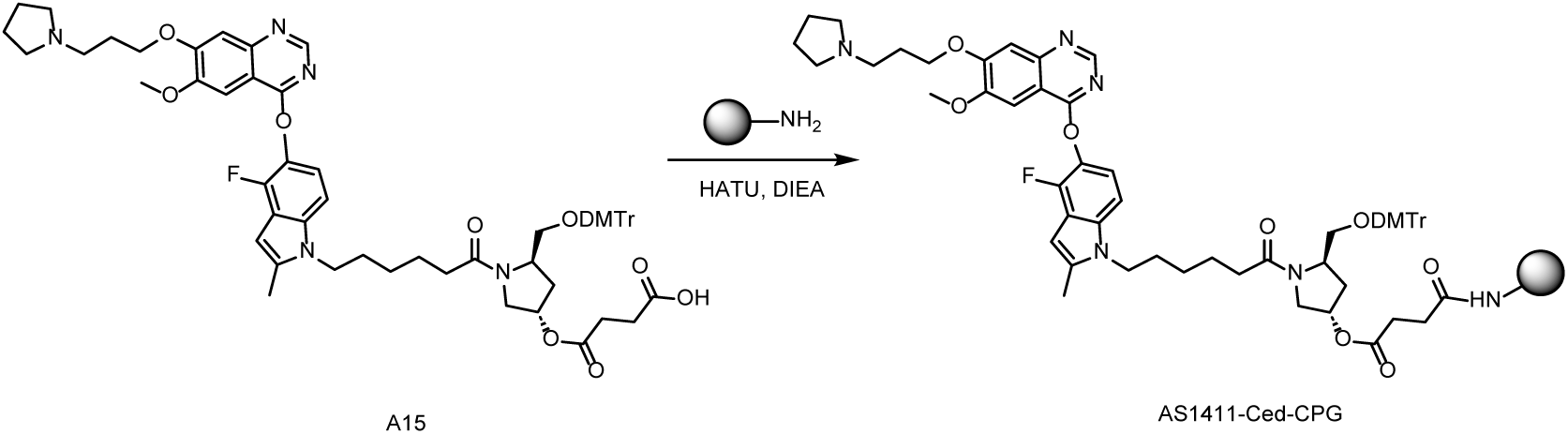

To a solution of **A15** (200 mg, 0.189 mmol) in ACN (12.0 mL) was added HATU (80 mg, 0.208 mmol), DIEA (80 μL), lcaa-CPG (1000 °A, 1000 mg) respectively at room temperature and oscillates for 12 hours. After the reaction was completed, CPG was washed with CAN. CAP A (acetic anhydride: tetrahydrofuran= 1:9, v/v, 4.0 mL) and CAP B (n-methylimidazole: pyridine: acetonitrile= 15:10:75, v/v/v, 4.0 mL) was added, the mixture was oscillated at room temperature for 1 hour. Then the mixture was filtered and washed with ACN (2 mL) for 3 times. **AS1411-Ced-CPG** was obtained after freeze-drying of the mixture as a white power (1000 mg).

**(11) Synthesis of AS1411-Ced**

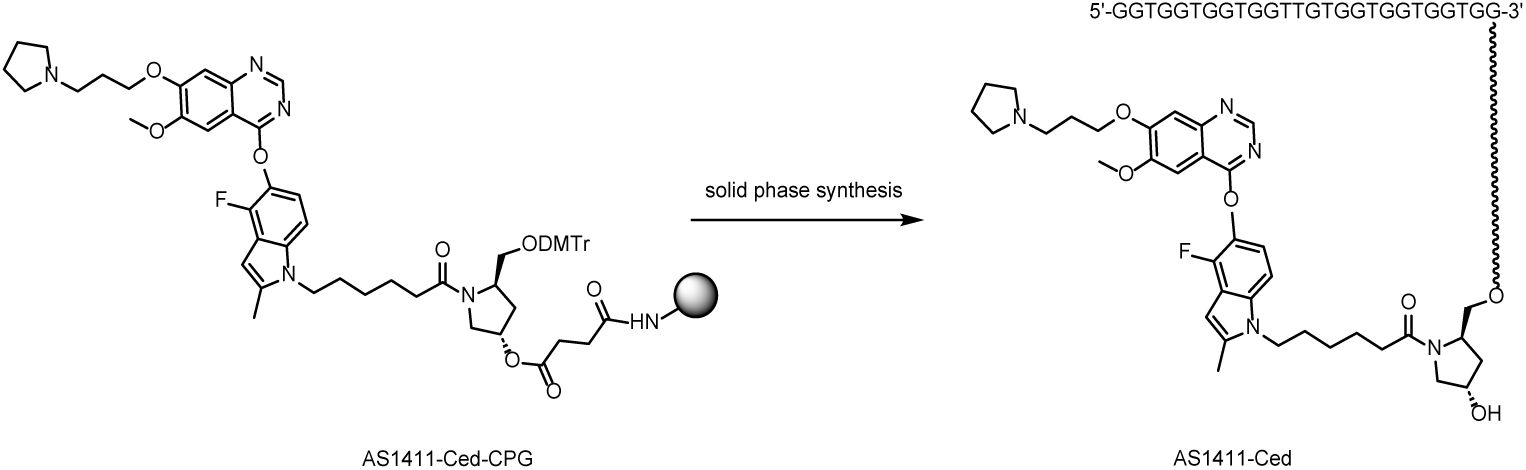

**AS1411-Ced-CPG** in synthesis columns (60 mg*8) and synthesize by K-A H-8 Solid phase synthesizer. Solid phase synthesis involves four steps: Detritylation, Coupling, Capping and Oxidation. After reaction was done, CPG of each synthesis column was added 1.5 mL aqueous ammonia: TEA = 10: 1 and heated in oven at 55 °C for 3 hours. Then collected the supernatant and washed with water (1 mL*3). The crude was purified by RP-IP-HPLC (WATERS 2489, 3767) (**Column:** XBridge Prep C18 10μm OBD^TM^. **RP-IP-HPLC method**:

Mobile phase A: 0.1% TEA+2% HFIP in water, Mobile phase B: methanol) to afford **AS1411-Ced** as a white power (6.66 mg, purity = 96.03%).

**UPLC-MS (WATERS ACQUITY PREMIER):** *m/z* = 8997.77465 [M]^-^(deconvolution); *t*_R_ = 12.044 min (260 nm). Mass error <50 ppm. **HPLC:** *t*_R_ = 11.663 min (260 nm), purity: 96.034%.

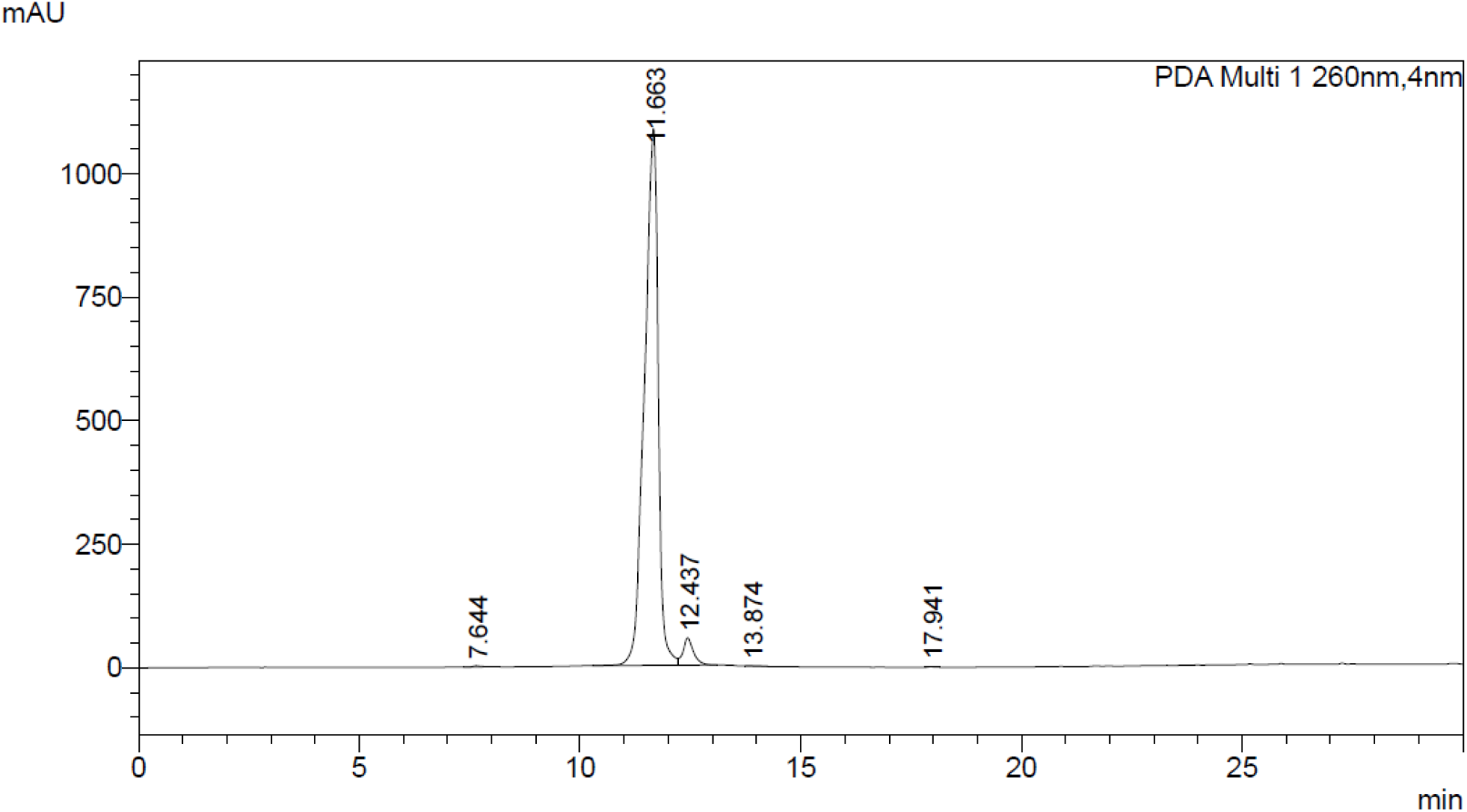
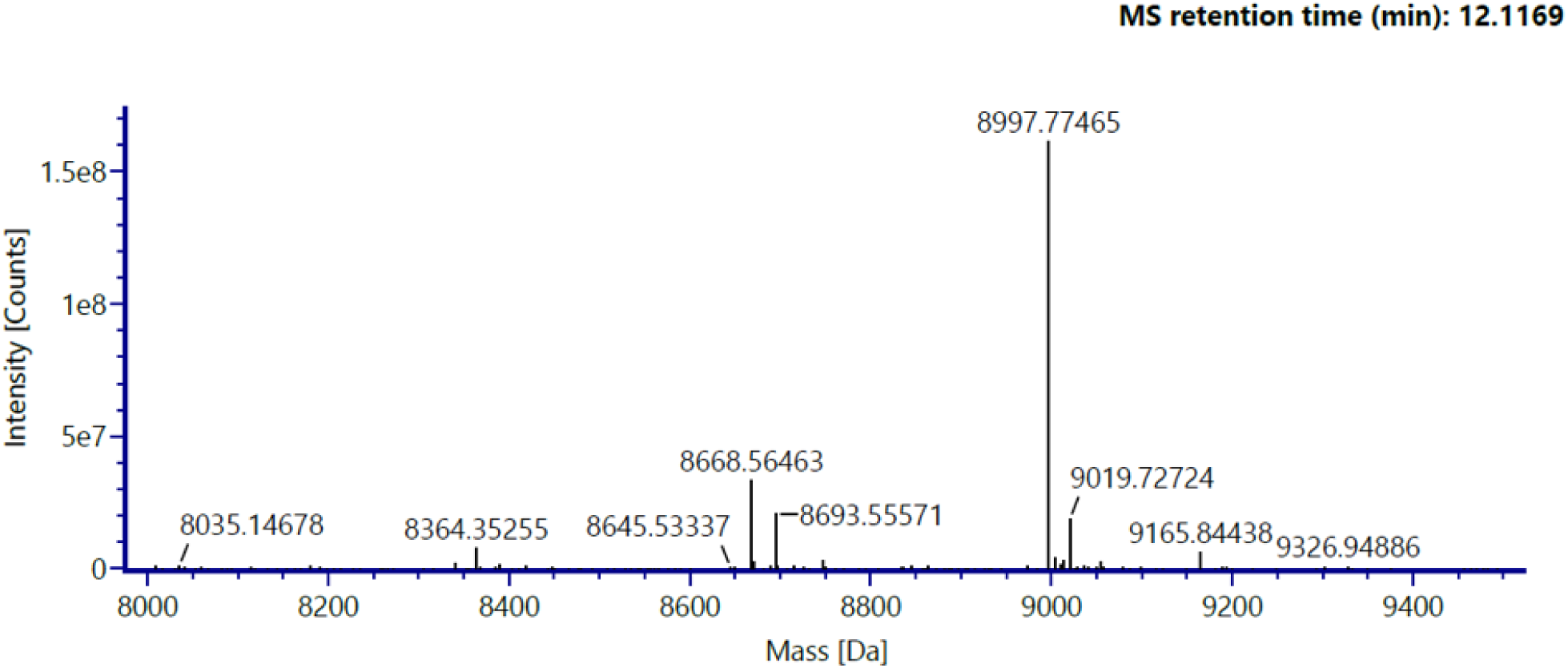

### Scheme 2. Synthetic route for AS1411-Gef

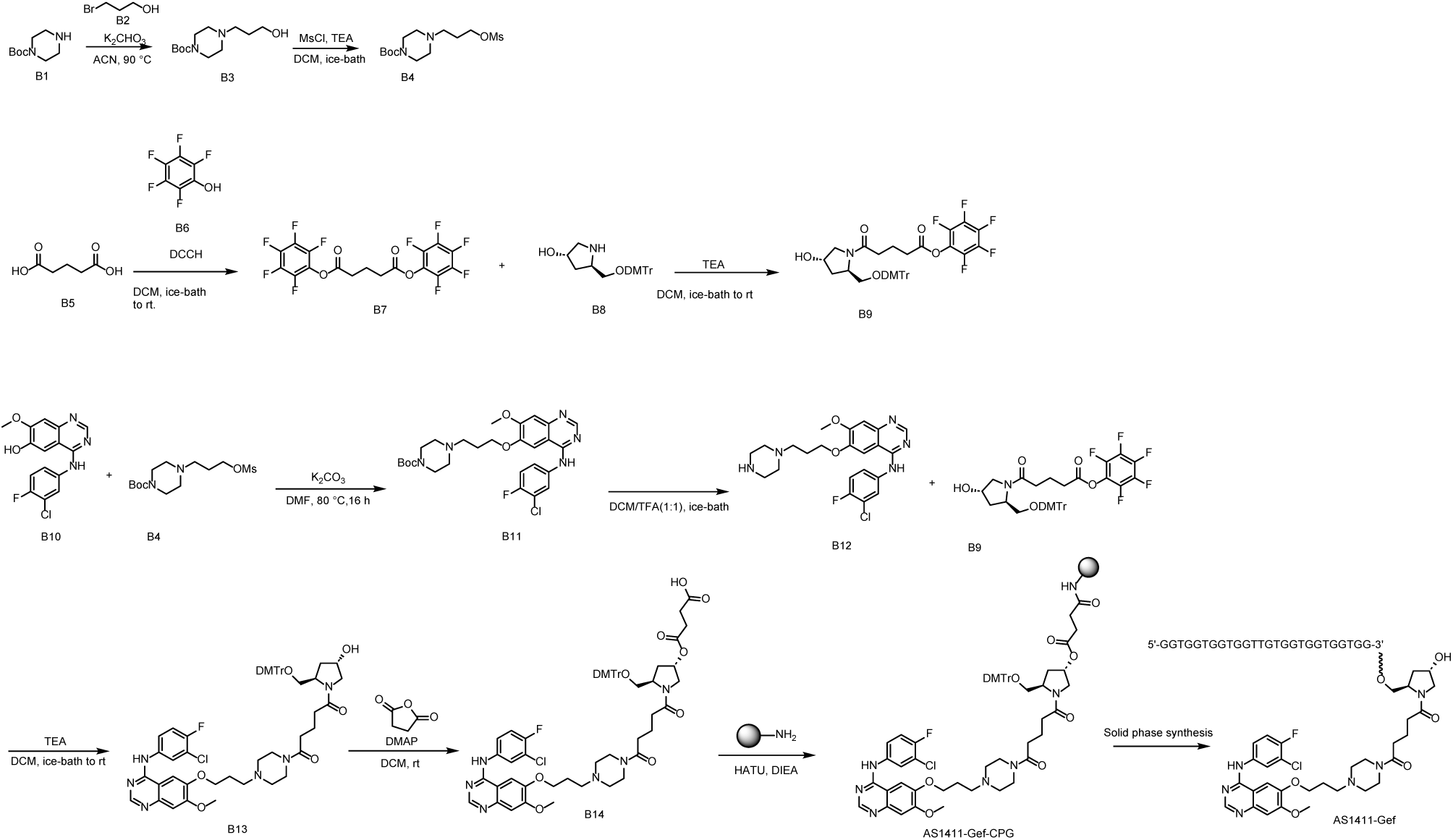

**(1) Synthesis of tert-butyl 4-(3-hydroxypropyl)piperazine-1-carboxylate (B3)**

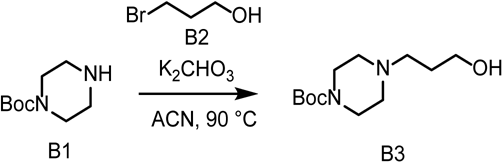

To a solution of **B1** (5.130 g, 27.54 mmol) in ACN (100 mL) was added **B2** (4.27 g, 30.72 mmol), K_2_CO_3_ (7.64 g, 55.28 mmol). The mixture was stirred at 90 degree for 4 hours. The mixture was concentrated and diluted with water (100 mL), extracted with DCM (100 mL*3). The organic phase was dried over Na2SO4, concentrated and purified by DCM: MeOH (0% to 10%) to obtain **B3** (4.7 g, 70 %) as a white solid.

**LCMS:** m/z = 245.3[M+H] + tR =5.347 min. Purity: 100.00% (214 nm). **^1^HNMR** (400 MHz, CDCl3) δ 4.88 (s, 1H), 3.92 – 3.71 (m, 2H), 3.55 – 3.29 (m, 4H), 2.71 – 2.57 (m, 2H), 2.55 – 2.35 (m, 4H), 1.74 (m, 2H), 1.46 (s,9H).

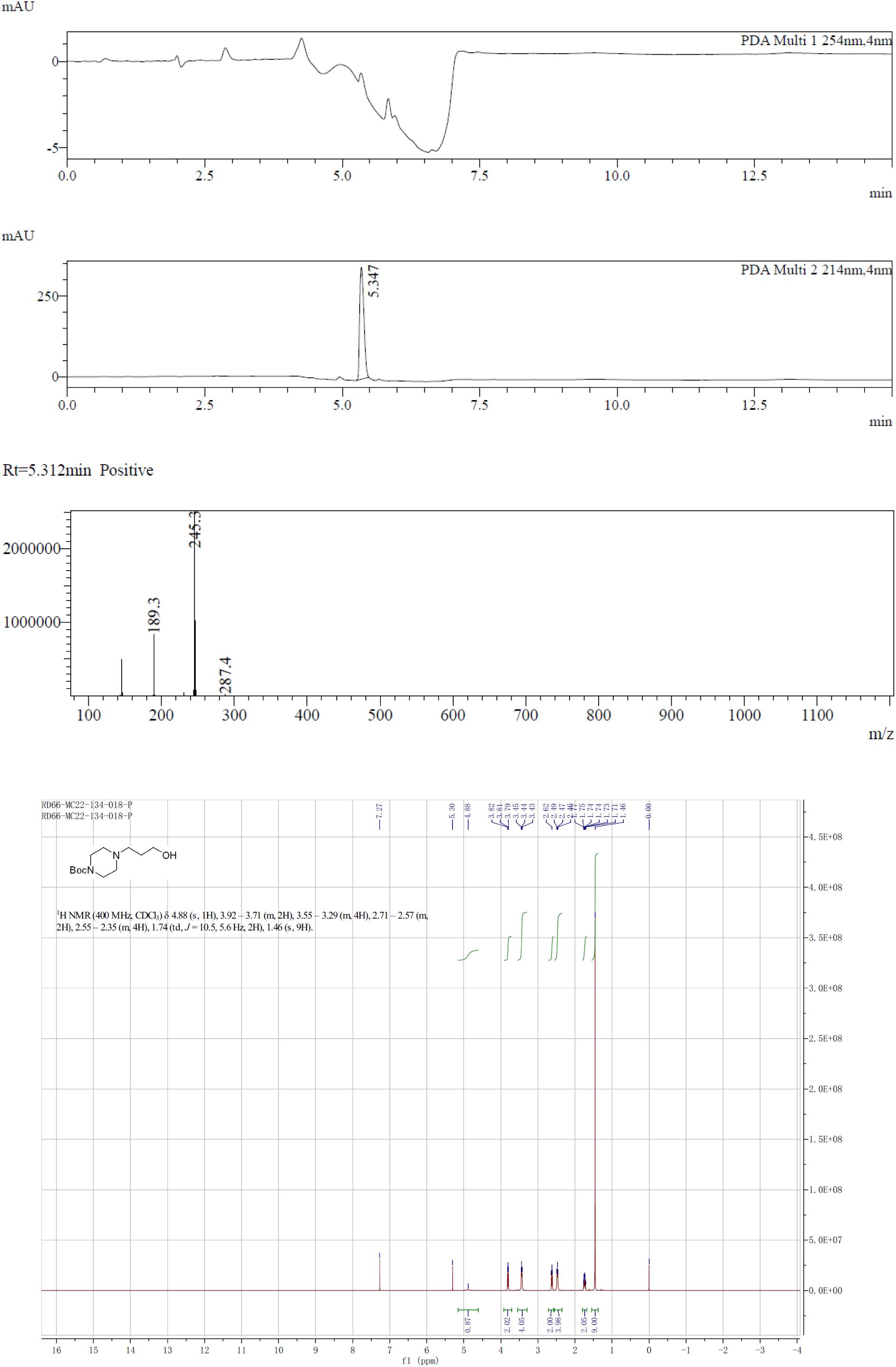

**(2) Synthesis of tert-butyl 4-(3-((methylsulfonyl)oxy)propyl)piperazine-1-carboxylate (B4)**

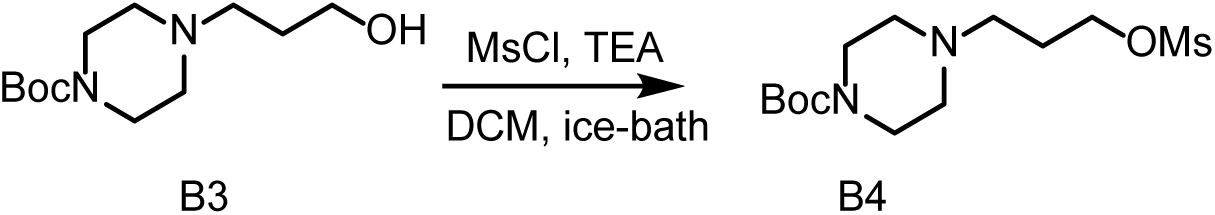

To a solution of **B3** (4.463 g, 18.27 mmol) in DCM (40 mL) was added TEA (3.7 g, 36.56 mmol), MsCl (2.492 g, 21.75 mmol) at 0 degree. The mixture was stirred at this temperature for 1 hour. LCMS showed target MS was found. To the mixture was added water (20 mL) at low temperature, extracted with DCM (3 * 25 mL). The combined organic layers were washed with brine (3 * 30 mL), dried over Na_2_SO_4_, filtered and concentrated in vacuo to give **B4** (5.681 g, 96% yield) as a yellow oil.

**LCMS:** *m/z* = 323.3[M+H]^+^ *t*_R_ = 5.843 min. **Purity:** 100.00% (214 nm). **^1^HNMR** (400 MHz, CDCl_3_) *δ* 4.31 (t, *J* = 6.4 Hz, 2H), 3.48 – 3.34 (m, 4H), 3.01 (s, 3H), 2.47 (t, *J* = 6.8 Hz, 2H), 2.42 – 2.32 (m, 4H), 1.93 (m, 2H), 1.46 (s, 9H).

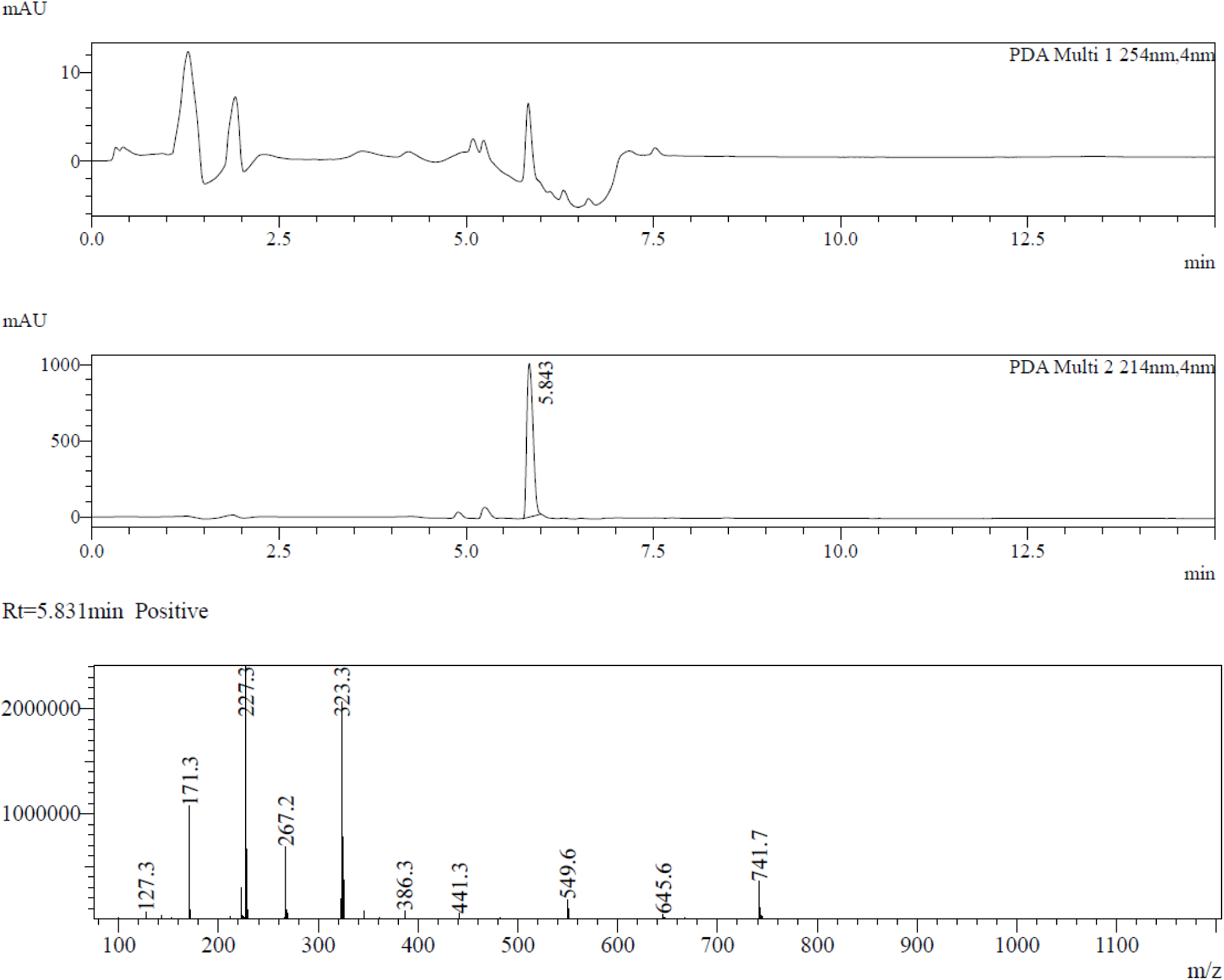
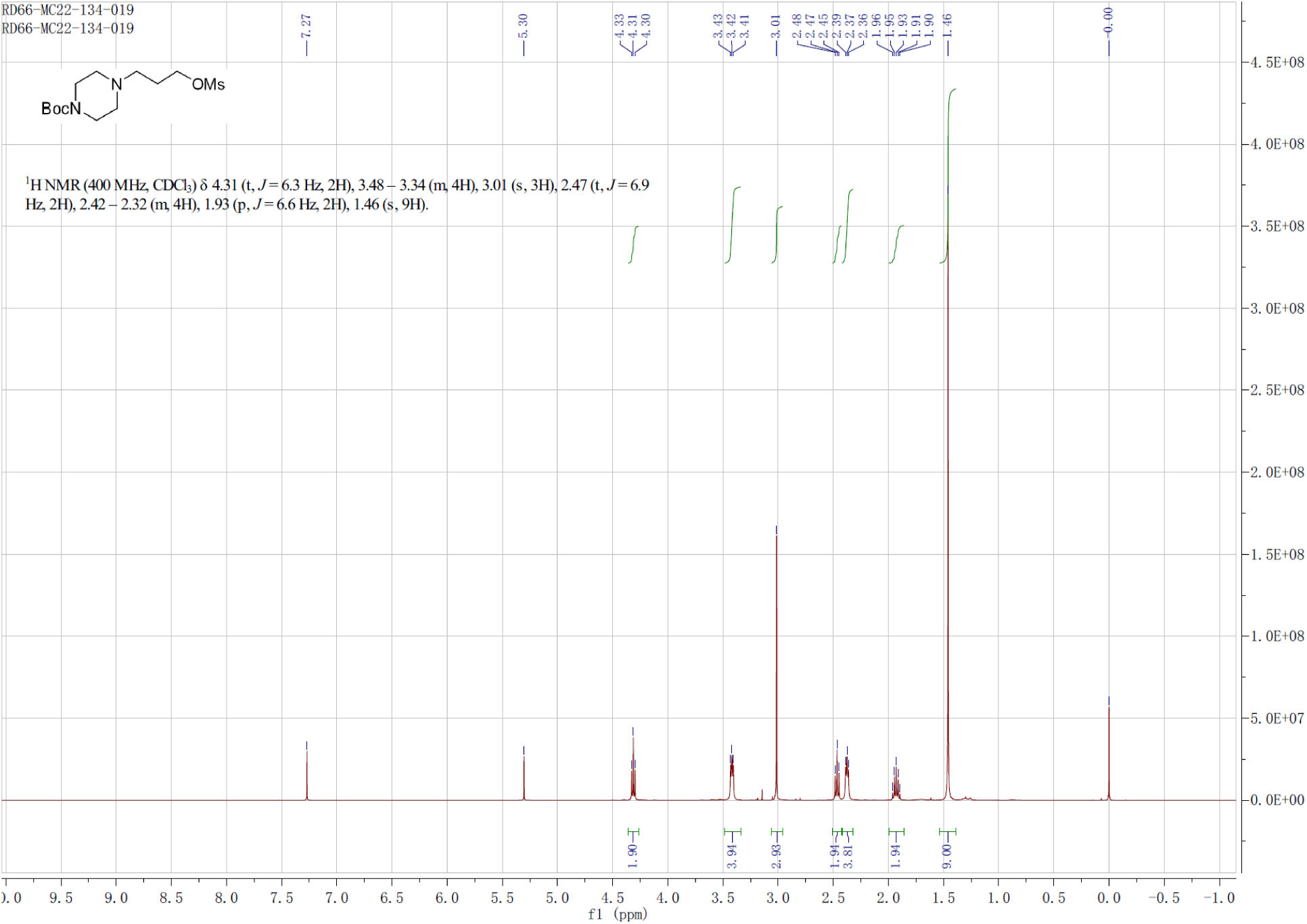

**(3) Synthesis of bis(perfluorophenyl) glutarate (B7)**

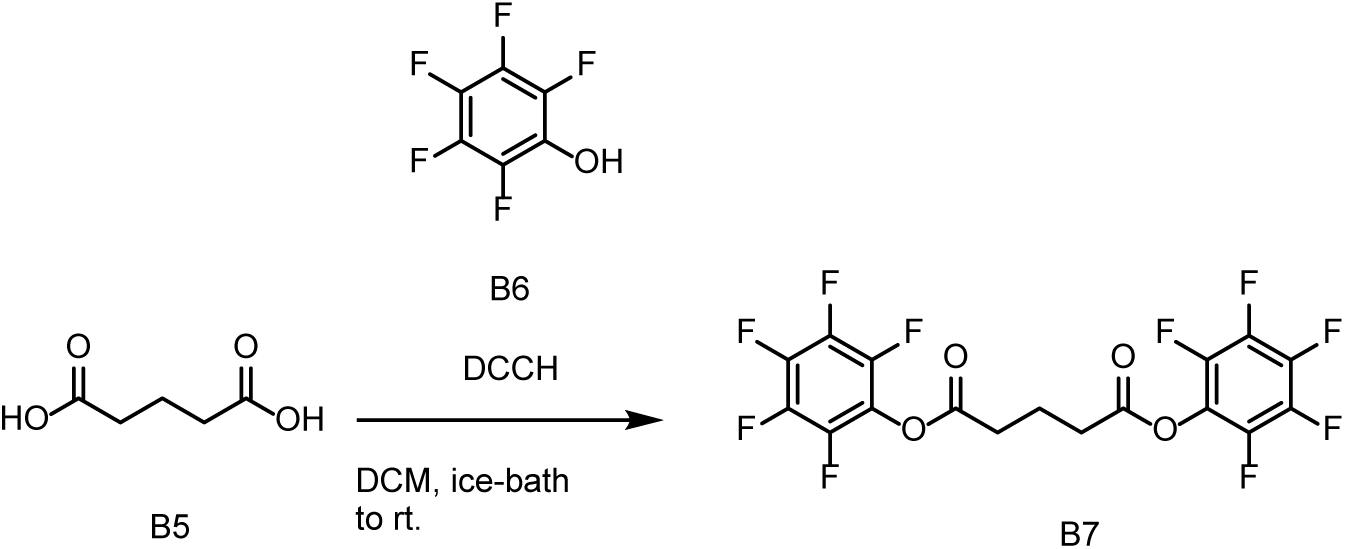

To a solution of **B5** (10 g, 75.69 mmol) and **B6** (33.5 g, 181.99 mmol) in DCM (120 mL) was added DCC (39 g, 189.16 mmol) slowly at 0 degree. The mixture was stirred at room temperature overnight. The reaction mixture was filtered and purified by SGC (DCM:100%) to obtain **B7** (25 g, 71 %) as a white solid.

**LCMS:** *m/z* = N/A[M+H]^+^ *t*_R_ =7.159 min. **Purity:** 100.000% (214 nm). **^1^HNMR** (400 MHz, DMSO-*d*_6_) *δ* 2.95 (t, *J* = 7.2 Hz, 4H), 2.07 (p, *J* = 7.4 Hz, 2H). **^19^FNMR** (400 MHz, DMSO-*d*_6_) *δ* 153.64 (m, 4F), 158.32 (m, 2F), 162.90 (m, 4F).

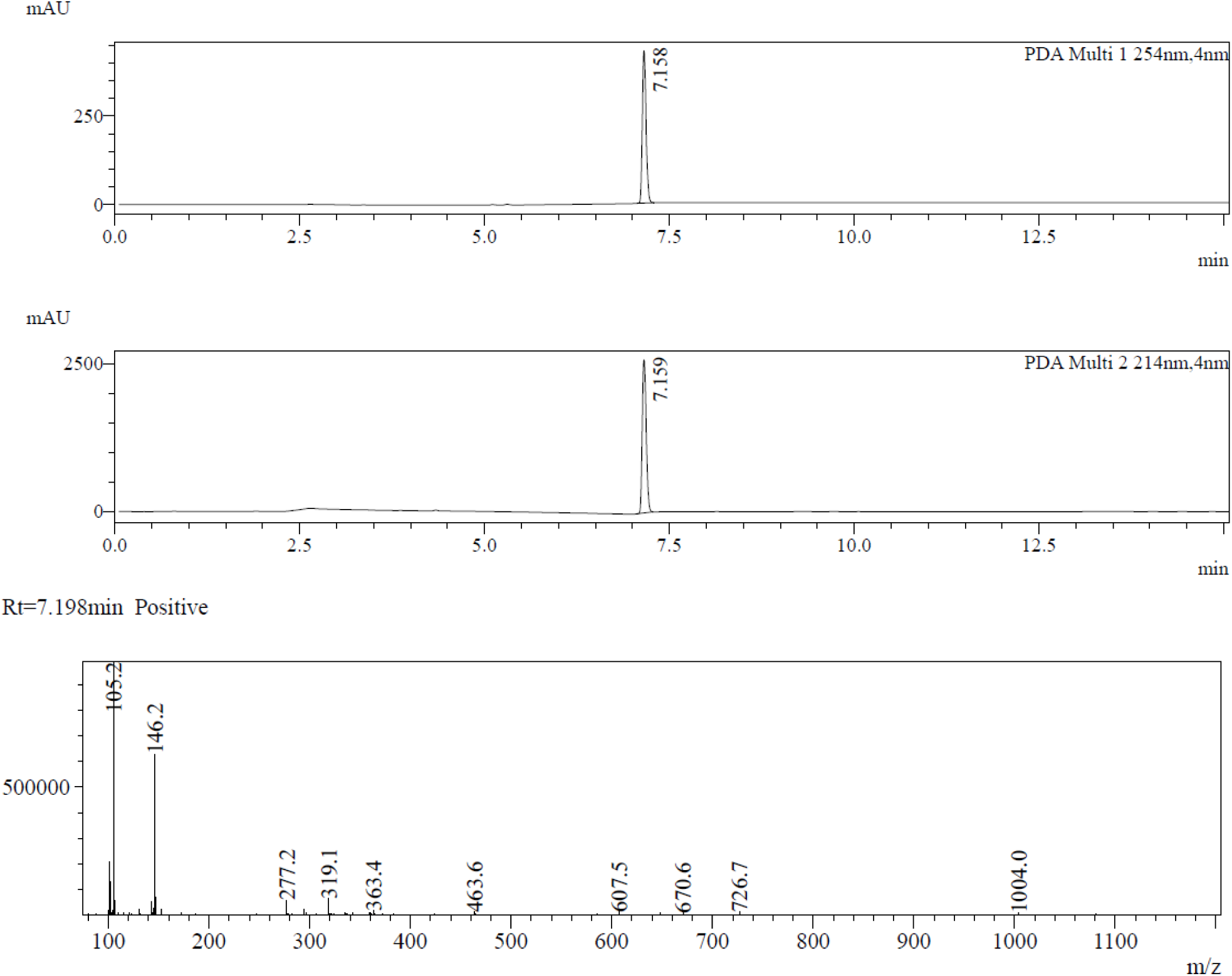
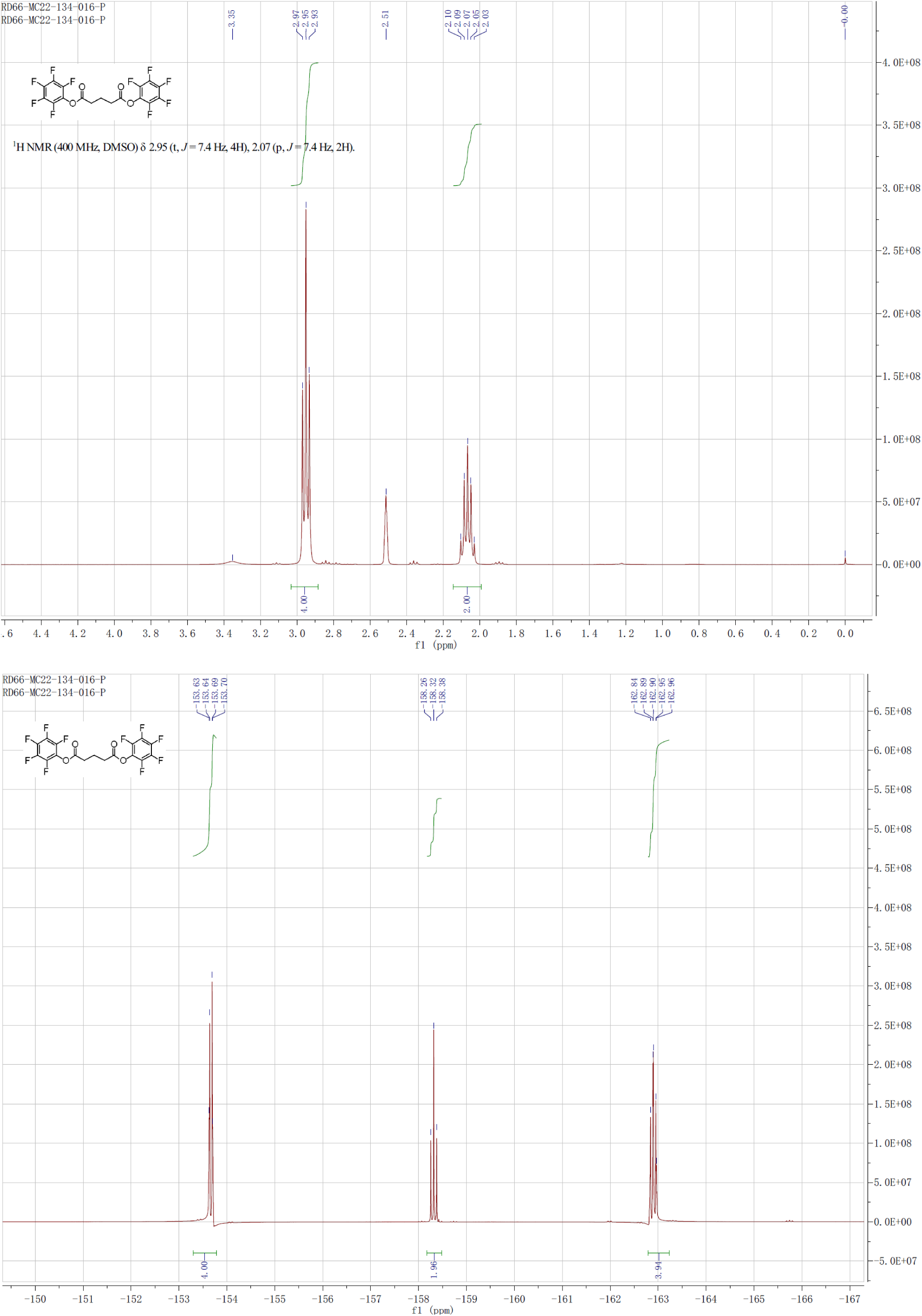

**(4) Synthesis of of perfluorophenyl 5-((2R,4S)-2-((bis(4-methoxyphenyl)(phenyl)methoxy)methyl)-4-hydroxypyrrolidin-1-yl)-5-oxopentanoate (B9)**

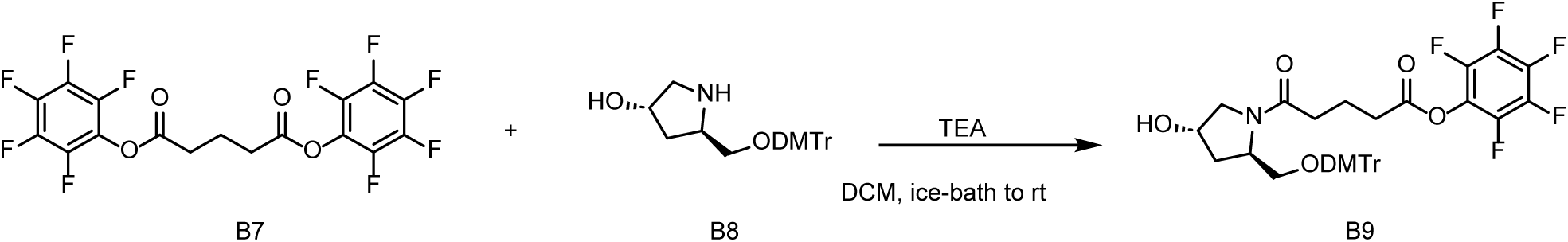

To a solution of **B7** (2.4 g, 5.17 mmol) in DCM (10 mL) was added TEA (1 mL, 7.19 mmol) and **B8** (1.081 g, 2.58 mmol in DCM (10 mL) slowly at 0 degree. The mixture was stirred at room temperature for 10 hours. LCMS showed target MS was found. The mixture was concentrated and purified by SGC (MeOH:DCM, 0% to 10%) to obtain **B19** (1.4 g, crude) as a colorless oil. **LCMS:** m/z = 700.5 [M+H]+ tR = 1.145 min. **Purity:** 100.00% (214 nm).

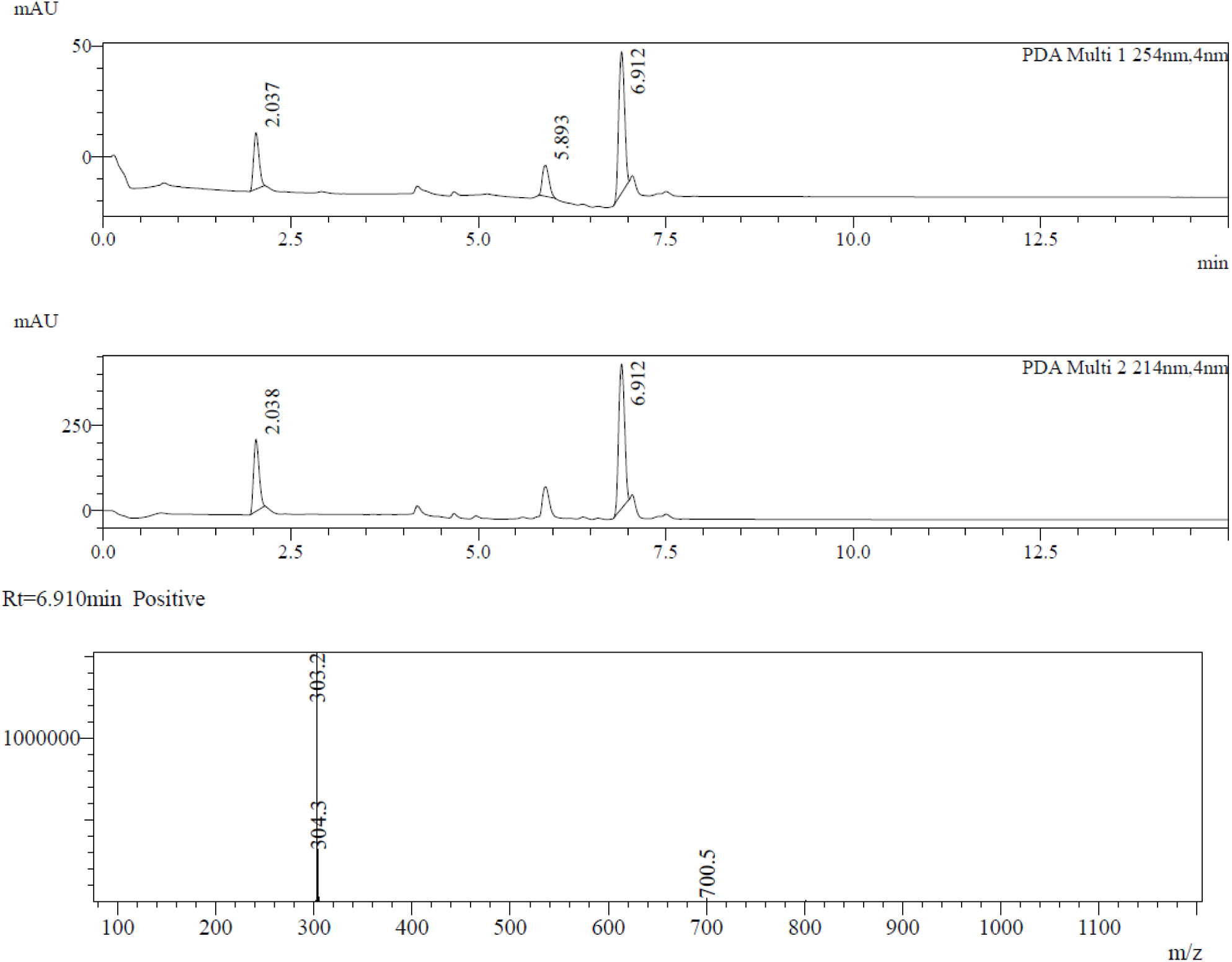

**(5) Synthesis of tert-butyl 4-(3-((4-((3-chloro-4-fluorophenyl)amino)-7-methoxyquinazolin-6-yl)oxy)propyl)piperazine-1-carboxylate (B11)**

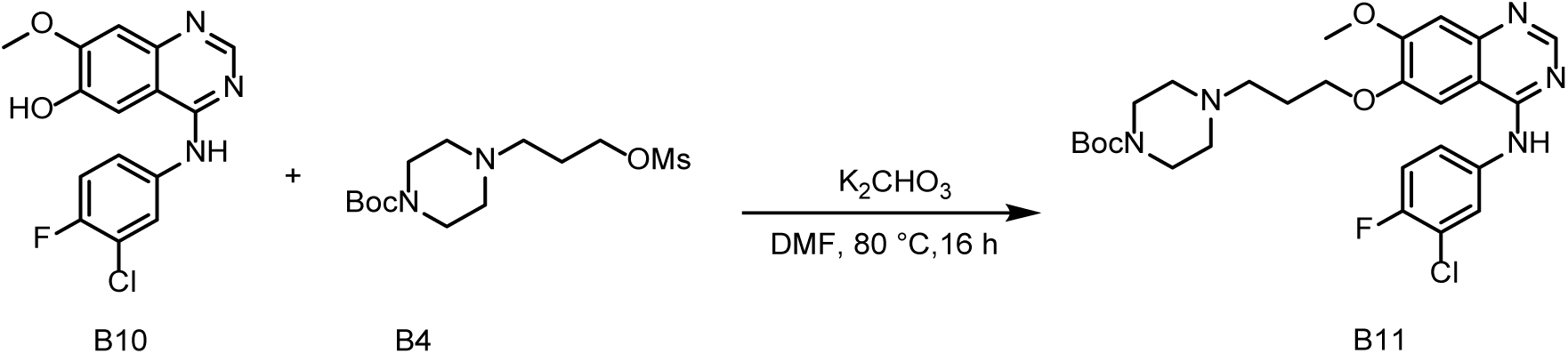

To a solution of **B10** (2.5 g, 7.82 mmol) in DMF (40 mL) was added K_2_CO_3_ (2.5 g, 18.09 mmol), **B4** (4.338 g, 13.61 mmol) under nitrogen and stirred at 80 °C for 16 hours. LCMS showed the target MS was found. The reaction mixture was concentrated, diluted with water (200 mL), extracted with ethyl acetate (100 mL*3). The organic phase was concentrated and purified by chromatography on a silica gel column (EA/PE=20∼100%) to afford **B11** (3.29 g, 77%) as a white solid.

**LCMS:** *m/z* = 546.5 [M+H]^+^ *t*_R_ = 3.853 min. **Purity:** 100.00% (214 nm). **^1^HNMR** (400 MHz, CDCl_3_) *δ* 8.66 (s, 1H), 7.90 – 7.88 (m, 1H), 7.61 – 7.53 (m, 1H), 7.50 (s, 1H), 7.25 (s, 1H), 7.20 – 7.11 (m, 2H), 4.17 (t, *J* = 6.4 Hz, 2H), 3.99 (s, 3H), 3.53 – 3.39 (m, 4H), 2.60 (t, *J* = 7.2 Hz, 2H), 2.50 – 2.42 (m, 4H), 2.14 – 2.09 (m, 2H), 1.46 (s, 9H). ^19^FNMR (400 MHz, CDCl3) *δ* 120.94 (s, 1F)

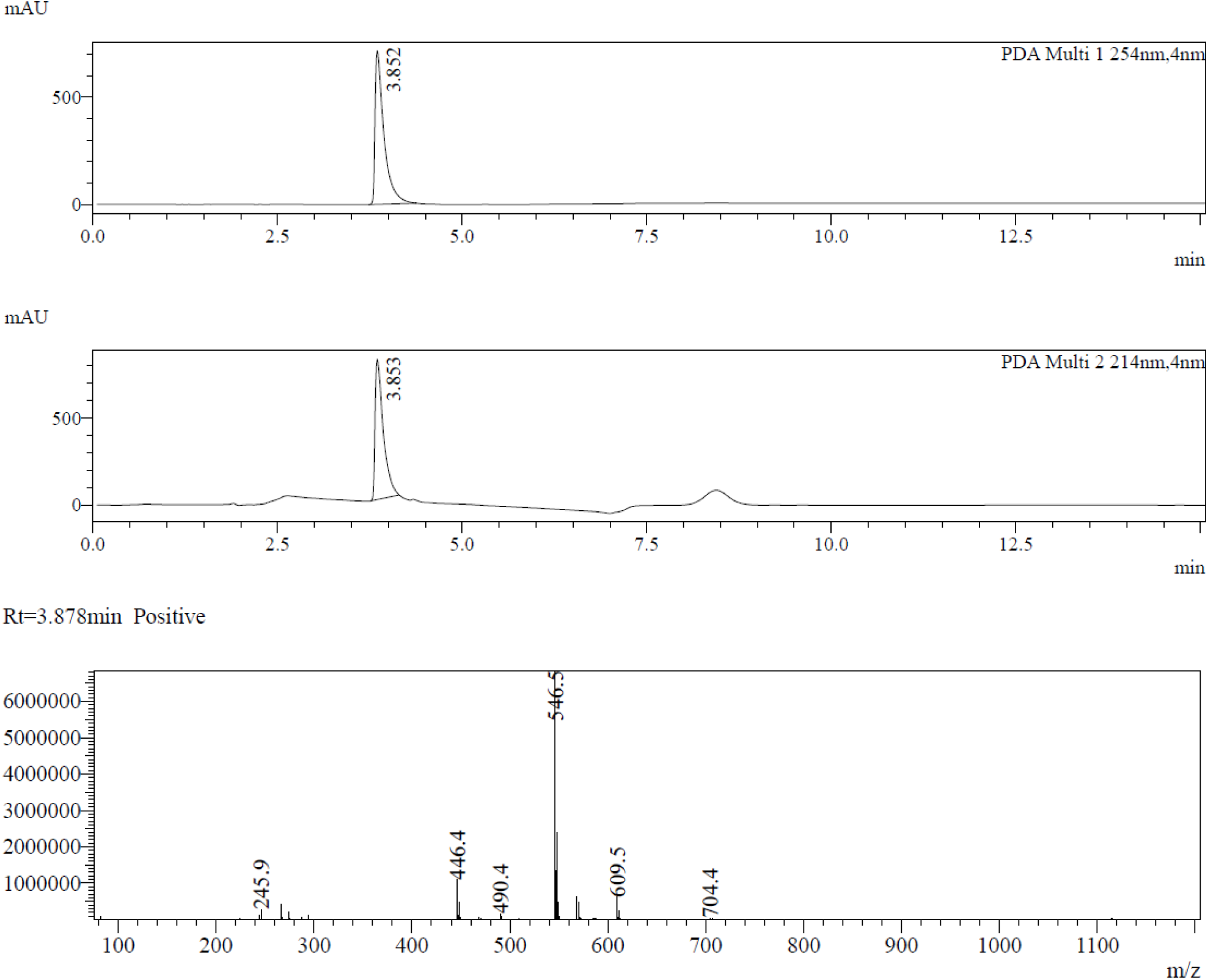
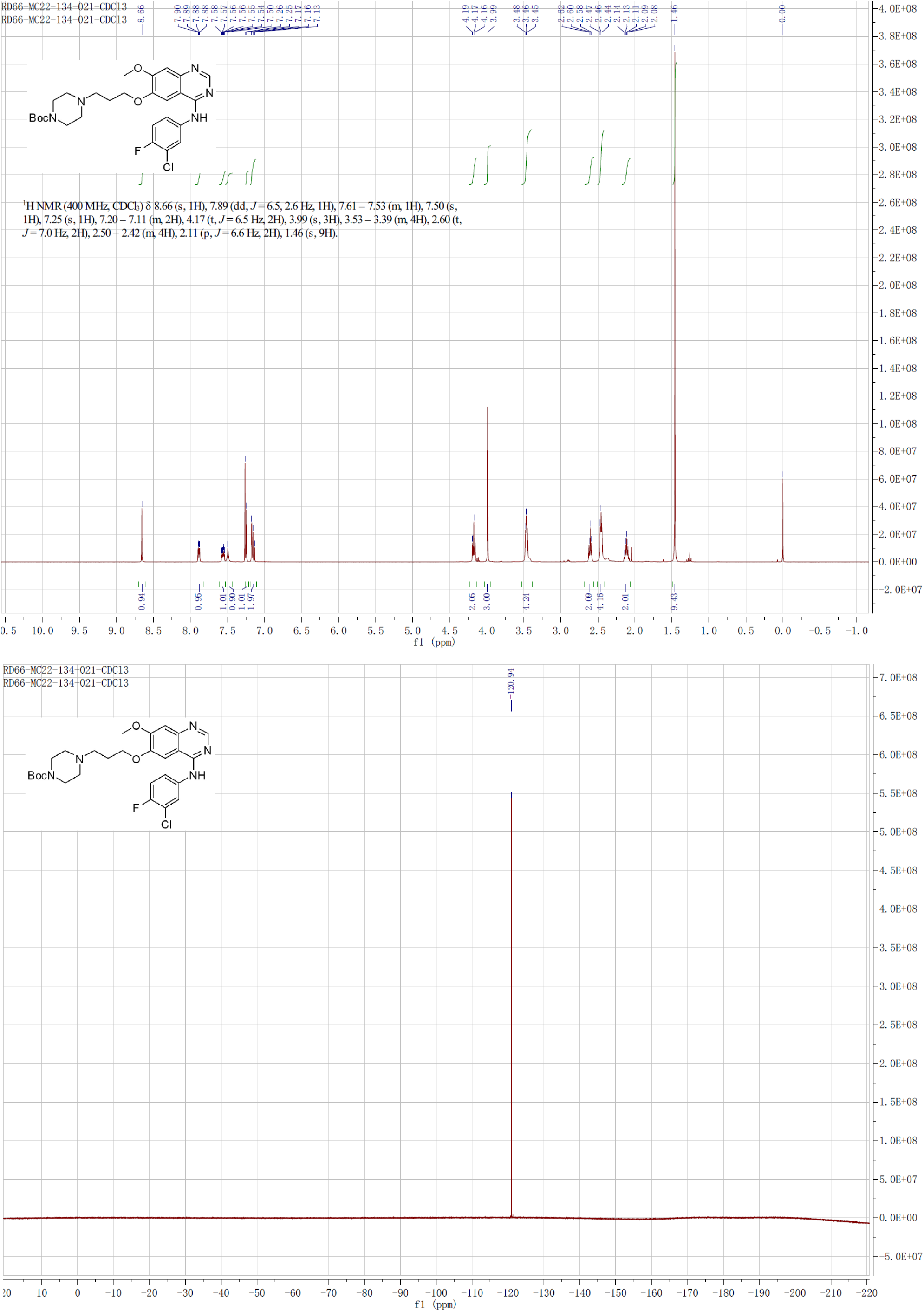

**(6) Synthesis of N-(3-chloro-4-fluorophenyl)-7-methoxy-6-(3-(piperazin-1-yl)propoxy)quinazolin-4-amine (B12)**

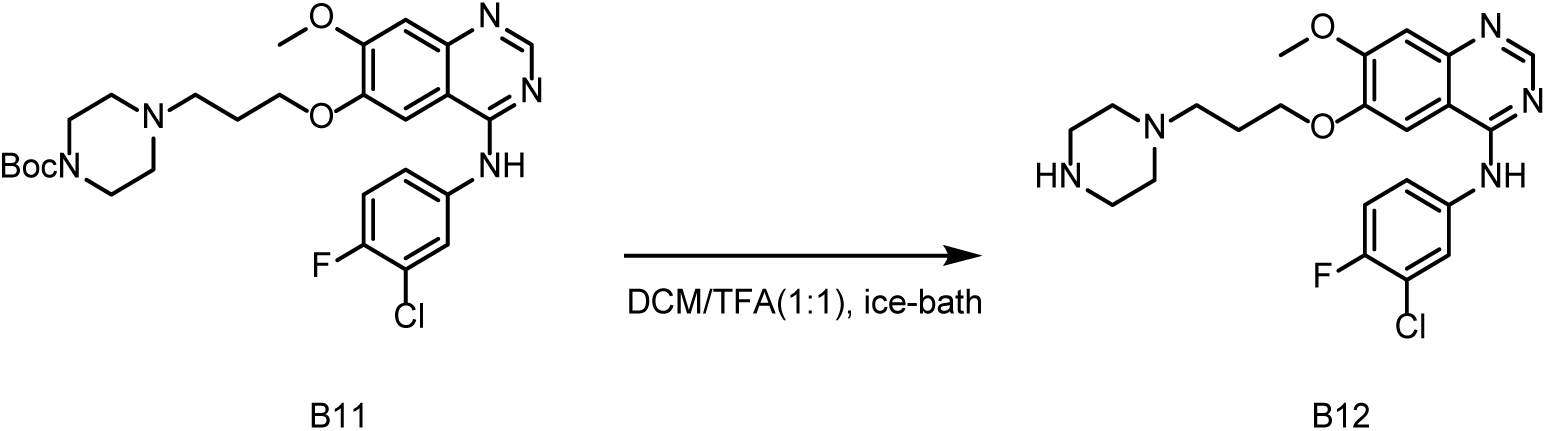

To a solution of **B11** (1.008 g) in DCM (10 mL) was added TFA (10 mL) under nitrogen and stirred at 0 degree for 1 hour. LCMS showed target MS was found. The reaction mixture was concentrated under vacuo to afford **B12** (about 1.4 g, TFA salt) as a brown oil which was used directly.

**LCMS:** *m/z* = 446.3[M+H]^+^ *t*_R_ = 0.758 min. **Purity:** 100.00% (214 nm).

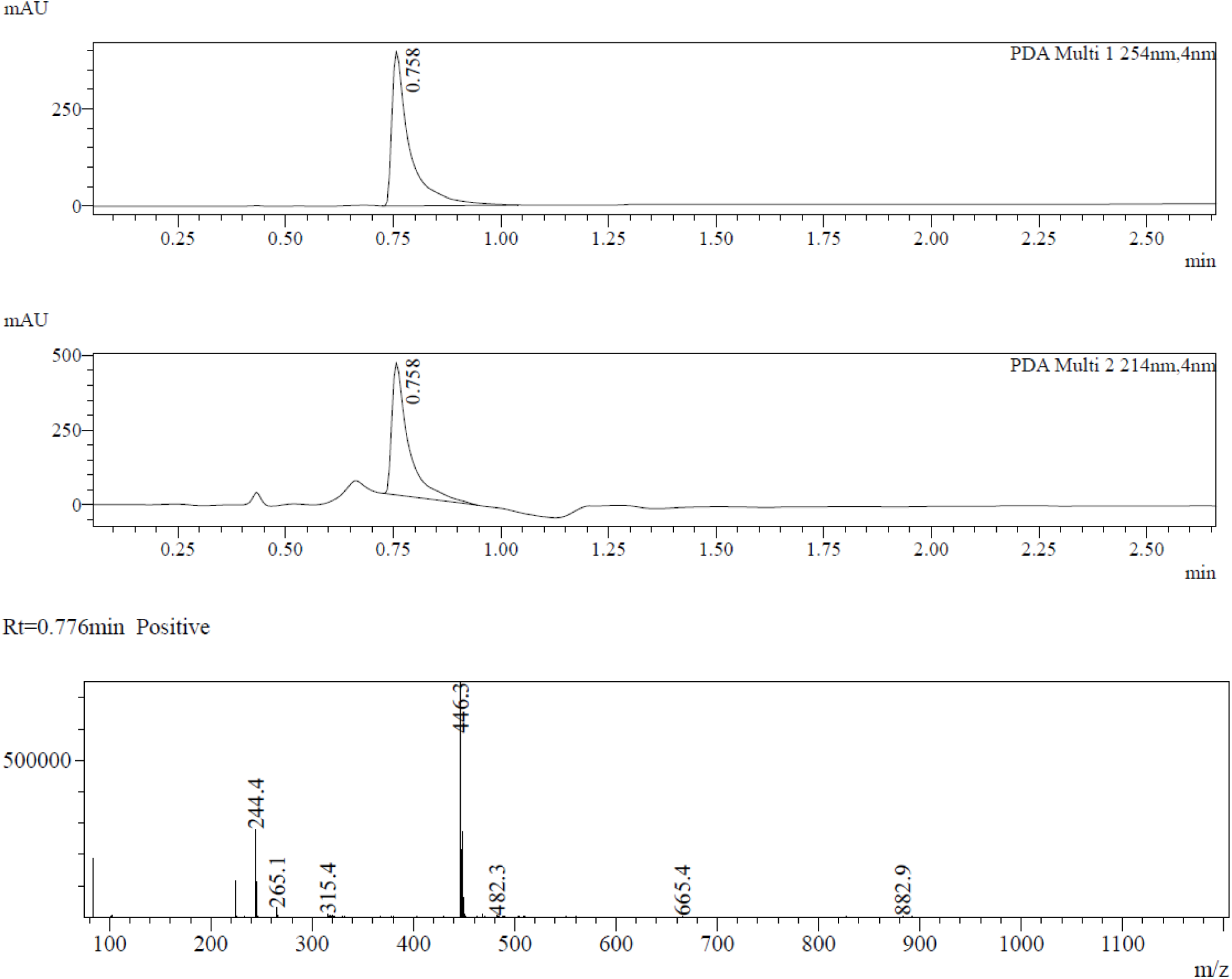

(7) Synthesis of 1-((2R,4S)-2-((bis(4-methoxyphenyl)(phenyl)methoxy)methyl)-4-hydroxypyrrolidin-1-yl)-5-(4-(3-((4-((3-chloro-4-fluorophenyl)amino)-7-methoxyquinazolin-6-yl)oxy)propyl)piperazin-1-yl)pentane-1,5-dione (B13)

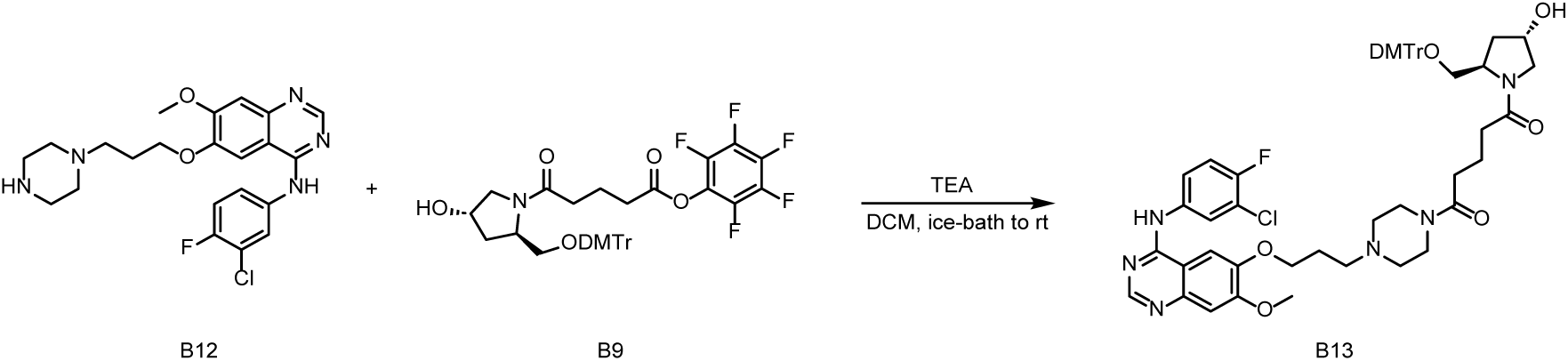

To a solution of **B12** (700 mg, TFA salt) in DCM (10 mL) was added TEA (5 mL) under nitrogen protected and stirred at 0 degree for 5 mins. **B19** (1.4 g, crude) in DCM (10 mL) was added slowly. The mixture was stirred at room temperature overnight. LCMS monitored. The reaction mixture was concentrated and purified by SGC (MeOH:DCM, 0% to 10%) to afford **B13** (670 mg, crude) as a white solid.

**LCMS:** *m/z* = 659.2 [M+H-DMTr]^+^ *t*_R_ = 11.10 min. **Purity:** 95.03 % (214 nm).

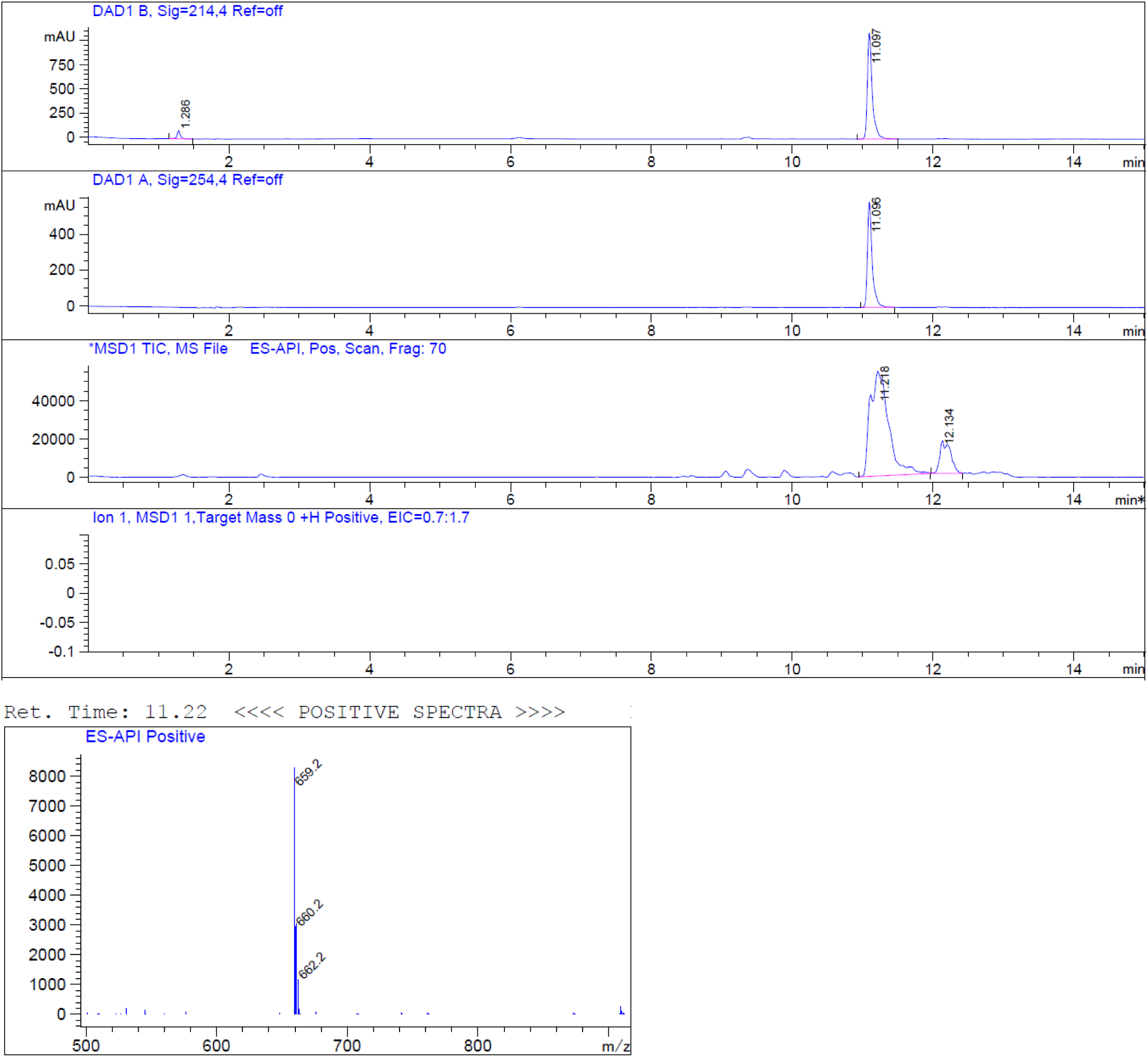

**(8) Synthesis of 4-(((3S,5R)-5-((bis(4-methoxyphenyl)(phenyl)methoxy)methyl)-1-(5-(4-(3-((4-((3-chloro-4-fluorophenyl)amino)-7-methoxyquinazolin-6-yl)oxy)propyl)piperazin-1-yl)-5- oxopentanoyl)pyrrolidin-3-yl)oxy)-4-oxobutanoic acid (B14)**

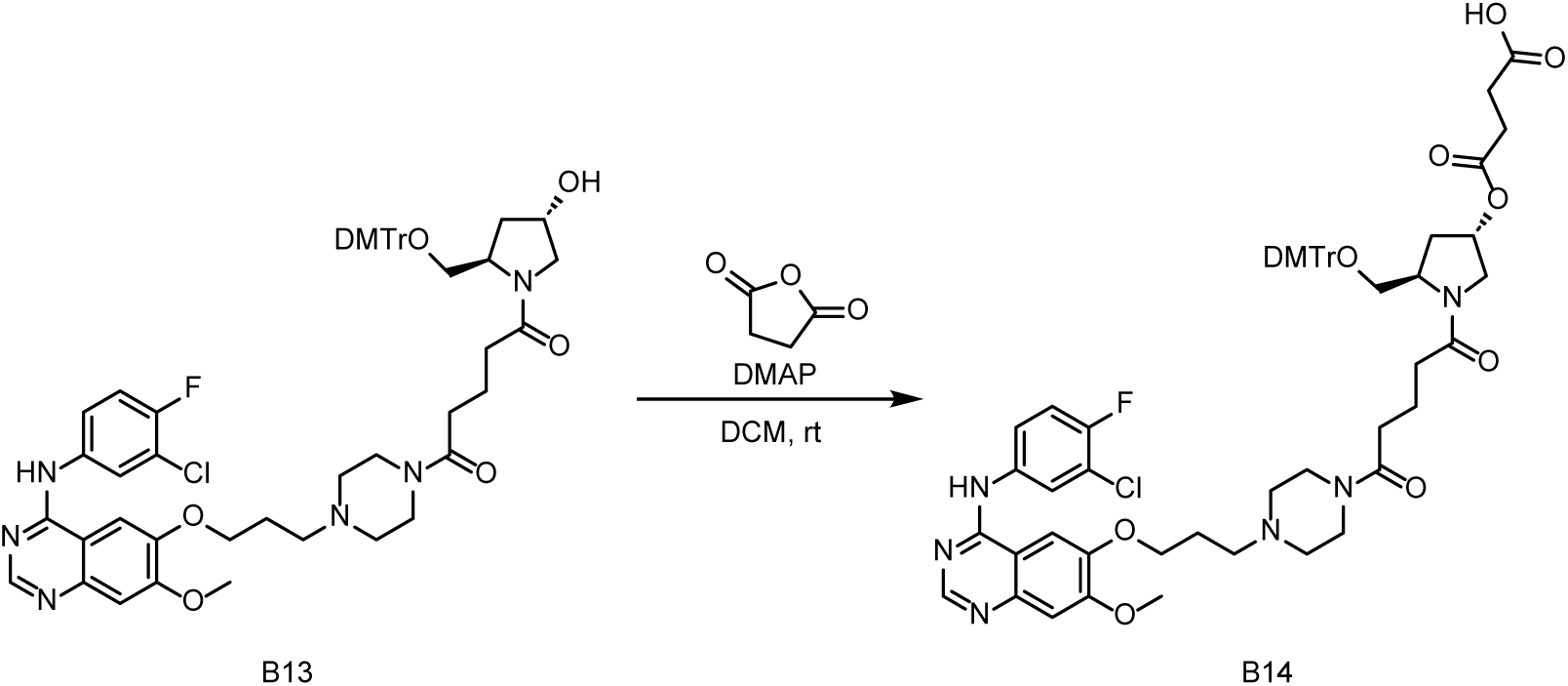

To a solution of **B13** (657 mg, crude) in DCM (20 mL) was added DMAP (205 mg, 1.68 mmol), Succinic anhydride (144 mg, 1.44 mmol). The mixture was stirred at room temperature for 8 hours. LCMS showed target MS was found. To the mixture was added TEA (0.5 mL) and stirred for 1 hour, concentrated and purified by reverse phase (**B/A**=5% ∼95%, **A**: TEA (aq, 5mL/5L), **B**: acetonitrile) to afford **B14** (349.4 mg, TEA salt) as a white solid.

**LCMS:** *m/z* = 1059.7[M-H]^-^ *t*R = 7.635 min. **Purity:** 100.00% (214 nm). **^1^HNMR** (400 MHz, DMSO-*d*_6_) *δ* 9.58 (s, 1H), 8.50 (s, 1H), 8.12 (m, 1H), 7.85 – 7.76 (m, 2H), 7.44 (m, 1H), 7.35 – 7.26 (m, 4H), 7.20 (m, 6H), 6.87 (d, *J* = 8.4 Hz, 4H), 5.30 (m, 1H), 4.19 (t, *J* = 5.6 Hz, 3H), 3.94 (s, 3H), 3.80 – 3.64 (m, 7H), 3.55 (d, *J* = 9.2 Hz, 1H), 3.40 (m, 6H), 3.24 – 3.19 (m, 1H), 2.98 (d, *J* = 6.4 Hz, 1H), 2.73 – 2.58 (m, 3H), 2.31 (m, 7H), 2.20 (m, 2H), 2.08 – 1.92 (m, 3H), 1.77 – 1.56 (m, 2H), 1.01 (t, *J* = 7.2 Hz, 4H). **^19^FNMR** (400 MHz, DMSO-*d*_6_) *δ* 123.31 (s, 1F)

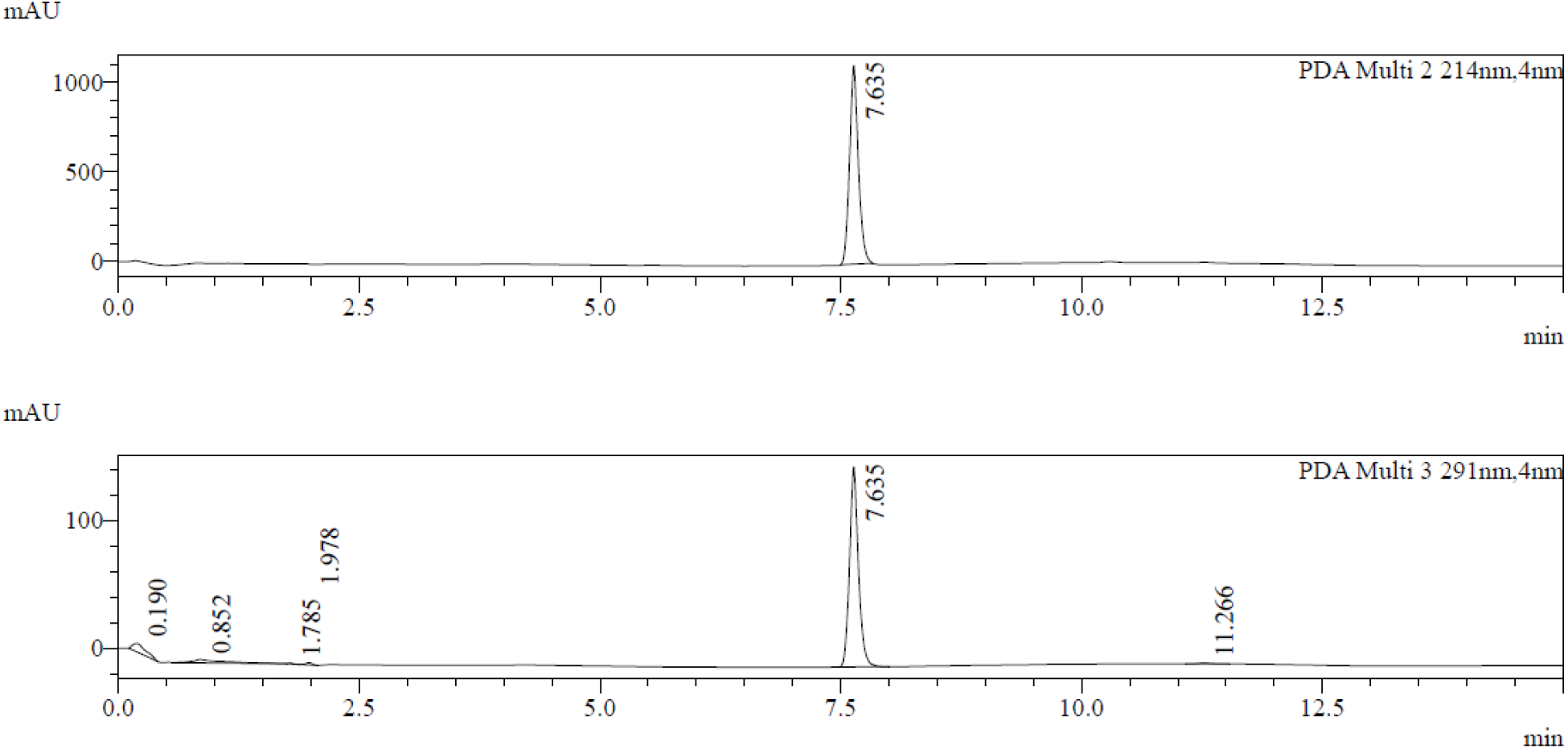
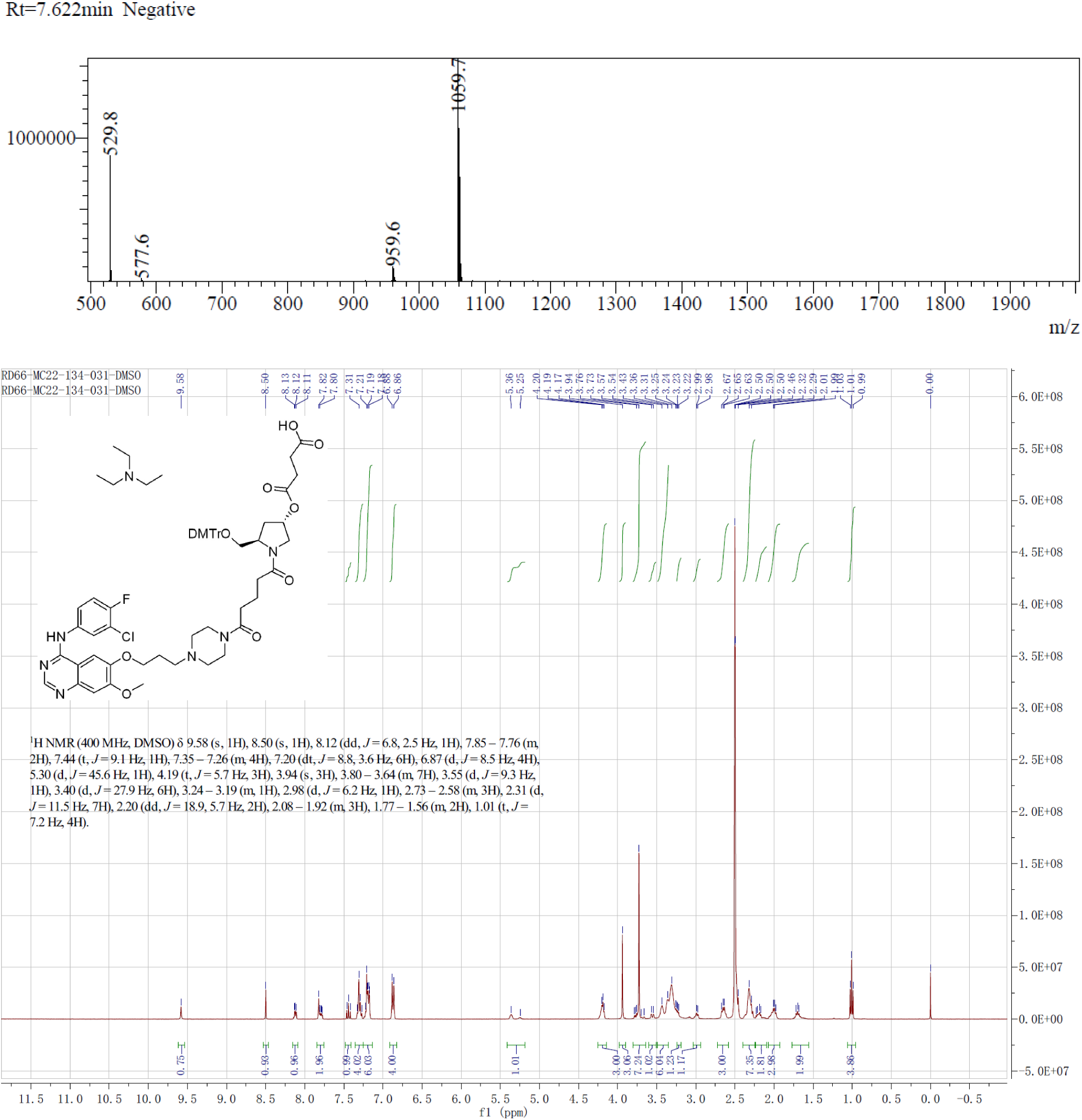
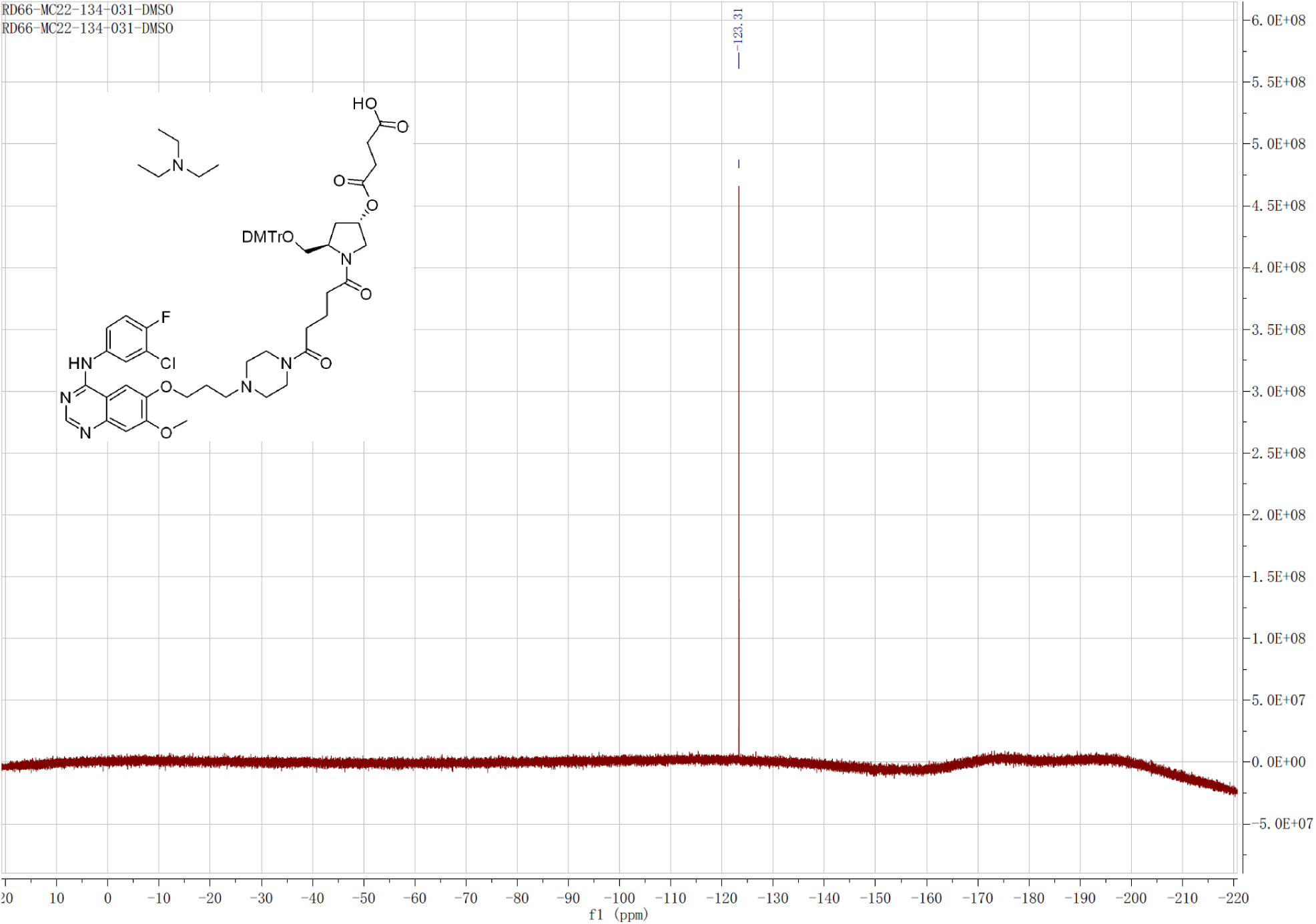

**(9) Synthesis of AS1411-Gef-CPG**

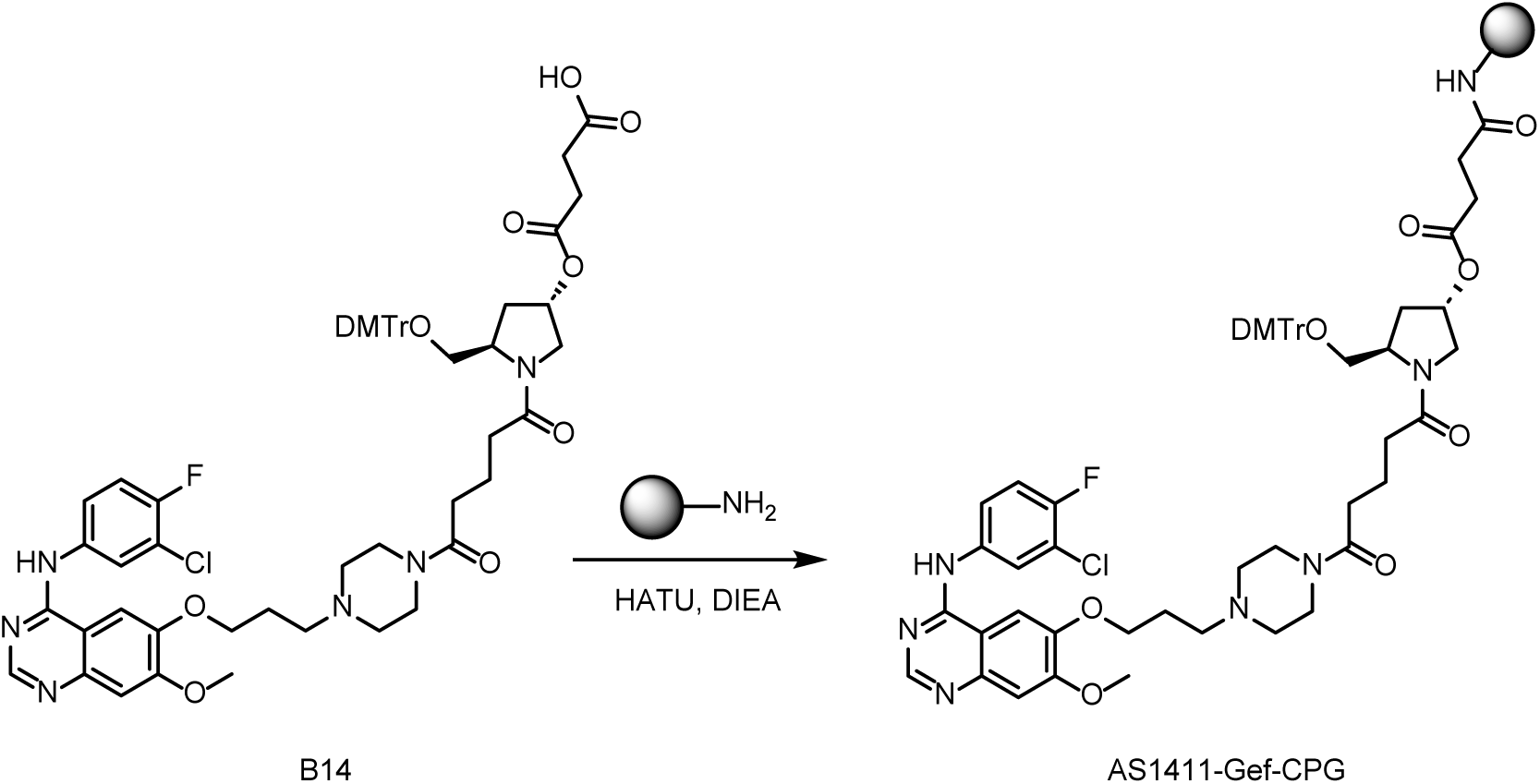

To a solution of **B14** (200 mg, 0.189 mmol) in ACN (12.0 mL) was added HATU (80 mg, 0.208 mmol), DIEA (80 μL), lcaa-CPG (1000 °A, 1000 mg) respectively at room temperature and oscillates for 12 hours. After the reaction was completed, CPG was washed with ACN, CAP A (acetic anhydride: tetrahydrofuran= 1:9, v/v, 4.0 mL) and CAP B (n-methylimidazole: pyridine: acetonitrile= 15:10:75, v/v/v, 4.0 mL) was added. The mixture was oscillated at room temperature for 1 hour. Then the mixture was filtered and washed with ACN (2 mL) for 3 times. **AS1411-Gef-CPG** was obtained after freeze-drying of the mixture as a white power (1000 mg).

**(10) Synthesis of AS1411-Gef**

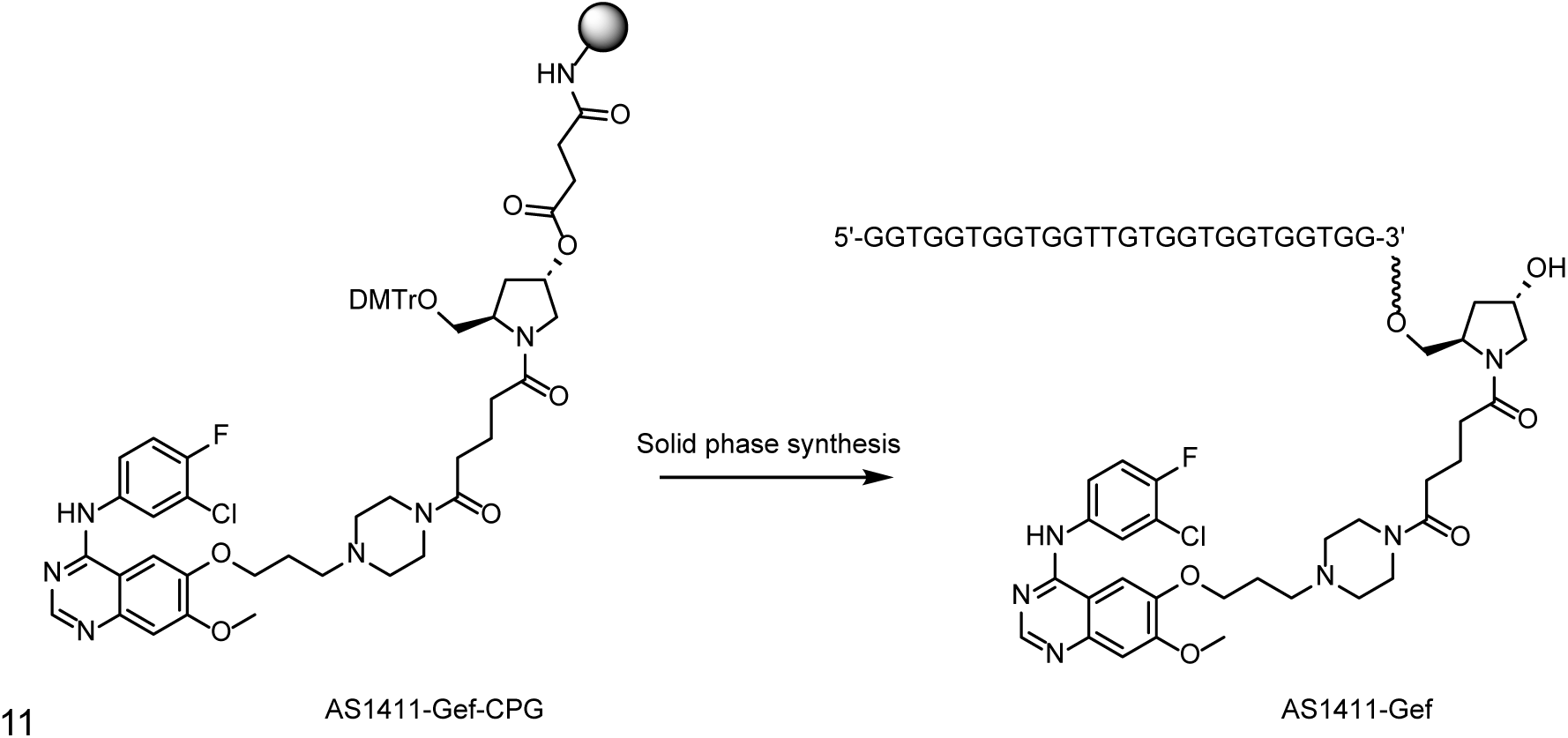

**As1411-Gef-CPG** in synthesis columns (60 mg*8) and synthesize by K-A H-8 Solid phase synthesizer. Solid phase synthesis involves four steps: Detritylation, Coupling, Capping and Oxidation. After reaction was done, CPG of each synthesis column was added 1.5 mL aqueous ammonia and heated in oven at 65 °C for 16 hours. Then collected the supernatant and washed with water (1 mL*3). The crude was purified by Protein purification system (Sepure, SDA) (**Column:** Mono Q, 1.0 mL, **method**: Mobile phase A: 40 mM NaOH in water, Mobile phase B: 40mM NaOH + 2.0 M NaCl in water) to afford **AS1411-Gef** as a white power (11.86 mg, purity = 98.18%).

**UPLC-MS (WATERS ACQUITY PREMIER):** *m/z* = 8992.79650 [M]^-^(deconvolution); *t*_R_ = 11.285 min (260 nm).

Mass error <50 ppm. **HPLC:** *t*_R_ = 11.306 min (260 nm), purity: 98.175%.

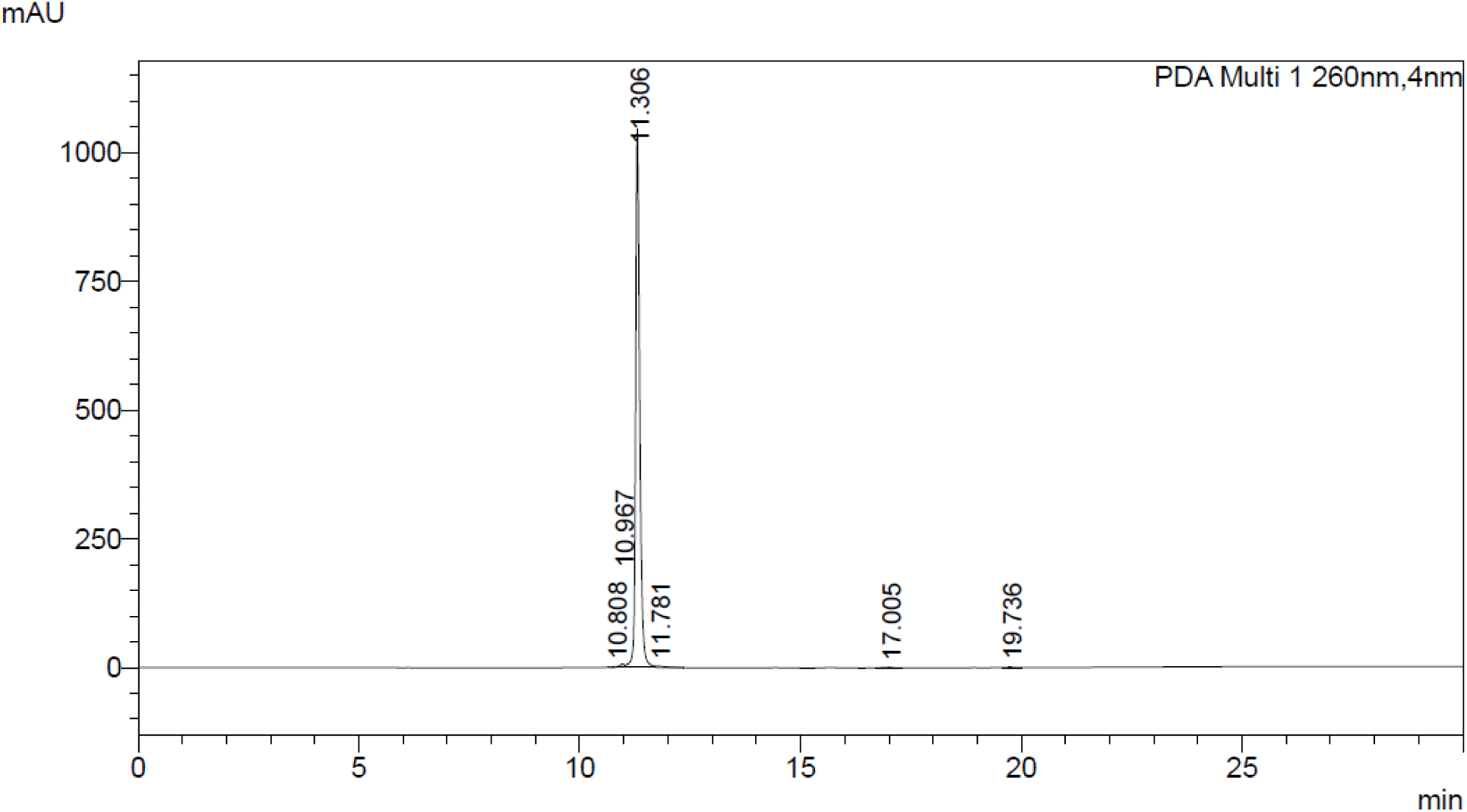
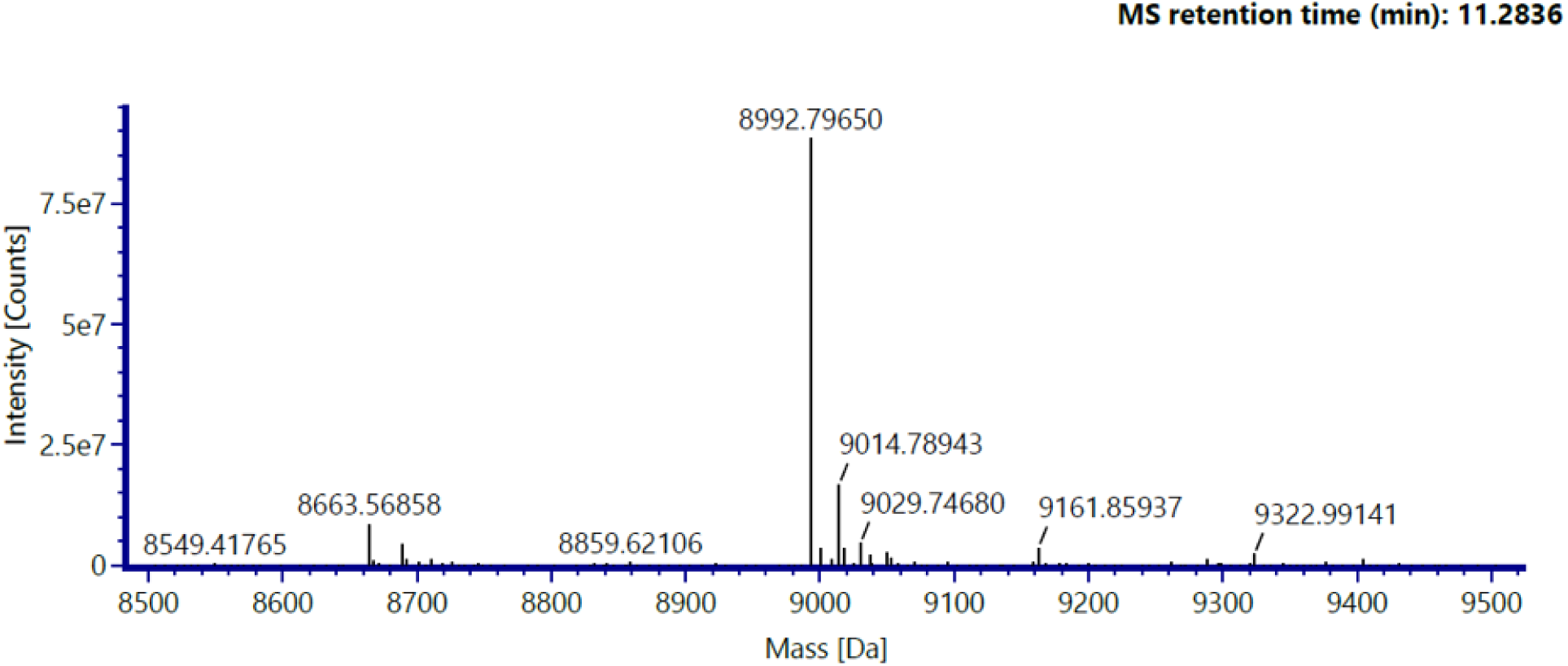

